# Covariation of amino acid substitutions in the HIV-1 envelope glycoprotein gp120 and the antisense protein ASP associated with coreceptor usage

**DOI:** 10.1101/2025.01.19.633671

**Authors:** Angelo Pavesi, Fabio Romerio

## Abstract

The tropism of the Human Immunodeficiency Virus type 1 (HIV-1) is determined by the use of either or both of the chemokine coreceptors CCR5 (R5) or CXCR4 (X4) for entry into the target cell. The ability of HIV-1 to bind R5 or X4 is determined primarily by the third variable loop (V3) of the viral envelope glycoprotein gp120. HIV-1 strains of pandemic group M contain an antisense gene termed *asp*, which overlaps *env* outside the region encoding the V3 loop. We previously showed that the ASP protein localizes on the envelope of infectious HIV-1 virions, suggesting that it may play a role in viral entry. In this study, we first developed a statistical method to predict coreceptor tropism based on the Fisher’s linear discriminant analysis. We obtained three linear discriminant functions able to predict coreceptor tropism with high accuracy (94.4%) when applied to a training dataset of V3 sequences of known tropism. Using these functions, we predicted the tropism in a dataset of HIV-1 strains containing a full-length *asp* gene. In the amino acid sequence of ASP proteins expressed from these *asp* genes we identified five positions with substitutions significantly associated with viral tropism. Interestingly, we found that these substitutions correlate significantly with substitutions at six amino acid positions of the V3 loop domain associated with tropism. Altogether, our computational analyses identify ASP amino acid signatures coevolving with V3 and potentially affecting HIV-1 tropism, which can be validated through *in vitro* and *in vivo* experiments.

## Introduction

The entry of the human immunodeficiency virus 1 (HIV-1) into human host cells is initiated by interaction of the viral envelope glycoprotein gp120 with the cellular CD4 receptor [1]. This primary interaction induces a conformational change in the gp120 ectodomain of the protein [2] that enables viral binding to one of the cell surface co-receptors, CCR5 or CXCR4 [3,4]. This sequence of events results in membrane fusion and penetration of the virus into the host cell [5]. The third hypervariable loop domain (V3) of gp120, a sequence of ~35 amino acids, is the major determinant of coreceptor tropism [6]. Viral strains are classified as R5-tropic when using the CCR5 coreceptor, X4-tropic when using CXCR4, and R5X4-tropic or dual tropic when using both coreceptors. While R5-tropic viruses are present at all stages of infection, progression towards AIDS disease is often associated with the emergence of X4-tropic strains [7].

Two types of methods have been developed for testing viral tropism: *i)* cell-based *in vitro* phenotypic tests such as ES-Trofile [8]; *ii) in silico* genotypic tests, based on sequencing the V3 loop domain of gp120 and using bioinformatics methods to predict coreceptor usage. Although phenotypic tests have a high sensitivity in distinguishing R5-, R5X4-, and X4-tropic viral strains, they are expensive and time consuming. As an alternative, genotypic tests are easily accessible, fast, and inexpensive.

The first genotypic method to predict coreceptor usage is the 11/25 charge rule, which classifies the virus as X4-tropic if a positively charged amino acid is observed in positions 11 or 25 of the V3 sequence [9]. As reported by Lengauer et al. [10], this simple rule shows a good specificity (93% of the non-X4-tropic strains predicted as non-X4-tropic) but a low sensitivity (only 60% of the X4-tropic strains predicted as X4-tropic), making it unsuitable for routine clinical use. Other methods use more complex statistical models, such as neural networks [11], support vector machines [12,13], position-specific scoring matrices [14], incorporation of physicochemical and structural properties of the V3 sequence into a numerical descriptor [15], coreceptor-specific weight matrices [16], and machine learning/hidden Markov model [17]. The method Geno2pheno [10] combines two machine learning approaches, support vector machine and decision tree, and uses clinical information such as viral loads and CD4 cell counts if available. Currently, Geno2pheno is the only genotypic method recommended for use in clinical routines by the European Consensus Group [18].

Since establishing HIV-1 tropism is crucial to address antiviral treatment, significant efforts have been made to improve the performance of genotypic tests in terms of sensitivity and specificity. Overall, the performance is high when predicting R5-tropic sequences but relatively low for X4-tropic sequences, likely because most of the V3 sequences from viral samples are in the R5-tropic group.

With the exception of the multiple linear regression [19], multivariate statistical methods have been overlooked as a method for the prediction of viral tropism. In the present study, we developed a genotypic test based on the Fisher’s linear discriminant analysis [20,21] to predict coreceptor tropism in a dataset of HIV-1 strains that contain the antisense gene *asp*, which overlaps the gp120 gene on the frame −2 and encodes the antisense protein ASP [22–25]. Using this test, we identified amino acid changes in ASP that are preferentially associated with R5 or X4 usage. Interestingly, we found that these substitutions in ASP correlate significantly with substitutions within the principal determinants of coreceptor tropism, the variable loops V3 and, to a lesser extent, V1/V2 of gp120. Altogether, our study identifies ASP residues coevolving with V3 and potentially affecting HIV-1 tropism, which can be validated through *in vitro* and *in vivo* experiments.

## 2. Materials and Methods

### 2.1 Assembly of a training dataset of V3 sequences with known tropism

We downloaded the dataset of V3 sequences developed by Shen et al. [16]. It contains 2679 V3 sequences from HIV-1 strains with a tropism determined by a variety of phenotypic tests, as reported in the Los Alamos HIV sequence database (www.hiv.lanl.gov/content/index). By exploring the Shen’s dataset to select V3 sequences from strains of genotype B or C, we assembled a training dataset of 1701 sequences R5-tropic (1229 of genotype B and 472 of genotype C) and 137 sequences purely X4-tropic (96 of genotype B and 41 of genotype C). Purely X4-tropic means that we did not include in the training dataset V3 sequences from viral strains that use both coreceptors.

### 2.2 Linear discriminant analysis (LDA) of the training dataset

To convert each V3 sequence of the training dataset into a numerical format, we extracted from the AAindex database [26] the subset of 46 indices that quantitatively evaluate the physicochemical properties of amino acids. Within the subset of 149 hydrophobicity indices, we selected the most widely-used hydropathy index by Kyte and Doolittle [27]. The numerical descriptor of each V3 sequence was a vector of 7 components, the first is the arithmetic mean of the hydropathy index and the others are the arithmetic mean of 6 indices of physicochemical properties randomly taken from the 46 ones. Thus, the input data for LDA were a matrix of V3 sequences R5-tropic (1701 rows and 7 columns) and a matrix of V3 sequences X4-tropic (137 rows and 7 columns). LDA yielded a linear function with 7 coefficients and the corresponding degree of accuracy. Accuracy was the number of R5-tropic sequences predicted correctly plus the number of X4-tropic sequences predicted correctly, divided by the total number of sequences, and multiplied by 100. To find the combination(s) of 7 amino acid indices with linear function(s) having the highest accuracy, we carried out 100,000 LDAs and selected the cases with an accuracy of ~94.0%.

To evaluate the robustness of LDAs showing the best accuracy, we carried out two cross-validation tests. The first depends on the fact that the training dataset contains a number of sequences R5-tropic more than 10-fold larger than that of sequences X4-tropic (1701 vs. 137). For each combination of amino acid indices, the validation test consisted of *i)* 1000 LDAs, each of them between the 137 X4-tropic sequences and a subset of R5-tropic sequences of equal size randomly taken from the 1701 ones; *ii)* calculation of the mean accuracy in the prediction of R5 or X4 usage.

For each combination of amino acid indices, the other cross-validation test consisted of *i)* creation of a training subset composed of 65 R5-tropic sequences and 65 X4-tropic sequences, randomly taken from the 1701 and 137 ones, respectively; *ii)* calculation of the linear discriminant function; *iii)* evaluation of its accuracy on a validation subset composed of 65 R5-tropic sequences and 65 X4-tropic sequences, randomly taken from the 1701 and 137 ones respectively, and different from those of the training subset. We repeated this procedure 1000 times and calculated the mean accuracy in the prediction of R5 or X4 usage.

### 2.3 Prediction of the tropism in a dataset of viral strains of genotype B or C containing an antisense gene asp

Selection of LDAs with an accuracy ~94.0%, followed by evaluation of their robustness by cross-validation testing, yielded 3 combinations of 7 amino acid indices in which LDA showed the highest prediction accuracy (see Results). Using the corresponding 3 linear functions, we predicted the tropism in a dataset of 2593 *env* sequences (1625 of genotype B and 968 of genotype C) taken from our previous study on the antisense gene *asp* [22]. These *env* sequences, aligned for a total of 1274 codon positions, contain an antisense gene termed *asp* [24], which overlaps the *env* gene at the surface and transmembrane boundary and outside the *env* region encoding the V3 loop domain [25,26]. We extracted from each aligned *env* sequence the V3 region (from codon position 479 to 531) and after translation into amino acids predicted its tropism. We classified a given V3 sequence as R5-tropic (or X4-tropic) only if predicted as R5-tropic (or X4-tropic) by all 3 linear discriminant functions. By this rule, we assigned the tropism to 2496 out of 2593 V3 sequences (see Results).

### 2.4 Comparative analysis of the amino acid composition in the V3 region between R5-tropic and X4-tropic viral strains

Once predicted the tropism, we compared the amino acid content in the V3 region of strains R5-tropic with that of strains X4-tropic, with the aim to detect amino acid positions significantly associated with coreceptor usage. We analyzed the dataset of 2496 V3 regions, aligned for a total of 53 amino acid positions, and selected for statistical analysis 35 positions in which the frequency of gaps was <5%. At each position, we compared the amino acid content in sequences R5-tropic with that in sequences X4-tropic using the χ^2^ contingency-table test. We performed the same analysis on the V1/V2 region (from codon position 189 to 350 in the full-length aligned *env* sequences), on the *env* region upstream V1/V2 and not overlapping the *vpu* gene (from codon position 51 to 188), on the *env* region between V1/V2 and V3 (from codon position 351 to 478), and on the *env* region downstream V3 and not overlapping the antisense *asp* gene (from codon position 532 to 605).

### 2.5 Comparative analysis of the amino acid composition in the env/asp overlap between R5-tropic and X4-tropic viral strains

The region of *env* overlapping *asp* is an antisense −2 coding region, in which a codon of *asp* overlaps with two contiguous codons of *env* on the opposite strand. In detail, third base of *asp* overlaps with third base at codon *n* of *env*, while second and first base of *asp* overlap respectively with first and second base at codon (*n +1*) of *env*. We extracted from each of the 2496 aligned *env* sequences the region that overlaps with *asp* (from codon position 606 to 894) and selected for statistical analysis 170 codon positions in which the frequency of gaps was <5%. At each position, we compared the amino acid content in sequences R5-tropic with that in sequences X4-tropic using the χ^2^ contingency-table test. We performed the same analysis on each codon position of *asp* that overlaps with two contiguous codons of *env* on the opposite strand.

### 2.6 Identification of significant patterns of pairwise associations between amino acid substitutions in ASP and amino acid substitutions in V3 loop region

We detected, both in ASP and V3, a number of positions in which the amino content in strains R5-tropic differs significantly from that in strains X4-tropic (see Results). To test if there is a significant association between V3 and ASP substitutions, we calculated for each pair of amino acid changes the φ binomial correlation coefficient. It is similar to Pearson’s correlation coefficient but is specifically used for categorical data arranged in a 2 χ 2 contingency table. A φ value comprised between 0.3 and 0.7 indicates a weak-medium pairwise association, while a φ value comprised between 0.7 and 1.0 indicates a strong pairwise association.

## 3. Results

### 3.1 Linear discriminant analysis (LDA) accurately predicts HIV-1 coreceptor usage

We carried out LDA on a training dataset of 1701 R5-tropic V3 sequences (1229 of genotype B and 472 of genotype C) and 137 X4-tropic V3 sequences (96 of genotype B and 41 of genotype C). Sequence data are reported in Supplementary File S1. The mean length of V3 sequences was 34.8 amino acids, with a very low standard deviation (0.6). We converted each V3 sequence in numerical form using the Kyte-Doolittle hydropathy index [27] and 6 indices of physicochemical properties of amino acids, taken at random from the subset of 46 indices in AAindex database [26].

We randomly generated 100,000 combinations of 7 amino acid indices and found that in most of them (86,892) LDA predicts the tropism with an accuracy from 87.3 to 94.7%. We found, however, that the accuracy is biased toward a more accurate prediction for R5 than X4 usage. Indeed, we found a very small number of combinations (11 cases) in which LDA predicts the tropism with an accuracy ~94% both for R5 and X4 usage. We evaluated the robustness of these LDAs by two cross-validation tests (see Methods) and found the best performance for 3 combinations of amino acid indices. Indeed, sensitivity, specificity, and accuracy of LDA#1, LDA#2, and LDA#3 are all ~94% (Table 1). The value of MCC (Matthew’s correlation coefficient) [28] is moderate (0.71), where value ‘1’ corresponds to the perfect prediction and ‘0’ to a completely random prediction. The moderate value of MCC is due to the robustness of this parameter when dealing with different class sizes, just like in this case where the number of sequences R5-tropic is one order of magnitude greater than that of sequences X4-tropic.

**Table 1.** Performance of LDA on the training datasets of V3 sequences from 1701 R5-tropic and 137 X4-tropic HIV-1 isolates.

| LDA | Sensitivity<sup>1</sup> | Specificity<sup>2</sup> | Accuracy<sup>3</sup> | MCC<sup>4</sup> | Mean sensitivity by first cross-validation test | Mean specificity by first cross-validation test | Mean accuracy by first cross-validation test | Mean sensitivity by second cross-validation test | Mean specificity by second cross-validation test | Mean accuracy by second cross-validation test |
| --- | --- | --- | --- | --- | --- | --- | --- | --- | --- | --- |
| LDA#1 | 94.2 | 94.5 | 94.5 | 0.71 | 93.7 | 94.5 | 94.1 | 92.3 | 93.0 | 92.6 |
| LDA#2 | 94.2 | 94.2 | 94.2 | 0.71 | 93.8 | 94.3 | 94.0 | 92.8 | 92.7 | 92.7 |
| LDA#3 | 94.2 | 94.7 | 94.6 | 0.72 | 93.3 | 94.3 | 93.8 | 92.3 | 93.1 | 92.7 |

<sup>1</sup>Sensitivity is $(TP / (TP + FN)) \times 100$ , where TP is the number of true positives (number of correctly predicted X4-tropic sequences) and FN is the number of false negatives (X4-tropic sequences predicted as R5-tropic) <sup>2</sup>Specificity is $(TN / (FP + TN)) \times 100$ , where TN is the number of true negatives (number of correctly predicted non-X4-tropic sequences) and FP is the number of false positives (R5-tropic sequences predicted as X4-tropic) <sup>3</sup>Accuracy is $[(TP + TN) / (TP + FP + TN + FN)] \times 100$ . <sup>4</sup>MCC, the Matthew's correlation coefficient [27], is:

In addition to the fixed index of hydropathy (AAindex entry KYTJ8201201), the 3 combinations of 7 amino acid indices have 4 physicochemical properties in common: residue volume (entry GOLD730102 for LDA#1 and entry BIGC670101 for LDA#2 and LDA#3), relative mutability (entry DAYM780201 for LDA#1 and LDA#3, and entry JOND920102 for LDA#2), entropy of formation (HUTJ700103), and side chain volume (KRIW790103). Unique to LDA#1 were indices OOBM850102 (Optimized propensity to form reverse turn) and ANDN9290101 (Alpha-CH chemical shifts), to LDA#2 indices GARJ730101 (Partition coefficient) and FAUJ880104 (Length of the side chain), and to LDA#3 indices CHAM830105 (Number of atoms in the side chain) and OOBM770102 (Short and medium range non-bonded energy per atom).

The high discriminant power of LDA#1, LDA#2, and LDA#3 can be appreciated by examining the distribution of the corresponding LDA scores in V3 sequences R5-tropic and in V3 sequences X4-tropic (Figure 1A, B, and C). A detailed description of the method (LDA#1, LDA#2, and LDA#3) is reported in Supplementary File S2.

**Figure 1.**
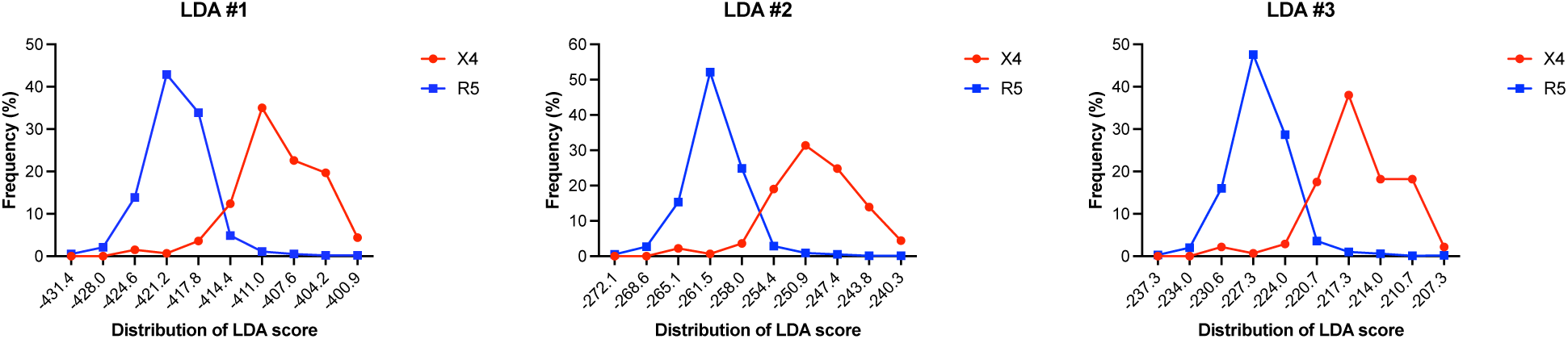
**A)** Percent frequency distribution of the LDA#1 score in 1701 V3 sequences R5-tropic and in 137 V3 sequences X4-tropic. With a cut-off score of −417.38, 1608 out of 1701 sequences R5-tropic (94.5%) were predicted as R5 (score below the cut-off), and 129 out of 137 sequences X4-tropic (94.2%) as X4 (score above the cut-off). **B)** Percent frequency distribution of the LDA#2 score in the same dataset. With a cut-off score of −258.33, 1603 out of 1701 sequences R5-tropic (94.2%) were predicted as R5 (score below the cut-off), and 129 out of 137 sequences X4-tropic (94.2%) as X4 (score above the cut-off). **C)** Percent frequency distribution of the LDA#3 score in the same dataset. With a cut-off score of −224.01, 1610 out of 1701 sequences R5-tropic (94.6%) were predicted as R5 (score below the cut-off), and 129 out of 137 sequences X4-tropic (94.2%) as X4 (score above the cut-off).

### 3.2 Prediction of tropism in a dataset of V3 sequences and detection of amino acid positions associated with coreceptor usage

Using the 3 linear functions yielded respectively by LDA#1, LDA#2, and LDA#3, we predicted the tropism in a dataset of 2593 aligned *env* sequences (1625 of genotype B and 968 of genotype C) taken from our previous study [22]. We extracted from each *env* sequence the V3 region and predicted the tropism based the following rule: a V3 region was classified as R5-tropic (or X4-tropic) only if predicted as such by all 3 linear functions. By this stringent approach, we assigned the tropism to 2496 out of 2593 V3 regions (96.2%): 2261 were predicted as R5-tropic and 235 as X4-tropic.

At each position of V3, we compared the amino acid content in sequences R5-tropic with that in sequences X4-tropic using the χ^2^ contingency-table test. Over a total of 35 amino acid positions suitable for statistical analysis (frequency of gaps <5%), we found 20 showing a significant difference between R5-tropic and X4-tropic. To highlight the substitutions most closely related to the tropism, we considered only amino acid changes with prevalence ≥5% in sequences R5- or X4-tropic, respectively. We found that the 20 amino acid positions account for 42 different substitutions, 34 of them showing a significant prevalence in strains of the X4-tropic group (Figure 2).

**Figure 2.**
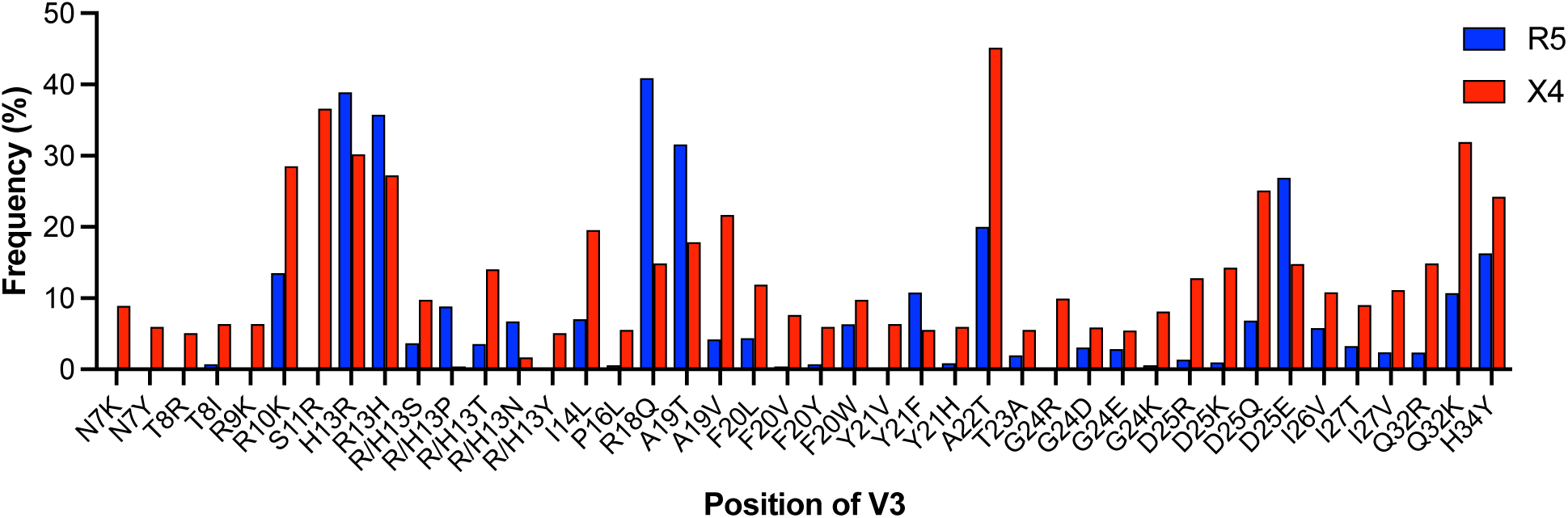
Percent frequency of the 42 amino acid substitutions in the V3 loop significantly associated to coreceptor tropism. 34 of them show a statistically significant prevalence in strains of the X4-tropic group (red column).

### 3.3 Detection of amino acid positions in the ASP protein associated with coreceptor usage

We used the dataset of 2496 *env* sequences with predicted tropism to test the presence in ASP of amino acid sites associated with R5 or X4 usage. We extracted from each aligned sequence the region of *env* that overlaps with *asp* (289 codon positions) and selected 170 positions suitable for statistical analysis (frequency of gaps <5%). The *env*/*asp* is an antisense −2 overlapping region, in which two contiguous codons of *env* overlap with a single codon of *asp*. By considering 170−1 pairs of contiguous codons in *env*, we obtained the same number of codon positions in the *asp* antisense frame.

At each codon position of *asp*, we compared the amino acid content in sequences R5-tropic with that in sequences X4-tropic using the χ^2^ contingency-table test. We repeated the analysis on each codon position of *env*. This analysis was crucial to detect the amino acid positions of ASP in which the presence of a significant difference in composition between R5 and X4 usage is paired, instead, to the lack of a significant difference between R5 and X4 usage in the two contiguous antisense positions of *env*.

Consider for example the amino acid position 106 of ASP, in which we found a significant difference in composition (P = 0.00007) between R5 and X4 usage (39% content in Asn for R5 and 22% for X4). In contrast, the two contiguous amino acid positions of ENV overlapping position 106 of ASP did not show any significance difference in composition between R5 and X4 usage (P = 0.40 and P = 0.86 respectively). This result indicates that the amino acid diversity at position 106 is exclusive to ASP and not imposed by amino acid substitutions in ENV.

Based on this rule and considering only substitutions with prevalence ≥5% in sequences R5- or X4-tropic respectively, we found five amino acid positions in ASP with substitutions associated preferentially with coreceptor usage (Figure 3): one is located in the N-terminal intracellular domain (position 20), two in the ectodomain (positions 106 and 119), and two in the C-terminal transmembrane domain (positions 157 and 161). At positions 20 and 157, the amino acid substitutions show a significant prevalence in strains X4-tropic, while at positions 106, 119, and 161 they significantly prevail in strains R5-tropic (Figure 3).

**Figure 3.**
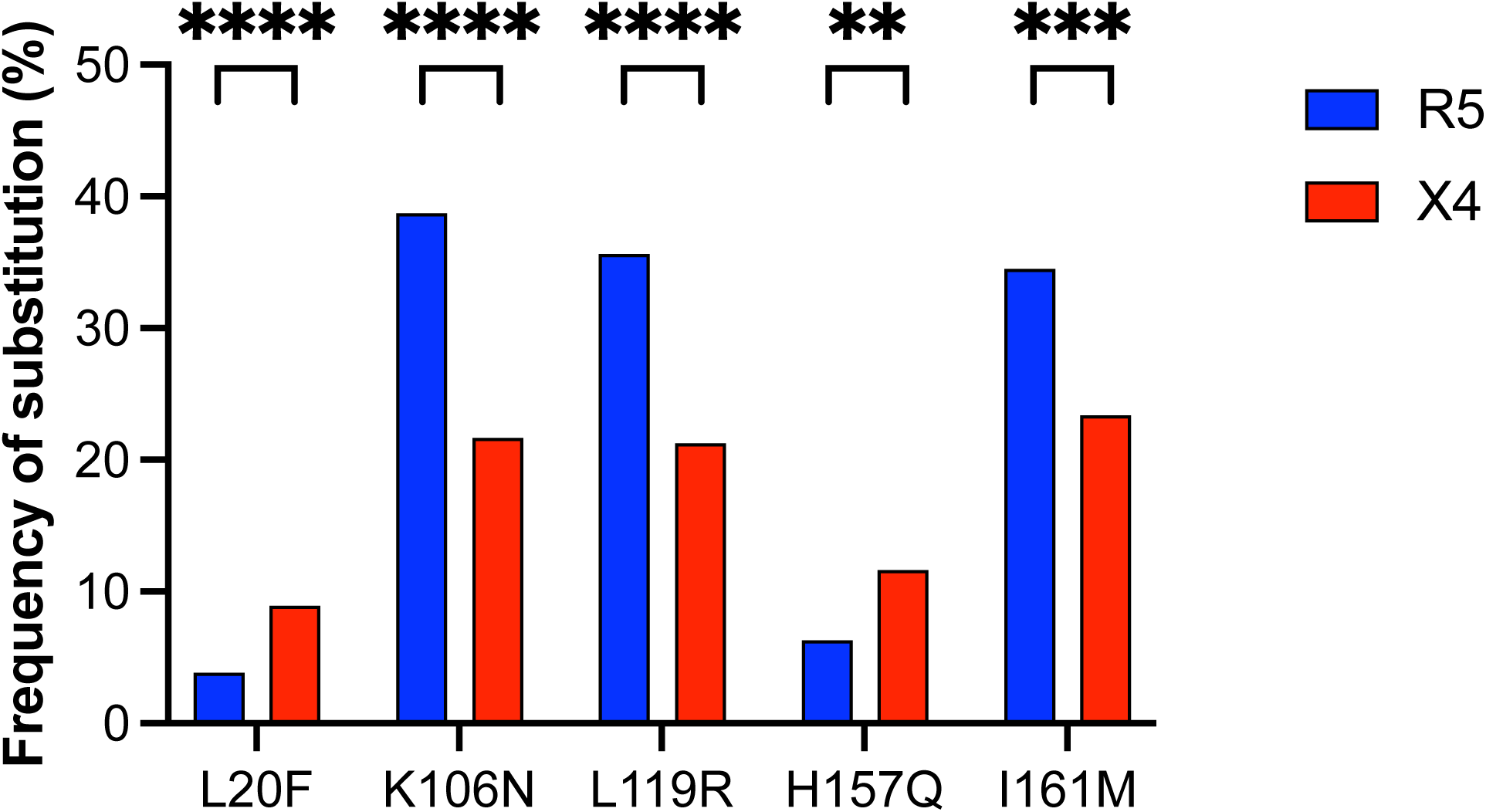
Percent frequency of the 5 amino acid substitutions in the ASP protein significantly associated to coreceptor tropism.

### 3.4 Detection of significant patterns of association between V3 and ASP amino acid substitutions

Next, we tested if there is a significant association between the 42 amino acid substitutions in V3 and the 5 amino acid substitutions in ASP associated with coreceptor usage (Figure 2 and 3). For each pair of substitutions, we calculated the φ correlation coefficient and its statistical significance. A positive and significant correlation (φ > 0.3) indicates that the co-occurrence of two amino acid substitutions is not due to chance and may confer advantage in terms of coreceptor usage.

We found that the highest values of the φ correlation coefficient involve substitutions located in the ectodomain of ASP (Table 2). We found that substitutions K106N and L119R of ASP are both strongly correlated (φ from 0.83 to 0.87) with substitutions H13R and R18Q of V3. This pattern of covariation occurs in ~90% of strains of genotype C (Table 2), whereas it is extremely rare in strains of genotype B. Substitution I161M of ASP is moderately correlated (φ = 0.56) with substitutions H13R and R18Q of V3. Again, covariation is specific to genotype C, yet in this case it occurs in ~67% of strains of genotype C (Table 2). All 3 substitutions in ASP (K106N, L119R, and I161M) are significantly correlated (φ from 0.37 to 0.56) with substitution A19T of V3. In this case, covariation between I161M and A19T (φ = 0.37) occurs in an even smaller fraction (47%) of strains of genotype C (Table 2).

**Table 2.** Amino acid substitutions in the ASP protein significantly associated with amino acid substitutions in the V3 loop of ENV.

| ASP substitution | Location in ASP | V3 substitution | Prevalent tropism | $\phi$ binomial correlation coefficient<sup>1</sup> | Subtype prevalence (%) |
| --- | --- | --- | --- | --- | --- |
| K106N | Ectodomain | H13R | R5 | 0.85 (***) | C (93%) |
|  |  | R18Q | R5 | 0.87 (***) | C (90%) |
|  |  | A19T | R5 | 0.56 (**) | C (63%) |
| L119R | Ectodomain | H13R | R5 | 0.83 (***) | C (86%) |
|  |  | R18Q | R5 | 0.85 (***) | C (87%) |
|  |  | A19T | R5 | 0.56 (**) | C (60%) |
| I161M | C-terminal<br>transmembrane domain | H13R | R5 | 0.56 (**) | C (67%) |
|  |  | R18Q | R5 | 0.55 (**) | C (68%) |
|  |  | A19T | R5 | 0.37 (*) | C (47%) |
| L20F | N-terminal<br>intracellular domain | I14L | X4 | 0.45 (*) | B (4.4%) |
|  |  | F20W | X4 | 0.67 (**) | B (5.7%) |
|  |  | D/E25Q | X4 | 0.37 (*) | B (3.9%) |
| H157Q | C-terminal<br>transmembrane domain | I14L | X4 | 0.34 (*) | B (5.6%) |
|  |  | F20W | X4 | 0.51 (**) | B (5.8%) |

<sup>1</sup>A $\phi$ value from 0.3 to 0.7 indicates a weak or medium association (\*; \*\*), a $\phi$ value from 0.7 to 1.0 indicates a strong association (\*\*\*)

As shown at the bottom of Table 2, we found a weak/moderate correlation (φ from 0.34 to 0.67) between substitutions L20F and H157Q of ASP and substitutions I14L and F20W of V3. This pattern of covariation is virtually absent in genotype C and it occurs only in a small fraction (~5%) of strains of genotype B (Table 2). Covariation between substitution L20F of ASP and substitution D/E25Q of V3 (φ = 0.37) occurs in an even smaller fraction (3.9%) of strains of genotype B (Table 2).

### 3.5 Detection of amino acid positions in the V1/V2 region associated with coreceptor usage and detection of significant patterns of association between V1/V2 and ASP substitutions

It has long been known that the V1/V2 region of ENV, although to a lesser extent than V3, can determine the selectivity of HIV-1 strains for CCR5 or CXCR4 [29–31]. Therefore, we carried out a comparative analysis of strains R5- and X4-tropic for the amino acid composition in the V1/V2 region, with the aim to detect positions associated with coreceptor usage.

We extracted from each of the 2496 *env* sequences with predicted tropism the V1/V2 region (from codon position 189 to 350). After translation into amino acids, we obtained 2496 V1/V2 regions, aligned for a total of 162 positions. We selected for statistical analysis 65 positions in which the frequency of gaps was <5% and compared their amino acid content in sequences R5-tropic with that in sequences X4-tropic using the χ^2^ contingency-table test. We found 10 positions showing a significant difference between R5 and X4 usage. They account for 13 different amino acid substitutions, 10 of them significantly prevailing in strains X4-tropic (Figure 4). We then tested if there is a significant covariation between the 13 substitutions in V1/V2 and the 5 substitutions in ASP associated with coreceptor usage (Figure 3).

**Figure 4.**
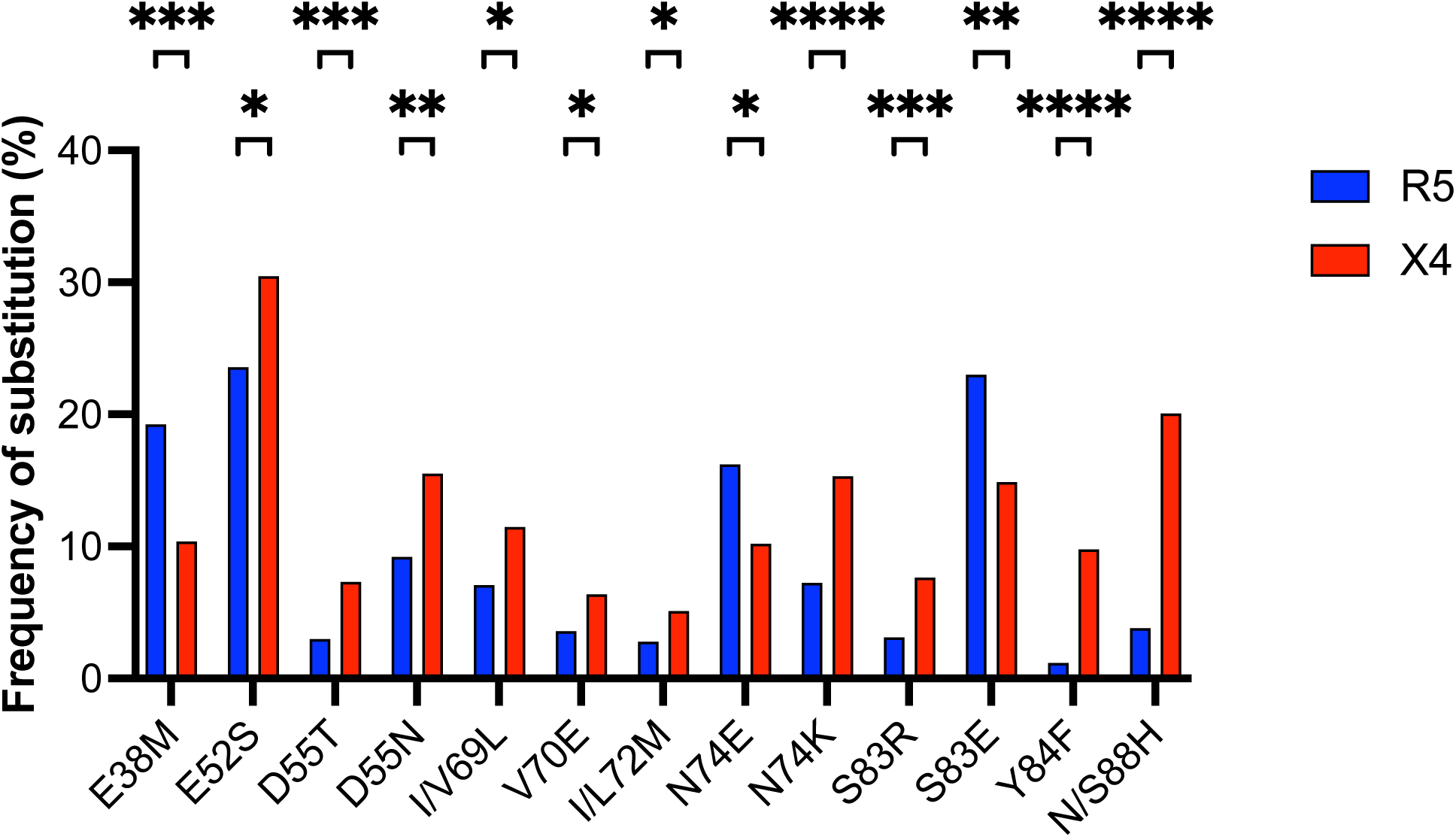
Percent frequency of the 13 amino acid substitutions in the V1/V2 loop significantly associated to coreceptor tropism. 10 of them show a statistically significant prevalence in strains of the X4-tropic group (red column).

As shown in Table 3, we found that substitutions K106N and L119R of ASP are strongly correlated (φ = 0.78) with substitution I/L72M and moderately correlated (φ = 0.60 and φ = 0.57) with substitution S83E of V1/V2. A weaker correlation was observed between substitutions I161M of ASP and I/L72M (φ = 0.51) and S83E (φ = 0.36) of V1/V2. This pattern of covariation is unique to genotype C, where it occurs with a frequency from 39 to 82% (Table 3). In addition, we found a strong/moderate correlation between substitutions L20F and H157Q of ASP and substitution D55T of V1/V2 (φ = 0.88 and φ = 0.68). This pattern of covariation occurs only in a small fraction (~5%) of strains of genotype B (Table 3).

**Table 3.**
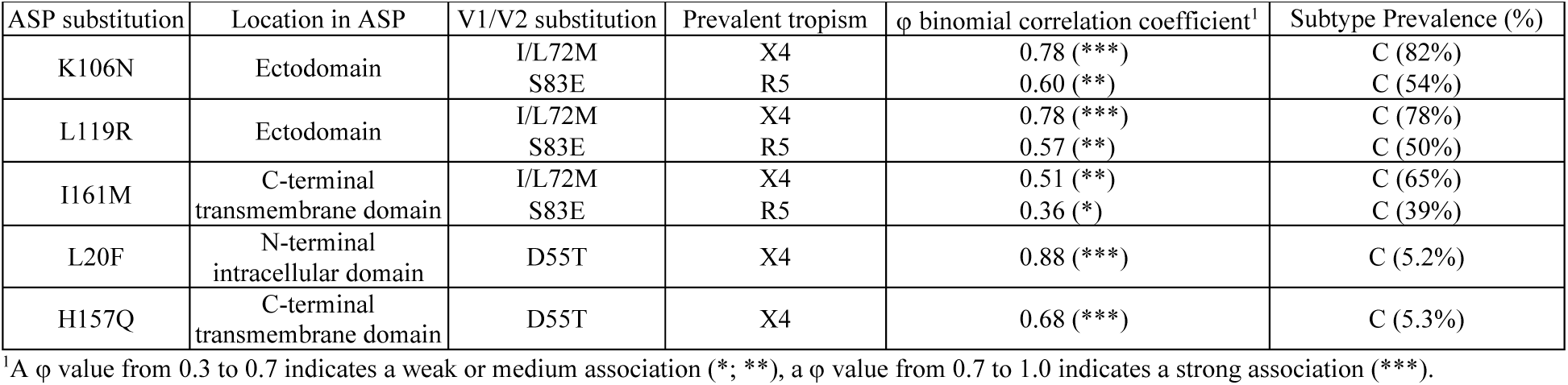
Amino acid substitutions in the ASP protein significantly associated with amino acid substitutions in the V1/V2 loop of ENV.

### 3.6 Extension of statistical analysis to the regions of gp120 protein outside V3 and V1/V2

We extended the analysis performed on V3 and V1/V2 to the remaining regions of gp120 protein. We considered these regions as controls, because they do not appear to be involved in viral tropism. We extracted from each of the 2496 *env* sequences with predicted tropism the following regions: *i*) the region upstream V1/V2 and not overlapping the *vpu* gene (from codon position 51 to 188 in the full-length aligned *env* sequences); *ii*) the region between V1/V2 and V3 (from codon position 351 to 478); *iii*) the region downstream V3 and not overlapping the antisense *asp* gene (from codon position 532 to 605).

After translation into amino acids, we obtained 3 datasets of 2496 sequences, aligned respectively for a total of 138, 128, and 74 positions. Within the 3 datasets, we selected respectively 88, 99, and 49 positions in which the frequency of gaps was <5%, and we compared their amino acid content in sequences R5-tropic with that in sequences X4-tropic using the χ^2^ contingency-table test. We found 3 positions in the region upstream V1/V2, 7 positions in the region between V1/V2 and V3, and 6 positions in the region downstream V3 showing a significant difference between R5 and X4 usage. Overall, we found that the fraction of amino acid sites associated with coreceptor usage is remarkably low (16 out of a total of 236; 6.7%), about one order of magnitude smaller than that found in V3 (20 out of 35; 57.1%) and half of that found in V1/V2 (10 out of 65; 15.4%). This result is in line with long established evidence that V3 and, to a lesser extent, V1/V2 are the determinants of co-receptor tropism.

The 3 positions we found in the region upstream V1/V2 account for 3 different amino acid substitutions. They show a significant covariation (φ from 0.36 to 0.86) with the ASP substitutions K106N, L119R and I161M (Table 4). The 7 positions we found in the region between V1/V2 and V3 account for 8 different amino acid substitutions. Four of them show a significant covariation (φ from 0.38 to 0.82) with the ASP substitutions K106N, L119R and I161M (Table 4). The 6 positions we found in the region downstream V3 account for 12 different amino acid substitutions. Five of them show a significant covariation (φ from 0.32 to 0.65) with the ASP substitutions K106N, L119R and I161M (Table 4). All these covariations are found only in subtype C strains, where they occur with a frequency from 20 to 89%. Unlike the V3 and V1/V2 regions, none of the amino acid substitutions found in the control regions show a significant covariation with the ASP substitutions L20F and H157Q.

**Table 4.** Amino acid substitutions in the ASP protein significantly associated with amino acid substitutions in three regions of ENV encoded in *env* sequences outside the overlap with *asp*.

| ASP substitution | Region of gp120 | Substitution | Prevalent tropism | $\phi$ binomial correlation coefficient<sup>1</sup> | Subtype Prevalence (%) |
| --- | --- | --- | --- | --- | --- |
| K106N | Upstream of V1/V2 | A1V | R5 | 0.63 (**) | C (64%) |
|  |  | T36K | R5 | 0.86 (***) | C (89%) |
|  |  | T51R | R5 | 0.37 (*) | C (26%) |
| L119R | Upstream of V1/V2 | A1V | R5 | 0.66 (**) | C (63%) |
|  |  | T36K | R5 | 0.84 (***) | C (85%) |
|  |  | T51R | R5 | 0.36 (*) | C (25%) |
| I161M | Upstream of V1/V2 | A1V | R5 | 0.42 (*) | C (48%) |
|  |  | T36K | R5 | 0.57 (**) | C (67%) |
| K106N | Between V1/V2 and V3 | V/T4A | R5 | 0.40 (*) | C (30%) |
|  |  | F28Y | R5 | 0.82 (***) | C (86%) |
|  |  | F99L | R5 | 0.67 (**) | C (79%) |
|  |  | N128V | R5 | 0.56 (**) | C (45%) |
| L119R | Between V1/V2 and V3 | V/T4A | R5 | 0.41 (*) | C (41%) |
|  |  | F28Y | R5 | 0.81 (***) | C (82%) |
|  |  | F99L | R5 | 0.69 (**) | C (76%) |
|  |  | N128V | R5 | 0.57 (**) | C (44%) |
| I161M | Between V1/V2 and V3 | V/T4A | R5 | 0.30 (*) | C (24%) |
|  |  | F28Y | R5 | 0.55 (**) | C (65%) |
|  |  | F99L | R5 | 0.45 (*) | C (59%) |
|  |  | N128V | R5 | 0.38 (*) | C (34%) |
| K106N | Downstream of V3 | A/V17S | R5 | 0.33 (*) | C (22%) |
|  |  | A/V17G | R5 | 0.34 (*) | C (20%) |
|  |  | Q28H | R5 | 0.65 (**) | C (67%) |
|  |  | V/I48K | R5 | 0.36 (*) | C (27%) |
|  |  | G70R | R5 | 0.63 (**) | C (69%) |
| L119R | Downstream of V3 | A/V17S | R5 | 0.32 (*) | C (21%) |
|  |  | A/V17G | R5 | 0.34 (*) | C (20%) |
|  |  | Q28H | R5 | 0.65 (**) | C (63%) |
|  |  | V/I48K | R5 | 0.34 (*) | C (25%) |
|  |  | G70R | R5 | 0.61 (**) | C (66%) |
| I161M | Downstream of V3 | Q28H | R5 | 0.45 (*) | C (51%) |
|  |  | G70R | R5 | 0.39 (*) | C (51%) |

<sup>1</sup>A $\phi$ value from 0.3 to 0.7 indicates a weak or medium association (\*; \*\*), a $\phi$ value from 0.7 to 1.0 indicates a strong association (\*\*\*)

The results from the analysis of all five regions of gp120 (upstream of V1/V2, V1/V2, between V1/V2 and V3, V3, and downstream of V3) are summarized in Table 5. When the five regions were combined in two groups (group 1 = V1/V2 + V3; group 2 = upstream of V1/V2 + between V1/V2 and V3 + downstream of V3), we found that the mean value of the correlation coefficient φ of group 1 (0.61; sd = 0.18) is significantly greater than that of group 2 (0.52; sd = 0.17; P = 0.03, one-tailed t-Student test).

**Table 5.**
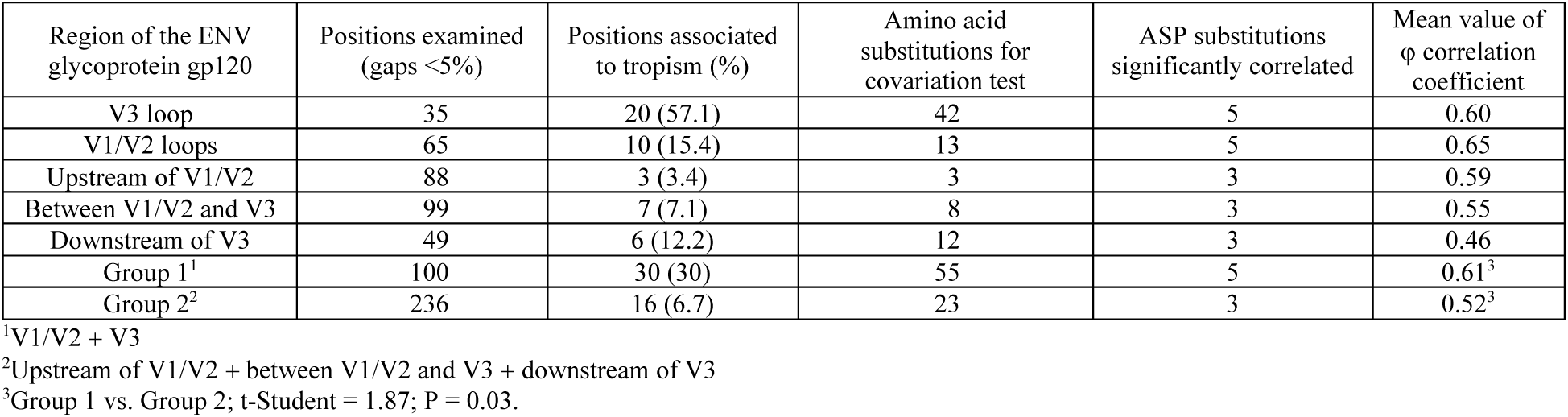
Sequence analysis for coreceptor tropism and covariation test for five different regions of the gp120 protein.

| Region of the ENV glycoprotein gp120 | Positions examined (gaps <5%) | Positions associated to tropism (%) | Amino acid substitutions for covariation test | ASP substitutions significantly correlated | Mean value of $\phi$ correlation coefficient |
| --- | --- | --- | --- | --- | --- |
| V3 loop | 35 | 20 (57.1) | 42 | 5 | 0.60 |
| V1/V2 loops | 65 | 10 (15.4) | 13 | 5 | 0.65 |
| Upstream of V1/V2 | 88 | 3 (3.4) | 3 | 3 | 0.59 |
| Between V1/V2 and V3 | 99 | 7 (7.1) | 8 | 3 | 0.55 |
| Downstream of V3 | 49 | 6 (12.2) | 12 | 3 | 0.46 |
| Group 1<sup>1</sup> | 100 | 30 (30) | 55 | 5 | 0.61<sup>3</sup> |
| Group 2<sup>2</sup> | 236 | 16 (6.7) | 23 | 3 | 0.52<sup>3</sup> |

<sup>1</sup>V1/V2 + V3<sup>2</sup>Upstream of V1/V2 + between V1/V2 and V3 + downstream of V3<sup>3</sup>Group 1 vs. Group 2; t-Student = 1.87; P = 0.03.

## 4. Discussion

The aim of this study was to identify amino acid signatures in the HIV-1 antisense protein ASP that are associated with coreceptor tropism. We have identified five positions of ASP where specific amino acid substitutions correlated with amino acid substitutions in the V3 and V1/V2 loops of gp120, which in turn are significantly associated with R5- or X4-tropism.

Identification of ASP amino acid signatures that are associated with co-receptor tropism required first the development of a computational method capable of predicting HIV-1 tropism from the amino acid sequence of the gp120 V3 loop with high sensitivity and specificity. Several computational methods for predicting viral tropism have been developed [9–19] and significant efforts are still underway to improve the accuracy of genotypic tests based on the sequence of the V3 loop domain. Given the considerable structural flexibility and sequence variability of the V3 loop, individual features of this region distinguishing between the two virus phenotypes R5- and X4-tropic are complex and difficult to define. Multivariate statistical methods [21] have the ability to summarize the information carried by individual variables into a few synthetic variables (e.g., the principal component analysis) or into a single synthetic variable (e.g., the linear discriminant analysis, LDA), leading to a better understanding of the biological process under examination [32–34]. In this study, we used the Fisher’s linear discriminant analysis [20,35] to find the best combination of amino acid physicochemical features in distinguishing between the two virus phenotypes R5- and X4-tropic. Our study showed that LDA is a promising approach to predict the viral tropism both in terms of sensitivity and specificity (Table 1). The ability of LDA to maximize the variance between groups and minimize the variance within groups can be appreciated by examining the 3 distributions of the LDA score in V3 sequences R5- and X4-tropic (Figure 1).

Despite the advantages, our method presents a few limitations. A first limitation is that it predicts the tropism only for genotypes B and C. Although they account respectively for 11.3% and 50.4% of the global genetic diversity of HIV-1 [36], LDA did not analyze V3 sequences from genotype A (global prevalence of 12.4%), from genotypes D, F, and G (6.4%), from circulating recombinant forms (15.1%), and from unique recombinant forms (2.0%). Another limitation is the low number of V3 sequences X4-tropic, only 137, in our training dataset. Although most of the V3 sequences from viral samples are in the R5-tropic group, it would be important to increase as much as possible the number of V3 sequences in the X4-tropic group. This should improve, in turn, the ability of the two validation tests to assess the robustness of LDA.

Although LDA was originally developed to classify subjects into one of the two clearly defined groups, it was later expanded to classify subjects into more than two groups. Indeed, as shown in his seminal study [20], Fisher successfully developed a linear discriminant model to distinguish three related species of the genus *Iris* from each other. Therefore, we could extend LDA to three groups: V3 sequences from strains R5-, X4-, and R5X4-tropic strains. It would be interesting to determine how dual tropic strains stack up against strains R5- and X4-tropic strains and test whether the switch from one tropism to the other is driven by small, but significant, changes in the sequence properties of the V3 loop domain.

After the initial discovery of the *asp* gene by Miller [37], Cassan et al. showed that an intact *asp* ORF is present only in pandemic HIV-1 strains (group M), and it is absent in all other primate lentiviruses, including non-pandemic HIV-1 groups [38]. We showed the existence of selective pressure that maintains an intact *asp* ORF in the HIV-1 genome by conserving the *start* codon and by avoiding early *stop* codons [23]. Recently, we proposed a role for the ASP protein in promoting intra-host viral spread or pathogenesis, based on a significant association between the frequency of viral strains encoding a full-length ASP and disease progression [22]. An extensive sequence analysis by Dimonte first hypothesized an involvement of ASP in coreceptor selection through identification of amino acid substitutions associated with classical signatures of viral tropism in the V3 loop of gp120 [39].

The first question in this study was whether there are positions in ASP where amino acid substitutions are preferentially associated with coreceptor usage. From a dataset of 2593 aligned ENV sequences we extracted the corresponding V3 sequences and predicted the tropism by means of linear discriminant analysis. The percent frequency of V3 sequences with a predicted X4 tropism (235 out of 2496; 9.4%) is comparable to that of V3 sequences with a known X4 tropism (137 out of 1838 in the training dataset; 7.5%). Our analysis identified five amino acid positions in ASP where substitutions were preferentially associated with tropism (Figure 3). The number of positions and substitutions we found in ASP (5 and 5, respectively) is much smaller than the ones reported by Dimonte (36 positions for 58 substitutions) [39]. This large difference is due to the fact that we restricted our analysis exclusively to substitutions in ASP caused by mutations that are non-synonymous in the *asp* ORF but synonymous in *env*. These substitutions are not a byproduct and occur independently of ENV evolution.

The second question of this study was whether there is a significant covariation between the tropism-associated substitutions in ASP and the tropism-associated substitutions in the V3 loop domain. We first identified 20 amino acid positions in V3 that are significantly associated with tropism, including the classical ones 11 and 25 [9], for a total of 42 substitutions (Figure 2). In this case, the result is very similar to the one obtained by Dimonte, who identified 24 amino acid positions in V3 for 44 substitutions [39]. In response to the question, we found that all 5 substitutions in ASP associated with tropism show a significant covariation with 6 of the 42 substitutions in V3 associated to tropism (Table 2). It should be noted that substitutions K106N and L119R map in the ectodomain of ASP and are the ones most significantly associated with substitutions in V3 that determine coreceptor tropism (correlation coefficient φ from 0.56 to 0.87). Altogether, these findings support the hypothesis that ASP may be involved in coreceptor selection, which can be validated through *in vitro* and *in vivo* studies.

Crystal and cryo-electron microscopy structure studies on ENV are indicative of a concerted action of V1/V2 and V3 loops during HIV-1 entry [40,41]. They have revealed that V1/V2 loops are located at the trimer apex in connection with V3 loop, and that the conformational change of V1/V2 loops upon CD4 binding triggers exposure of V3 loop to the coreceptor. In the V1/V2 region we identified 10 amino acid positions and 13 substitutions significantly associated with tropism (Figure 4). Despite the greater length of V1/V2, these numbers were remarkably lower than the ones we found in V3 (20 positions and 42 substitutions), in line with the notion that the V3 loop is the major determinant of coreceptor tropism [6]. Interestingly, we found that all the 5 substitutions in ASP associated to tropism show a significant covariation with 3 of the 13 substitutions in V1/V2 associated to tropism (Table 3). Also in this case, the two substitutions K106N and L119R in the ASP ectodomain are the ones showing strongest covariation with substitutions in V1/V2 that are associated with coreceptor tropism (correlation coefficient from 0.56 to 0.87). Again, this finding supports the hypothesis of an ASP involvement in viral tropism.

Extending our analyses to regions of gp120 outside of the first three hypervariable loops (excluding the overlaps with the *vpu* and *asp* genes), we identified a total of 16 positions and 23 substitutions associated with co-receptor tropism. Of these, 12 show covariation with substitutions in ASP, with some significant differences (Table 4). First, covariation was observed with only 3 of 5 substitutions in ASP (K106N, L119R, and I161M). Second, the mean coefficient of covariation φ in three regions combined is significantly lower than the one observed for the combined V3 and V1/V2.

It is also noteworthy that the majority of covariations in V3, in V1/V2, and in the regions outside these three hypervariable loops were found almost exclusively in subtype C strains at frequencies as high as >90%. Subtype C is by far the most prevalent HIV-1 clade worldwide [38], suggesting a potential association between specific substitutions in ASP and viral spread within and among hosts. This hypothesis had already been proposed in a previous report [38], which indeed noted a correlation between the frequency of viral strains within each HIV-1 subtype containing an uninterrupted *asp* ORF and the prevalence of the subtype. This hypothesis is also supported by two recent studies. In the first one, we reported that viral isolates with an uninterrupted *asp* ORF are found at significantly higher frequency in people living with HIV-1 (PLWH) who progress to AIDS in <3 years (rapid progressors) than in those who progress in >12 years (long term non-progressors) [22]. Another recent study described the presence of antibodies to ASP in serum of PLWH [42]. Remarkably, these responses were found in viremic and elite controllers and were found to be directed primarily to epitopes mapping in the ectodomain of ASP [42].

Altogether, our studies suggest that amino acid substitutions in ASP are linked to substitutions in gp120 that are signatures of R5 or X4 coreceptor tropism. Whether the covariations identified in the present study have actual bearing on R5- versus X4-dependent viral entry remains to be addressed experimentally.

## Acknowledgements

This work was supported by National Institutes of Health grants AI144983 and AI172556 (F.R.). This work has benefited from the equipment and framework of the COMP-HUB and COMP-R Initiatives, funded by the “Departments of Excellence” program of the Italian Ministry for Uni- versity and Research (MIUR, 2023–2027).

