## Supplementary Files for "Covariation of amino acid substitutions in the HIV-1 envelope glycoprotein gp120 and the antisense protein ASP associated with coreceptor usage"

**Supplementary File S1**

Training dataset of 1838 amino acid sequences of the V3 loop domain with known tropism: 1701 sequences (1229 of genotype B and 472 of genotype C) are CCR5-tropic (R5) and 137 sequences (96 of genotype B and 41 of genotype C) are CXCR4-tropic (X4).

| NCBI ac. number | Genotype | Tropism | V3 amino acid sequence | length (aa) |
|---|---|---|---|---|
| HQ644967 | B | CCR5 | CTRPNNNTRKSIHIGPGRAFYATGDIIGDIRKAHC | 35 |
| AJ418502 | B | CCR5 | CTRLNNNTRKSIHMGPGRAFYTTGEIIGDIRQAHC | 35 |
| JN687773 | B | CCR5 | CTRPYNNTRRSIPIGPGRAFYATGEVIGNIRKAYC | 35 |
| DQ061525 | B | CCR5 | CIRPNNNTRKSIHIGPGRAFYATGEIIGDIRQAHC | 35 |
| HQ377462 | B | CCR5 | CTRPNNNTRKSISMGPGRAFYATGGIIGNIRQAHC | 35 |
| AF541040 | B | CCR5 | CTRPNNNTRKSIHIGPGRAFYTTGEIIGDIRQAHC | 35 |
| DQ061827 | B | CCR5 | CTRPNNNTRKGIHMGPGKVFYATGQIIGDIRQAHC | 35 |
| EF600077 | B | CCR5 | CTRPNNNTRKSIHIGPGRAFYATGDIVGDIRQAHC | 35 |
| EU604591 | B | CCR5 | CTRPNNNTRKSINIGPGRAWYATGQIIGDIRQAYC | 35 |
| EF600098 | B | CCR5 | CTRPNNNTRKGIHIGPGRAFYATGDIIGDIRQAHC | 35 |
| JN687814 | C | CCR5 | CTRPGNNTRKSIRIGPGQTFYATGEIIGDIRQAHC | 35 |
| AY835439 | B | CCR5 | CTRPGNNTRKSIHIGPGRAFYATGDIIGDIRQAHC | 35 |
| EU293448 | C | CCR5 | CTRPGNNTRQSIRIGPGQAFYTNNIIGDIRQAHC | 34 |
| JX973238 | C | CCR5 | CTRPGNNTRKSIRIGPGQTFFATGDVIGDIREAHC | 35 |
| FJ977092 | C | CCR5 | CTRPNNNTRKSIRIGPGQTFYATGDIIGDIRQAHC | 35 |
| EF643670 | B | CCR5 | CTRPSNNTRKSITIGPGRAFYATGEIIGDIRKAHC | 35 |
| DQ002184 | B | CCR5 | CTRPNNNTRRSITIGPGRAFYATDIIGDIRQAHC | 34 |
| FJ653401 | B | CCR5 | CTRPNNNTRKSIHIAPGRAFYATGDIIGDIRQAHC | 35 |
| HQ377491 | B | CCR5 | CTRPGNNTRKSIPIGPGRAFYATGDIIGDIRKAHC | 35 |
| KC156221 | C | CCR5 | CTRVGNNTRKSVRIGPGQTFYATGDIIGDIREAHC | 35 |
| AF259038 | B | CCR5 | CTRPGNNTRKGIHIGPGRAFYTTGQIIGNIRQAHC | 35 |
| GU455514 | B | CCR5 | CTRPNNNTRKGIHIGPGRAFYTTGQIIGDIRQAHC | 35 |
| JN687721 | C | CCR5 | CTRPNNNTRKSVRIGPGQAFYATNGIVGDIRQAHC | 35 |
| HQ644912 | B | CCR5 | CMRPNNNTRKSIHIGPGRAFYTTGDIIGDIRQAHC | 35 |
| JF896843 | B | CCR5 | CTRPNNNTRKSIHIGPGRAFYATGQIIGNIRQAHC | 35 |
| KC113010 | B | CCR5 | CTRPNNNTRKIHIGPGKTFFATEVIGIRKAHC | 32 |
| JN001998 | B | CCR5 | CTRPNNNTRKSIHITPGRAFYATGEKIGDIRQAHC | 35 |
| KF770337 | C | CCR5 | CIRPGNNTRKSMRIGPGQTFYATGEIIGDIRQAHC | 35 |
| KC312501 | B | CCR5 | CTRPNNNTRKGIHIGPGRTFYVTGQIIGDIRQAHC | 35 |
| HM215411 | C | CCR5 | CTRPNNNTRRSIRIGPGQTFFAHGEIIGDIRQAHC | 35 |
| HQ708060 | C | CCR5 | CTRPNNNTRKSVRIGPGQTFYATGIIGNIRQAYC | 34 |
| KC156283 | C | CCR5 | CTRPNNNTRKSMRIGPGQTFYATGAIIGNIRQAYC | 35 |
| KC156440 | C | CCR5 | CIRPNNNTRKSVRIGPGQTFYATGEIIGDIREAYC | 35 |
| AY010823 | B | CCR5 | CTRPNNNTRKGIHIGPGSAIYATGDIIGDIRQAHC | 35 |
| KC156432 | C | CCR5 | CTRPNNNTRRSVRIGPGQTFYATGEIIGDIREAYC | 35 |
| DQ002206 | B | CCR5 | CTRPNNNTRKSIPIGPGRAFYATGEIIGDIRQAYC | 35 |
| JN002047 | B | CCR5 | CTRPNNNTRKSIPIGPGRAIYTTGGIIGDIRQAHC | 35 |
| EU744095 | B | CCR5 | CTRLNNNTRKSITFGPGRAFYATGDIIGNIRQAQC | 35 |
| EU578419 | B | CCR5 | CTRPNNNTRRGVTIGPGRVFYTGEVTGDIRQAHC | 34 |
| EU577028 | B | CCR5 | CTRPNNNTRKGITIGPGSVFYTGEIIGDIRQAHC | 34 |
| JQ779187 | C | CCR5 | CTRPGNNTRRSVRIGPGQTFYATGQIIGNIREAHC | 35 |
| AF153133 | C | CCR5 | CTRPNNNTRKSMRIGPGQTFYATGDIIGDIRQAHC | 35 |
| AF541049 | B | CCR5 | CTRPSNNTRKSIPMGPGKAFYATGDITGDIRKAHC | 35 |
| AF541010 | B | CCR5 | CTRPNNNTRKSIPMGPGKAFYATGDIIGDIRKAHC | 35 |
| JX972948 | C | CCR5 | CTRPNNNRRRSVRIGPGQSFYAYNDIIGNIREAYC | 35 |
| AF153144 | C | CCR5 | CTRPNNNTRKSIRIGPGQTFYANNAIIGNIRQAHC | 35 |
| DQ235642 | C | CCR5 | CTRPNNNTRKSIRIGPGQAFYATNDIIGDIRQAHC | 35 |
| DQ061825 | B | CCR5 | CTRPNNNTRKGIHMGSGAVFYATGQIIGDTRQAHR | 35 |
| HQ377375 | B | CCR5 | CTRPNNNTRKGIHLGPGGAFYTTGEIIGDIRKAHC | 35 |
| KC312591 | B | CCR5 | CVRPHNNTRKSIRIGPGSTFYATGEVIGDIRQAHC | 35 |
| HQ708066 | C | CCR5 | CIRPYNNTRKSMRIGPGQTFYATEEVIGDIRQAYC | 35 |
| AY010777 | B | CCR5 | CIRPNNNTRRSIHMGPGKAFYATGDIIGDIRQAYC | 35 |
| AF153152 | C | CCR5 | CTRPYNNTRKSMRIGPGQTFYATGDIIGDIRQAHC | 35 |
| EU744114 | B | CCR5 | CTRPNNNTRKSIHIGPGRAFYATGEIIGNIRQAHC | 35 |
| JF896859 | B | CCR5 | CTRPNNNTRRSIGIGPGRAFYATGIIGDIRQAHC | 34 |
| HM179798 | C | CCR5 | CIRPNNNTRTSIRIGPGQAFFATNGIIGNIRQAYC | 35 |
| AY253308 | C | CCR5 | CTRPGNNKRRSVRIGPGQTFYATGEIIGNIREAHC | 35 |
| HM179719 | C | CCR5 | CTRPNNNTRKSMRIGPGQAFYATGEIIGNIREAHC | 35 |
| FJ653102 | B | CCR5 | CTRPNNNTRKSINIGPGRAFYAATDIIGDIRQAHC | 35 |
| DQ061505 | B | CCR5 | CTRPSNNTRKSITIGPGRAFYTTGEILGEIRQAHC | 35 |
| HQ644953 | B | CCR5 | CTRPNNNTRKSIHMGPGRAFYATGEIIGDIRQAHC | 35 |
| HQ644892 | B | CCR5 | CTRPNNNTRKGIHIGPGRAFYTTGEITGDIRKAHC | 35 |

| | | | | |
|---|---|---|---|---|
| FJ375997 | C | CCR5 | TRPNNNTRKSVRIGPGQTFYATGDIIGDIRQAHC | 34 |
| DQ235622 | C | CCR5 | CARPNNNTRKSIRIGPGQAFYATGEIIGNIRQAHC | 35 |
| EU744023 | B | CCR5 | CTRPSNNTRKSIHIGPGRAFYATGEIIGNIRQAHC | 35 |
| EU604569 | B | CCR5 | CTRPSNNTRKSINMGPGRAFYTTGEIIGDIRQAHC | 35 |
| HQ644821 | B | CCR5 | CTRPNNNTRKSIHIGPGRAFYATGEIIGDIRQAHC | 35 |
| DQ869033 | B | CCR5 | CTRPNNNTRKGVHIGPGRAFYATGEIIGDIRKAHC | 35 |
| U08717 | B | CCR5 | CTRPNNNTRKSIPLGPGQAWYTTGQIIGDIRQAHC | 35 |
| DQ002106 | B | CCR5 | CTRPNNNTRKSITIGPGRAFYATGDIIGDIRQAHC | 35 |
| AF541016 | B | CCR5 | CTRPNNNTRKSIPIGPGRAFYTTGEIIGDIRQAHC | 35 |
| EU744048 | B | CCR5 | CTRPSNNTRKGIHIGPGRALYATGEIIGDIRQAHC | 35 |
| AF384315 | B | CCR5 | CTKPYKKKKSRRIHIGPGRTFHTTGSIGDIRRAHC | 35 |
| HQ377411 | B | CCR5 | CTRPNNNTRRSIHIGPGSAFYATGDIIGDIRQAHC | 35 |
| EU744165 | B | CCR5 | CTRPNNNTRRSIHMGPGRALYTTGAIIGNIRQAHC | 35 |
| AF384271 | B | CCR5 | CTRPGNNTRRSIRIGPGSAFYATGDIIGDIRKAHC | 35 |
| JF508028 | B | CCR5 | CTRPNNNTRKSINIGPGRAFYTTGEIIGNIRQAHC | 35 |
| KF770409 | C | CCR5 | CTRPGNNTRKSMRIGPGQTFYATGDIIGDIRKAHC | 35 |
| DQ061444 | B | CCR5 | CTRPSNNTRKSIHIGPGRAFYTTGEIIGDIRQAHC | 35 |
| HQ708056 | C | CCR5 | CIRPNNNTRKSVRIGPGQTFYATGDIIGDIRRAYC | 35 |
| DQ516156 | B | CCR5 | CTRPNNNTRKDIHIGPGRAFYATGDIIGDIRQAHC | 35 |
| AF199039 | B | CCR5 | CIRPNNNTRQGIHIGPGKALYTTIIGNIRQAHC | 33 |
| EU272303 | B | CCR5 | CTRPNNNTRKSIHLGPGRAWYATGEIIGNIRQAHC | 35 |
| DQ235621 | C | CCR5 | CVRPNNNTRKSIRIGPGQTFYATGDIIGNIRQAHC | 35 |
| JX508865 | C | CCR5 | CTRPNNNTRKSFRIGPGQTFYATGDIIGDIRQAHC | 35 |
| DQ002248 | B | CCR5 | CTRPNNNTRKSIHIAPGRAFYATGEIIGDIRQAHC | 35 |
| AM156917 | B | CCR5 | CTRPSNNTRKGIHIGPGRAFYATGEIIGDIRQAHC | 35 |
| FJ653141 | B | CCR5 | CTRFNNNTRKSIHIGPGRAFYATGEIIGNIRQASC | 35 |
| JX140657 | B | CCR5 | CIRPGNNTRKSITMGPGRAFYATGEIIGNIRQAHC | 35 |
| DQ002070 | B | CCR5 | CIRPNNNTRKSIHIGPGRAFYTTGDIIGDIRQAHC | 35 |
| DQ002035 | B | CCR5 | CTRPHNNTRKSIPMGPGRAFYTAGQIIGDIRQAHC | 35 |
| AJ418494 | B | CCR5 | CTRPNNNTRKSISFGPGSAMYATGAIIGDIRQAHC | 35 |
| DQ002144 | B | CCR5 | CTRPNNNTRRGIHIGPGRAFYTTGEIIGDIRQAYC | 35 |
| EF600078 | B | CCR5 | CTRPNNNTRKSIHIGPGRALYATGDIIGDIRQAHC | 35 |
| KC312489 | B | CCR5 | CTRPNNNTRKGIHIGPGRTFYATGEIIGDIRQAHC | 35 |
| AF540999 | B | CCR5 | CTRPNNNTRKSIPIGPGRALYATGEIIGQIRRAYC | 35 |
| EU272203 | B | CCR5 | CTRPNNNTRKSIHLGQGRAWYATGEIIGDIRQAHC | 35 |
| KC312390 | B | CCR5 | CTRPNNNTRKSIHIGPGSAFYTTGEIIGDIRQAHC | 35 |
| EF657940 | B | CCR5 | CIRPNNNTRKSIHMGPGGAFYATGGIIGNIRQAHC | 35 |
| JQ779286 | C | CCR5 | CIRPNNNTRKSVRIGPGQTFYATGDIIGDIREAYC | 35 |
| JQ779910 | C | CCR5 | CTRPGNNTRKSVRIGPGQTYFSTGEIIGNIRQAHC | 35 |
| HQ644825 | B | CCR5 | CTRPNNNTRRSIHIGPGKAFFATGDIIGDIRQAHC | 35 |
| AF541011 | B | CCR5 | CTRPNNNTGKSIPIGPGRAFYATGEIIGDIRQAHC | 35 |
| AF541008 | B | CCR5 | CTRPSNNTRKSIPMGPGKAFYATGDIIGDIRKAHC | 35 |
| KC312576 | B | CCR5 | CVRPHNNTRKGIHIGPGSTFYATGEVIGDIRQAHC | 35 |
| HM179933 | C | CCR5 | CTRPGNNTRKSMWIGPGQAFYATGDIIGDIRQAYC | 35 |
| FJ653234 | B | CCR5 | CTRPSNNTSGSIHIGPGRAFDATKTITGDIRQAHC | 35 |
| JQ779139 | C | CCR5 | CTRPGNNTRKSVRIGPGQTFYATGEIIGDIRKAHC | 35 |
| DQ002082 | B | CCR5 | CTRPNNNTRKSIHIGPGRAFYTTGSIIGDIRQAHC | 35 |
| AJ810480 | B | CCR5 | CTRPSNNTRKSVHIGPGRALYTTDIIGDIRKAYC | 34 |
| JX972470 | C | CCR5 | CTRPNNNTRKSVRIGPGQTFYATGEIIGDIRQAYC | 35 |
| EU744039 | B | CCR5 | CTRPSNNTRKGIHIGPGRAFYTTGEIIGDIRQAHC | 35 |
| AF022263 | B | CCR5 | CTRPNNNTRRSISIGPGRAFYTTGEIIGNIRQAHC | 35 |
| EU293445 | C | CCR5 | CIRTGNNTRKSVRIGPGQTFYATDGIIGDIRKAYC | 35 |
| DQ061432 | B | CCR5 | CTRPNNNTRKSITIGPGRAFYTTGEIIGDIRQAHC | 35 |
| EF579980 | B | CCR5 | CERPGNNTSKGIHIGPGRAFYATENIIGDIRKAHC | 35 |
| EU272329 | B | CCR5 | CTRPNNNTRKSIHLGQGRAWYTTGQIIGDIRQAHC | 35 |
| KC156238 | C | CCR5 | CIRPGNNTRRSMRIGPGQTFYATGDIIGDIRKAHC | 35 |
| EU576584 | B | CCR5 | CTRPNNNTRKGIHIGPGKAFYTTGEIIGNIRQAHC | 35 |
| EU744156 | B | CCR5 | CTRPNNNTRRSIHIGPGRAFYATGDIIGDIRQAHC | 35 |
| HQ644881 | B | CCR5 | CTRPSNNTRKSINIGPGRAFYATGDIIGDIRKAYC | 35 |
| HM239634 | B | CCR5 | CIRPNNNTRKSIHVGPGSALYTTKIIGNIRQAHC | 34 |
| HM239547 | B | CCR5 | CTRPNNNTRKSIPMGPGQALYATGEIIGDIRQAHC | 35 |
| EU744150 | B | CCR5 | CTRPNNNTRKSIHIGPGRAFYTTGGIIGDIRQAHC | 35 |
| HQ644861 | B | CCR5 | CTRPSNNTRKSIHMGPGRAFYVTGDVIGDIRQAHC | 35 |
| AF258965 | B | CCR5 | CTRPNNNTRKSIHLGPGRAFYTTGGIVGNIRQAHC | 35 |
| HM368250 | B | CCR5 | CIRPHNNTRKSIHIGPGRTFYATGDIIGDIRKAHC | 35 |
| AF541112 | B | CCR5 | CVRPNNNTRRGIHIGLGRRFYTTEIVGDIRKAYC | 34 |
| AF543913 | C | CCR5 | CTRPSNNTRKSVRIGPGQTFFATGDIIGDIRQAHC | 35 |
| HQ644883 | B | CCR5 | CTRPNNNTRKSIHIGPGGAFYATGEIIGDIRQAHC | 35 |
| JN687752 | B | CCR5 | CVRPNNNTRTSIHMGPGKAFYAAGEVIGDIRRAYC | 35 |
| U08771 | B | CCR5 | CTRPNNNTRKSIHMGWGRAFYATGEIIGNIRQAHC | 35 |

| | | | | |
|---|---|---|---|---|
| FJ653124 | B | CCR5 | CTRFYNNTRKSIHIGPGRAFYATGEIIGNIRQASC | 35 |
| AF491742 | B | CCR5 | CTRPNNNTRKSIPMGPGKAFYATGDIIGDIRQAHC | 35 |
| AF384280 | B | CCR5 | CIRPNNNTRTSIPMGPGRAWYAMGDIIGDIRQAHC | 35 |
| KF384808 | B | CCR5 | CTRPGNNTRKGITIGPGRAFYATGKIIGDIRQAHC | 35 |
| DQ061418 | B | CCR5 | CTRPNNNTRKSINIGPGRAFYTTGEIIGDIRQAHC | 35 |
| JN687784 | C | CCR5 | CTRPGNNTRRSVRIGPGQTFYATGDIIGNIRQAHC | 35 |
| JX972406 | B | CCR5 | CVRPGNNTRKSITIGPGRAFYATGEIIGDIRKAHC | 35 |
| JX972238 | B | CCR5 | CTRPNNNTRKSIHLGPGSAIYATGQIIGDIRQAHC | 35 |
| U08711 | B | CCR5 | CTRPNNNTRKSITIGPGRAFYATGEIIGDIRQAHC | 35 |
| KF770437 | C | CCR5 | CIRPGNNTSRSIRIGPGQAFYATGRIVGDIRQAHC | 35 |
| HM246239 | B | CCR5 | CTRPSNNTRKSISIGPGRAFYATGDIIGDIRQAHC | 35 |
| JF508012 | B | CCR5 | CTRPNTNTRKSINIGPGRAFYTTGEIIGNIRQAHC | 35 |
| AY010852 | B | CCR5 | CTRPSNNTRKSIYIGPGRAFYATGSIIGDIRQAHC | 35 |
| EU744106 | B | CCR5 | CTRPNNNTRKSINIGPGRALYTTGEIIGNIRQAHC | 35 |
| DQ516243 | B | CCR5 | CTRPNNNTRKGIHIGPGRAFYTTGEIIGDIRQAYC | 35 |
| JF896862 | B | CCR5 | CTRPGNNTRKGIHMGPGKAFFTTETVIGNVRQAHC | 35 |
| EF579987 | B | CCR5 | CERPGNNTSKGIHIGPGRAFYATENIIGDIRKARC | 35 |
| AF541098 | B | CCR5 | CIRPNNNTKRSIHLGLGRRFYTTEIVGDIRKAYC | 34 |
| KF770305 | C | CCR5 | CTRPDNNTRRSVRIGPGQVFYTNDIIGDIRRAHC | 34 |
| DQ061835 | B | CCR5 | CTRPNNNTRKGIHMGPGKVFYATGQIIGDIRQAYC | 35 |
| FJ653296 | C | CCR5 | CTRPNNNTRKSIRIGPGQTFFATEIVGNIRQAHC | 34 |
| AF384307 | B | CCR5 | CTRPNNNTRKSIHIGPGRAFYATGDIIGDIRQAHC | 35 |
| U08716 | B | CCR5 | CTRPNNNTRKSIHLGPGRAWYTTGQIIGDIRQAHC | 35 |
| JX973081 | C | CCR5 | CIRPGNNTRKSVRIGPGQAFYATGDIIGDIRQAHC | 35 |
| DQ235628 | C | CCR5 | CTRPNNNTRKSIRIGPGQTFYATNEIIGDIRQAHC | 35 |
| HQ708084 | C | CCR5 | CTRPNTNKRTSYTIGPGRAFFATGDIIGDIRRADC | 35 |
| JF507865 | B | CCR5 | CTRPGNNTRRSINIGPGRAFYTTGDIIGNIRQAHC | 35 |
| HM246216 | B | CCR5 | CTRPNNNTRKSIHIAPGGAFYATGDIIGDIRQAHC | 35 |
| AJ418500 | B | CCR5 | CTRLNNNTRKSIHMGPGRAFFATGEIIGDIRQAHC | 35 |
| EU272197 | B | CCR5 | CTRPNNNTRKSINLGPGQAWYTTGQIIGDIRQAHC | 35 |
| JN687771 | B | CCR5 | CTRPNNNTRKSINIGPGRAWYATGEIIGNIRQAHC | 35 |
| HQ708028 | C | CCR5 | CTRPDNNTRKSVRIGPGQTFYTTGDIIGNIRQAYC | 35 |
| HQ708094 | C | CCR5 | CTRPNNNIRKSYTIGPGRAFFATGDIIGDIRQADC | 35 |
| DQ061797 | B | CCR5 | CIRPNNNTRKGIHIGPGRAFYATGEIIGNIRQAHC | 35 |
| KF770354 | C | CCR5 | CTRPNNNTSQSIRIGPGQTFYAMGRIIGDIRQAHC | 35 |
| DQ061529 | B | CCR5 | CTRPNNNTRRSIHIAPGSTFFATGDIIGDIRQAHC | 35 |
| JX508864 | C | CCR5 | CTRPNNNTRQSFRIGPGQTFYATGDIIGDIRQAHC | 35 |
| DQ061693 | B | CCR5 | CIRPNNNTRKGVHLGPGGALYATGAIIGDIRQAYC | 35 |
| HQ708047 | C | CCR5 | CTRPNNNTRKSIRIGPGQTFYATGAVTGDIRKAYC | 35 |
| EF688444 | B | CCR5 | CTRPNNHTRKRMTLGPGRVYYTTGEIVGDIRQAHC | 35 |
| FJ653195 | B | CCR5 | CTRPNNNTRKGIHIGPGRAFYATEKITGDIRQAHC | 35 |
| KF770374 | C | CCR5 | CTRPGNNTSRSIRIGPGQTFYATGRITGNIRQAHC | 35 |
| HM179822 | C | CCR5 | CTRPGNNTRQSMRIGPGQTFYATGDIIGDIRPAHC | 35 |
| KF770306 | C | CCR5 | CTRPDNNTRRSVRIGPGQVFYTNDIIGDIRQAHC | 34 |
| EF600079 | B | CCR5 | CTRPNDNTRKSIHIGPGRALYATGDIIGDIRQAHC | 35 |
| DQ002072 | B | CCR5 | CIRPNNNTRKSIHIGPGRVFYTTGDIIGDIRQTHC | 35 |
| HQ644954 | B | CCR5 | CTRPNNNTRRSINIGPGRAFYATGEIIGDIRQAHC | 35 |
| JF508059 | B | CCR5 | CTRPNNNTRKSIHIGPGRAFYATGEIIGNIRQASC | 35 |
| AF180900 | B | CCR5 | CTRPNNNTRRSIHIGPGRAFYTTGQIIGDIRQAYC | 35 |
| DQ002105 | B | CCR5 | CTRLNNNTRRSINIGPGRAWYTTGEVIGDIRKANC | 35 |
| AJ418531 | B | CCR5 | CTRPHNNTRKSIHIGPGRAFYATGEIIGDIRQAHC | 35 |
| HQ645007 | B | CCR5 | CTRPSNNTRKSINFGPGRAIYTTGQIIGDIRQAHC | 35 |
| HM239583 | B | CCR5 | CTRPNNNTRKSINIGPGKALYATGDIIGDIRQAHC | 35 |
| EF657913 | B | CCR5 | CMRPNNNTRKSIHMGPGRAFYATGEIIGNIRQAHC | 35 |
| AF021510 | B | CCR5 | CTRPNNNTRKSIHIGPGRAFYTTGQIIGDIRQAYC | 35 |
| JF507860 | B | CCR5 | CTRPNNNTRKSINIGPGRAFYTTGQIIGDIRQAHC | 35 |
| JF508121 | B | CCR5 | CTRPNNNTRKSIHIGPGRTFYATGEIIGDIRQAHC | 35 |
| HQ644851 | B | CCR5 | CTRPNNNTRRSITIGPGRAFYTTDIIGDIRQAYC | 34 |
| DQ235636 | C | CCR5 | CTRPGNNTRKSVRIGPGQTFYATGEIIGDIRRAHC | 35 |
| DQ002324 | B | CCR5 | CTRPNNNTRRSIHMGPGRAFYTTGEIVGDIRQAYC | 35 |
| HM179926 | C | CCR5 | CTRPNNNIRKSIRIGPGQTFYATGDIIGDIRQAYC | 35 |
| JF896850 | B | CCR5 | CTRPNNNTRKSIHMGPGKAFYTAGDIIGDIRKAHC | 35 |
| AF391249 | C | CCR5 | CTRPNNNTRRSMRIRPGQTFYATGEIIGDIRQAYC | 35 |
| DQ516119 | B | CCR5 | CTRPNNNTIKSIHIGPGRAFYTTGQIIGNIRQAHC | 35 |
| DQ002304 | B | CCR5 | CTRPNNNTRKGIHIGPGRAFYATGDIIGDIRKAHC | 35 |
| AF540997 | B | CCR5 | CTRPNNNTRKSIPIGPGRAFYATGDIIGDIRQAHC | 35 |
| AY010871 | B | CCR5 | CTRPNNNTRTSIPMGPGRAMYATGDIIGNIRQAYC | 35 |
| DQ061462 | B | CCR5 | CTRPNNNTRKGIHIGPGRASYTTGEIIGDIRKAHC | 35 |
| DQ061538 | B | CCR5 | CTRPNNNTRRGIHIAPGSAFYATGEIIGDIRQAHC | 35 |
| JF896815 | B | CCR5 | CTRNNNRRGHVGPGGALFTNHIIGDIRQAHC | 31 |

| | | | | |
|---|---|---|---|---|
| AY669739 | C | CCR5 | CTRPNNNTRRSIRIGPGQVFYANNDIIGDIRQAHC | 35 |
| KF770352 | C | CCR5 | CTRPGNNTRRSVRIGPGQAFYATGDIIGDIRAAHC | 35 |
| AF153185 | C | CCR5 | CIRPGNNTRKSVRIGPGQAFFATGDIIGDPRQAHC | 35 |
| EU272291 | B | CCR5 | CTRPSNNTRKSINLGPGTAWYTTGQIIGDIRQAHC | 35 |
| AY887888 | C | CCR5 | CTRPGNNTRKSMRIGPGQTFYATGAIIGDIRQAHC | 35 |
| HQ644811 | B | CCR5 | CTRPNNNTRRSINIGPGRAIFTTGEIIGDIRQAHC | 35 |
| JF508074 | B | CCR5 | CTRPNNNTRKSIHIGPGKAFYTTGEIIGDIRQAHC | 35 |
| HM239611 | B | CCR5 | CTRPNNNTRKSIHIGPGRAFYATGGIIGNIRHAYC | 35 |
| EF579995 | B | CCR5 | CERPGNNTSKGIHIGPGRAFYATENIVGDIRKAHC | 35 |
| JF507729 | B | CCR5 | CTRPNNNTRKGIHIGPGRTFFATGEIIGDIRRAHC | 35 |
| JQ779248 | C | CCR5 | CIRPGNNTRKSIRIGPGQVFFAATDIIGNIREAHC | 35 |
| DQ002306 | B | CCR5 | CTRPNNNTRKSINIGPGRAWYATGDIIGDIRKAHC | 35 |
| HQ678254 | B | CCR5 | CTRPNNNTRKGIHMGPGGAFYTRGDIIGDIRKAHC | 35 |
| FJ653236 | B | CCR5 | CTRPSNNTSQSIHIGPGRAFDATKTITGDIRQAHC | 35 |
| JX972841 | C | CCR5 | CTRPNNNTRTSIRIGPGQTFYATGDIIGEIRQAYC | 35 |
| GQ372988 | B | CCR5 | CTRPNNNTRKSIHVGWGRALYTTGQIIGDIRKAHC | 35 |
| EU578319 | B | CCR5 | CTRPNNNTRKSIHIGPGRAFYTGEIIGDIRQAHC | 34 |
| AY010797 | B | CCR5 | CIRPNNNTRKSIHMGPGKAFYATGDIIGDIRQAYC | 35 |
| DQ422948 | C | CCR5 | CTRPGNNTRKSVRIGPGQTFYATGAIIGDIRKAHC | 35 |
| HM239629 | B | CCR5 | CTRPNNNTRKGIPVGPGKAIYATGEIIGNIRQAHC | 35 |
| AY842838 | B | CCR5 | CIRPNNNTRKSINIGPGRAMYATEQIIGDIRQAHC | 35 |
| HM179732 | C | CCR5 | CARPNNNTRKSMRIGPGQAFYATGEIIGNIREAHC | 35 |
| HQ644828 | B | CCR5 | CTRPNNNTRRSIHIGPGRAFFATGDIIGDIRQAHC | 35 |
| HQ644920 | B | CCR5 | CMRPQNNTRKSIPIGPGRAFYTTGDIIGDIRQAHC | 35 |
| EF600086 | B | CCR5 | CIRPNNNTRQGIHIGPGKALYTTKIIGNIRQAHC | 34 |
| KC473834 | B | CCR5 | CTRPNNNTRKSINIGPGRAFYTTGDIIGDIRQAHC | 35 |
| KF770320 | C | CCR5 | CTRPNNNTRKSIRIGPGQTFFATDDIIGDIRKAYC | 35 |
| DQ235640 | C | CCR5 | CTRPNNNTRKSIRIGPGQTFYATNDIIGDIRQAHC | 35 |
| HM246201 | B | CCR5 | CTRPNNNTRKSINIGPGRAFYVAGEIIGDIRQAHC | 35 |
| JF507887 | B | CCR5 | CTRPNNNTKKSVNIGPGRAIYTTDIIGDIRKAYC | 34 |
| EU575490 | B | CCR5 | CTRPNNNTRRSIHIGPGRAFYTTGEVIGDIKQAHC | 35 |
| KC156324 | C | CCR5 | CTRPNTNIRKSMRIGPGQTFYATGDIIGDIRQAHC | 35 |
| JN002017 | B | CCR5 | CTRPNNNTRKGIHIGPGRAFYATGDIVGDIRQAHC | 35 |
| KC312494 | B | CCR5 | CTRPNNNTRKGIHIGPGRTFYATGEVIGNIRQAHC | 35 |
| HM179941 | C | CCR5 | CTRPGNNTRKSMWIGPGQVFYATGDIIGDIRQAYC | 35 |
| JX140664 | C | CCR5 | CTRPNNNTRKSIRIGPGQAFYATGDIIGNIREAHC | 35 |
| DQ061734 | B | CCR5 | CTRPNNNTRRSINLGPGRAYYATGDIIGDIRQAHC | 35 |
| KC156117 | C | CCR5 | CTRPSNNTRKSVRIGPGQTFYATGRIIGDIREAHC | 35 |
| EU272299 | B | CCR5 | SRPNNNTRKSIHLGLGRAWYATGEIIGNIRQAHC | 34 |
| DQ002334 | B | CCR5 | CTRPSNNTRKSIHIAPGRAFYATGDIIGNIRQAYC | 35 |
| DQ061412 | B | CCR5 | CTRPNNNTRKSITIGPGRAFYTAGEIIGDIRQAHC | 35 |
| AY010854 | B | CCR5 | CTRPSNNTRKSIHIGPGRAFYTTGAIIGDIRQAHC | 35 |
| EU293447 | C | CCR5 | CIRPNNNTRESIRIGPGQAFYATGGIIGDIRQAHC | 35 |
| JN002023 | B | CCR5 | CTRPNNNTRKSIHIGPGRAFYATGDIVGDIREAHC | 35 |
| HQ678294 | B | CCR5 | CTRPSNNTRRSIHMGPGKAYYTTDIIGDIRQAHC | 34 |
| KF384814 | B | CCR5 | CVRPNNNTRRSIHIGPGSAFYTTDIIGDIRQAHC | 34 |
| DQ061768 | B | CCR5 | CTRPSNNTRKGIHIGPGRAFYATGEIIGNIRQAHC | 35 |
| KC156321 | C | CCR5 | CNRPHNNTRKSMRIGPGQAFYATGDVTGDIRQAHC | 35 |
| AF153135 | C | CCR5 | CTRPNNNTRKSMRIGPGQTFYATGEIIGDIRQAHC | 35 |
| AY529676 | C | CCR5 | CTRPNNNTRQSVRIGPGQVFYATNDIIGDIRQAYC | 35 |
| DQ061537 | B | CCR5 | CTRPNNNTRRSIHIAPGSAFYATGEIIGDIRQAHC | 35 |
| AF541025 | B | CCR5 | CTRPNNNTRKSIHIGPGRAFYTTGEIIGNIRQAHC | 35 |
| DQ061583 | B | CCR5 | CMRPNNNTRKSIPIGPGRAFYTTGEIIGDIRQAHC | 35 |
| DQ388514 | C | CCR5 | CTRPNNNTRKSIRIGPGQTFYATGEIIGKIREAHC | 35 |
| AF153157 | C | CCR5 | CERPGNNTRKSVRIGPGQTFYATGEIIGNIRQAHC | 35 |
| HQ644853 | B | CCR5 | CTRPGNNTRKSINIGPGRALYTTGDIIGDIRQAHC | 35 |
| JF896836 | B | CCR5 | CTRPNNTKSIHIGPGRAFYATGEIIGDIQAHC | 32 |
| HQ377436 | B | CCR5 | CTRPNNNTRKSIHIGPGKAFYATGDIIGDIRQAHC | 35 |
| HQ377417 | B | CCR5 | CTRPNNNTRKSIHIGPGSAFYATGEIIGDIRQAHC | 35 |
| FJ375979 | C | CCR5 | ARPNNNTRKSVRIGPGQTFATGEIIDIRQAHC | 32 |
| KF770294 | C | CCR5 | CIRPNNNTRKSVRIGPGQTFYATGEVIGDIRKAHC | 35 |
| HQ377483 | B | CCR5 | CTRPGNNTRKSIPIGPGRAFYATGDIIGDVRKAHC | 35 |
| HQ644873 | B | CCR5 | CTRPNNNTRKGIHIGPGRAFYATGEIIGDIRQAHC | 35 |
| AY842814 | B | CCR5 | CTRPNNNTRKSIPIGPGRAMYATGDIIGDIRQAHC | 35 |
| EU744061 | B | CCR5 | CTRPNNNTRKSIHVGPGKTLYATGDIIGDIRQAHC | 35 |
| DQ869029 | B | CCR5 | CTRPNNNTRKSIHMGPGKAFYATGEIIGDIRKAYC | 35 |
| HM368238 | B | CCR5 | CTRPNNNTRRGIHIGPGGAFYSTGDIIGDIRQAHC | 35 |
| DQ061501 | B | CCR5 | CTRPSNNTRKSIHIGPGRAVYTTGEIIGRIRQAHC | 35 |
| DQ177208 | B | CCR5 | CTRLNNNTRKSIPMGPGRAFYATGDIIGDIRKAHC | 35 |
| EU744004 | B | CCR5 | CTRPNNNTRKSIPIGPGRAFYATGDIIGDIRKAHC | 35 |

| | | | | |
|---|---|---|---|---|
| EU744147 | B | CCR5 | CTRPNNNTRKGIHIGPGRAFYTTGEIIGDIRQAHC | 35 |
| DQ002187 | B | CCR5 | CTRPGNNTRKSIHIGPGRAFYATDIIGDIRQAYC | 34 |
| AY010805 | B | CCR5 | CTRPNNNTRKGIHLGPGSAFYATGDIIGDIRQAHC | 35 |
| AF082837 | B | CCR5 | CTRPNNNTRKSIHIGPGRAFYATGDNIGDIRQAHC | 35 |
| EU744084 | B | CCR5 | CTRLNNNTRKSITLGPGRAFYATGDIIGNIRQAQC | 35 |
| HM179838 | C | CCR5 | CTRPGNNTRQSMRIGPGQTFYATGDIIGDIRQAHC | 35 |
| HM246191 | B | CCR5 | CTRPSNNTRKSIPIGPGRAFYATGDIIGDIRQAHC | 35 |
| AY510062 | C | CCR5 | CTRPGNNTRKSVRIGPGQTFYATGAIIGDIRQAHC | 35 |
| AF153158 | C | CCR5 | CTRPNNNTRKGVRIGPGQTFYATGDIIGDIRQAHC | 35 |
| AY570011 | B | CCR5 | CTRPNNNTRKSINLGPGRALYTTGEITGDIRQAHC | 35 |
| DQ002323 | B | CCR5 | CTRPNNNTRRSIHMGPGRAFYTTGEIIGDIRQAYC | 35 |
| JF508007 | B | CCR5 | CTRPNNNTRKSINIGPGRAFYTTGTIIGDIRQAHC | 35 |
| KC156401 | C | CCR5 | CIRPGNNTRKSIRIGPGQTDFSTGEIIGNIRQAHC | 35 |
| AF384268 | B | CCR5 | CTRPDNNTRKSINIGPGRAFFATGDIIGDIRQAHC | 35 |
| DQ061772 | B | CCR5 | CTRPDNNTRKGIHIGPGRAFYATGEIIGNIRQAHC | 35 |
| DQ382371 | C | CCR5 | CARPGNNTRRSVRIGPGQAFYATGEIIGDIRKAHC | 35 |
| HM239626 | B | CCR5 | CTRPNNNTRRGIHIGPGRTLFATDIIGDIRQAHC | 34 |
| KF770366 | C | CCR5 | CIRPGNNTRKSVRIGPGQTFYATGDIIGDIRQAHC | 35 |
| AY010851 | B | CCR5 | CTRPNNNTRKSIHLGPGSAFYATGDIIGDIRQAHC | 35 |
| AF384246 | B | CCR5 | CTRPNNNTRKSIHIAPGRTYYATGDIIGDIRQAHC | 35 |
| AF153155 | C | CCR5 | CTRPGNNTRKSIRIGPGQAFYATGDIIGDIRQAHC | 35 |
| HQ644905 | B | CCR5 | CTRPTNNTRKGIHIGPGRAFYTTGEIIGDIRKAHC | 35 |
| DQ002233 | B | CCR5 | CTRPSNNTRKSIPIGPGRAFYATGDIIGDIRKAHC | 35 |
| JF896847 | B | CCR5 | CTRPNNNTRKSIPLGPGSAFYATDRIGDIRQAYC | 34 |
| HQ644832 | B | CCR5 | CTRPNNNTRRSITIGPGRAFYTTGEIIGDIRQAHC | 35 |
| AF258968 | B | CCR5 | CTRPNNNTRKGIHLGPGRAFYATGGIVGNIRQAHC | 35 |
| JQ779902 | C | CCR5 | CIRPGNNTRKSVRIGPGQTYFSTGEIIGNIRQAHC | 35 |
| HQ644902 | B | CCR5 | CTRPNNNTKKSIHIGPGRAFYTTGDIIGDIRKAHC | 35 |
| DQ516262 | B | CCR5 | CTRPNNNTRKGIHMGPGRAFFTTGAIIGDIRQAHC | 35 |
| DQ002288 | B | CCR5 | CIRPNNNTRKSINIGPGRAFYATGDIIGDIRQAHC | 35 |
| EU272192 | B | CCR5 | CTRPNNNTRKSINLGPGRAWYTTGQIIGDIRQAHC | 35 |
| AF384331 | B | CCR5 | CIRPNNNTRKGIHIGPGRTFYATGEIIGNIRQAHC | 35 |
| KF770449 | C | CCR5 | CTRPGNNTRRSVRIGPGQAFYTTGEIIGDIRAAHC | 35 |
| HQ644993 | B | CCR5 | CIRPNNNTRKSIPIGPGSAFYATGDIIGDIRQAHC | 35 |
| FJ376018 | C | CCR5 | CIRPNNNTRESIRIGPGQTFYATDIIGIRQAHC | 33 |
| EU272255 | B | CCR5 | CTRPNNNTRKRVSLGPGRVWYTTGQIVGDIRQALC | 35 |
| KF770379 | C | CCR5 | CTRPGNNTRKSIRIGPGQTFYATGDIIGDIRQAHC | 35 |
| HQ644857 | B | CCR5 | CIRPGNNTRKSIHIGPGRAFYATGDIIGDIRQAHC | 35 |
| DQ002336 | B | CCR5 | CTRPNNNTRKSIHIAPGRAFYATGAIIGNIRQAYR | 35 |
| FJ998109 | B | CCR5 | CTRPNNNTRRGIHMGPGRAFYATDIIGDIRQAHC | 34 |
| EU744051 | B | CCR5 | CTRPSNNTRKDIHIGPGRAFYATGEIIGDIRQAHC | 35 |
| HM239569 | B | CCR5 | CTRPNNNTRKGINIGPGRAFYATTDIIGDIRQAHC | 35 |
| AJ418508 | B | CCR5 | CTRLNNNTRKSIHMGPGRAFYATGEIIGDIRQAHC | 35 |
| AF541012 | B | CCR5 | CTRPNNNTRKSIPMGPGRAFYATGEIIGDIRQAHC | 35 |
| KC312506 | B | CCR5 | CTRPNNNTRKGIHIGPGRTFYATGEIIGNIRQAHC | 35 |
| AY010893 | B | CCR5 | CTRPNNNTRTSIHMGPGRALYATGDIIGDIRHAYC | 35 |
| AF307756 | C | CCR5 | CTRPNNNTRESIRIGPGQTFYATGDIIGDIRQAHC | 35 |
| JX973174 | C | CCR5 | CTRPGNNTRKSIRIGPGQTFFATNDIIGDIRQAYC | 35 |
| DQ109997 | B | CCR5 | CTRPNNNTRSIHIPGRAFYTGKIIGIRQAHC | 31 |
| DQ002318 | B | CCR5 | CTRPNNNTRKSIHMGPGRAFYTTGEIIGDIRQAYC | 35 |
| HM246198 | B | CCR5 | CTRPNNNTRKSINIGPGRAFYAIGDIIGDIRQAHC | 35 |
| AY010890 | B | CCR5 | CTRPNNNTRTSIHMGPGRAMYATGDIIGNIRQAYC | 35 |
| AY253304 | C | CCR5 | CVRPGNNTRKSVRIGPGQTFYATGDIIGDIRQAHC | 35 |
| U27443 | B | CCR5 | CTRPNNNTRKSITIGPGRAFYTTGQIIGDIRQAYC | 35 |
| AY842810 | B | CCR5 | CTRPNNNTRKSIHMGPGRAMYATGDIIGNIRQAHC | 35 |
| JF507953 | B | CCR5 | CTRPNNNTAKGIHLGPGRAMYATQRIVGDIRRAHC | 35 |
| HM179833 | C | CCR5 | CTRPGTNTRQSMRIGPGQTFYATGDIIGDIRQAHC | 35 |
| DQ061688 | B | CCR5 | CIRPNNNTRKGIHLGPGGALYATGAIIGDIRQAYC | 35 |
| DQ177190 | B | CCR5 | CTRPNNNTRRSINIGPGRAFYATGEIIGDIRQAYC | 35 |
| HM239541 | B | CCR5 | CTRPNNNTRRSINIGPGRAVYATGEIIGDIRVAHC | 35 |
| JN002002 | B | CCR5 | CTRPNNNTRKSIHIAPGRAFYATGEKIGDIRQAHC | 35 |
| FJ653402 | B | CCR5 | CTRPNNNTRKSIPIGPGRAFFATGDIIGDIRQAHC | 35 |
| DQ002167 | B | CCR5 | CTRPNDNTRKSIPMGPGRAFYATGAIIGDIRQAHC | 35 |
| AY510059 | C | CCR5 | CTRPGNNTRKSMRIGPGQAFYTTGDIIGDIRQAHC | 35 |
| DQ516164 | B | CCR5 | CTRPNNNTRRGIHIGPGRAFYAGDVTGDIRQAHC | 34 |
| U08802 | B | CCR5 | CTRPNNNTRKSIPLGPGQAWYTTGQIIGNIRQAHC | 35 |
| EF643684 | B | CCR5 | CIRPNNNTRKSIHIGPGRAFYATGEIIGNIRQAHC | 35 |
| DQ358761 | C | CCR5 | CTRPNNNTRESIRIGPGQTFYATGEIIGDIRQAYC | 35 |
| JQ779182 | C | CCR5 | CTRPGNNTRRSVRIGPGQTFYATGAIIGNIREAHC | 35 |
| KF716494 | B | CCR5 | CIRPNNNTRKGIHIGPGRVFYATGDIIGDIRQAHC | 35 |

| | | | | |
|---|---|---|---|---|
| HQ678290 | B | CCR5 | CIRPGNNTRKSISIGPGRAFYATGGIIGDIRQAHC | 35 |
| JQ777168 | C | CCR5 | CIRPGNNTRRSVRIGPGQTFYATGDIIGDIRKAHC | 35 |
| AF153176 | C | CCR5 | CTRPSNNTRKSVRIGPGQTFYATGDIIGDIRQAHC | 35 |
| HQ377382 | B | CCR5 | CTRPNNNTRKGIHIGPGRAFYTTGDIIGDIRQAHC | 35 |
| DQ382361 | C | CCR5 | CTRPNNNTRESIRIGPGQTFFATGDIIGDIRQAYC | 35 |
| EF657896 | B | CCR5 | CIRPNNNTRKSIPMGPGKAFYATGGIIGNIRQAHC | 35 |
| JQ779231 | C | CCR5 | CIRPGNNTRKSIRIGPGQVFFTATDIIGNIREAHC | 35 |
| EF688449 | B | CCR5 | CTRPNNNTRRSINIGPGRAFYTTGEIIGDIRQAHC | 35 |
| GU945308 | C | CCR5 | CIRPNNNTRKSIRIGPGQTFYATGDIIGDIRQASC | 35 |
| JF508049 | B | CCR5 | CTRPNNNTRKSIHIGPGRAFYTTGEIIGNIRQASC | 35 |
| EU744052 | B | CCR5 | CTRPSNNTRRGIHIGPGRAFYTTGEIIGDIRQAHC | 35 |
| DQ002282 | B | CCR5 | CIRPNNNTRKSINIGPGRAFYATRDIIGDIRQAHC | 35 |
| JQ779091 | C | CCR5 | CTRPGNNTRKSVRIGPGQTFFATGDIIGDIRKAYC | 35 |
| FJ375995 | C | CCR5 | CTRPNNNTRKSIRIGPGQTLFATGAIIGNIRQAHC | 35 |
| DQ002339 | B | CCR5 | CTRPSNNTRKSIHLGLGRAFYATGEIIGDIRQAHC | 35 |
| AY887873 | C | CCR5 | CTRPNNNTRKSIRIGPGQTFYATGDIIGNIRQAHC | 35 |
| HM239567 | B | CCR5 | CTRPNNNTRRSIHIAPGRAFYATGEIIGNIRQAHC | 35 |
| HM239610 | B | CCR5 | CTRPNNNTRRSINIAPGRAFYATGDVIGDIRQAHC | 35 |
| EF600096 | B | CCR5 | CTRLNNNTRKGIHIGPGRAFYATGDIIGDIRQAHC | 35 |
| JN001997 | B | CCR5 | CTRPNNNTRKSIHITPGRAFYATGEIIGDIRQAHC | 35 |
| DQ002224 | B | CCR5 | CTRPSNNTRKSITIGPGKAFYATGEIIGDIRKAHC | 35 |
| HM246205 | B | CCR5 | CTSNNTRKSIHIGPGRAFYATGEIIGNIRQAHC | 33 |
| DQ061699 | B | CCR5 | CTRSNNNTRKGIHMGPGRAFYATGNIIGDIRQAYC | 35 |
| AY010847 | B | CCR5 | CTRPNNDTRKSIHLGPGSAFYAPGDIIGDIRQAHC | 35 |
| AF384326 | B | CCR5 | CTRPNNNTRKSISIGPGRAFYAHGDIVGDIRQAHC | 35 |
| KF770403 | C | CCR5 | CTRPGNNTRKSIRIGPGQTFYATGDIIGDIRKAHC | 35 |
| AF153131 | C | CCR5 | CTRPDNNTRKSMRIGPGQTFYATGDIIGNIRQAHC | 35 |
| HQ678259 | B | CCR5 | CTRPNNNTSQSIHIGPGRAFYATGRIIGDIRQAHC | 35 |
| AY833059 | C | CCR5 | CARPNNNTRKSMRIGPGQTFYATGDIIGDIRQAHC | 35 |
| DQ061483 | B | CCR5 | CTRPHNNTRKSIHIGPRRAFYTTEEIIRNIRQAHC | 35 |
| KC156338 | C | CCR5 | CTRPNTNIRKSMRIGPGQTFYATGDIIGNIRQAHC | 35 |
| KF770308 | C | CCR5 | CTRPDNNTRRSVRIGPGQAFYTNDIIGDIRQAHC | 34 |
| HM239528 | B | CCR5 | CERTNNNTRKSIHIGPGQALYATTNIIGDIRKAHC | 35 |
| HQ644886 | B | CCR5 | CIRPNNNTRKSIHIGPGRAFYATGDIIGNIRQAHC | 35 |
| AF153161 | C | CCR5 | CTRPNNNTRKSVRIGPGQTFYATGEIIGDIRQAHC | 35 |
| AF384248 | B | CCR5 | CTRPNNNTRKGIHIGPGRAFYTTGEIIGDIRKAHC | 35 |
| FJ653136 | B | CCR5 | CTRFNNNTRKSMHIGPGRAFYATGEIIGNIRQASC | 35 |
| JQ779094 | C | CCR5 | CTRPGNNTRKSVRIGPGQTFFATGDIIGDIRQAYC | 35 |
| AM156918 | B | CCR5 | CTRPSNNTRKGIHIGPGRAFYATEEITGDIRQAHC | 35 |
| HM239526 | B | CCR5 | CTRPSNNTRRGIHMAVGRAFYATGQIIGDIRQAHC | 35 |
| EU272330 | B | CCR5 | CTRPNNNTRKSIHLGQGRVWYTTGQIIGDIRQAHC | 35 |
| HQ678271 | B | CCR5 | CIRPNNNTRKSISIGPGRAFVTTGQIIGNIRQAHC | 35 |
| HQ644815 | B | CCR5 | CTRPNNNTRRSINIGPGRAFFATGDIIGDIRQAHC | 35 |
| DQ002098 | B | CCR5 | CTRPSNNTRKSITIGPGRAFYTAEVIGNIRAAYC | 34 |
| HQ644818 | B | CCR5 | CTRPNNNTRKSINIGPGRAFYATGEIIGDIRQAHC | 35 |
| GU945309 | C | CCR5 | CIRPNNNTRKSIRIGPGQTFYATGDVIGDIRQASC | 35 |
| AF384320 | B | CCR5 | CTRPNNNTRKGIHIRPGGAIYATGAITGDIRQAHC | 35 |
| U08780 | B | CCR5 | CTRPNNNTRKSIHLGWGRAFYATGEIVGDIRQAHC | 35 |
| JF507746 | B | CCR5 | CIRPNNNTRKGIHIGPGRTFFATGEIIGDIRRAYC | 35 |
| DQ516264 | B | CCR5 | CTRPNNNTRKGIHMGPGRAFYTTGAIIGDIRQAHC | 35 |
| HQ644938 | B | CCR5 | CTRPNNNTRKSINIGPGRAIYATGEIIGDIRQAHC | 35 |
| EU576867 | B | CCR5 | CTRPNNNTRRGIHMGPGKAYFTGEIIGDIRQAHC | 34 |
| DQ382374 | C | CCR5 | CTRPNNNTRKSMRIGPGQAFYAMGDIIGDIRQAHC | 35 |
| AF082381 | B | CCR5 | CTRPNNNTRKGIHIGPGRAFYATDIIGDIRQAHC | 34 |
| KF770344 | C | CCR5 | CTRPGNNTRRSIRIGPGQAFYATGGIIGDIRQAHC | 35 |
| KC473833 | B | CCR5 | CVRPNNNTRQGIHMGPGRTFYTTGGIIGDIRQAYC | 35 |
| EU744161 | B | CCR5 | CTRPNNNTRRSIHIGPGRAFYATGDIIGNIRQAHC | 35 |
| AF541079 | B | CCR5 | CTRPSNNTRKGIHIGLGRRFYTTEIVGDIRKAYC | 34 |
| FJ653098 | B | CCR5 | CTRPNNNARKSINIGPGRAFYAATDIIGDIRQAHC | 35 |
| AF153141 | C | CCR5 | CVRPNNNTRKSIRIGPGQTFYANDIIGDIRQAHC | 34 |
| AF153156 | C | CCR5 | CVRPSNNTRKSVRIGPGQTFYATKDIIGDIRQAHC | 35 |
| AY010895 | B | CCR5 | CTRPNNNTGKSMPMGPGRAFYSTGEIMGHLRKTYS | 35 |
| KC312574 | B | CCR5 | CERPNNNTRKSIRIGPGSTFYATGEIIGNIRQAHC | 35 |
| HQ708048 | C | CCR5 | CTRPNNNTRKSVRIGPGQTFYATGGIIGDIRRAYC | 35 |
| DQ516285 | B | CCR5 | CTRPNNNTRKSIHMGPGRAFYATGEIIGNIRQAHC | 35 |
| EU272261 | B | CCR5 | CTRPNNNTRKSINLGPGRAWYATGQIIGDIRQAHC | 35 |
| EU272281 | B | CCR5 | CTRPNNNTRKSIPLGPGKAWYTTGDIIGDIRQAHC | 35 |
| DQ061481 | B | CCR5 | CTRPNNNTRKSISIGPGRAFYTTGEIIGEIRQAYC | 35 |
| AY510057 | C | CCR5 | CTRPNNNTRKSVRIGPGQAFYATGDIIGNIRQAYC | 35 |
| KF384804 | B | CCR5 | CTRPGNNTSKGIHIGPGRALHVTRTRGKIIGDIRQA | 38 |

| | | | | |
|---|---|---|---|---|
| HQ377487 | B | CCR5 | CTRPGNNTRKSIPIGPGRAFYATGDVIGDVRKAHC | 35 |
| HM179824 | C | CCR5 | CTRPGNNTRQSMRIGPGQTFYATGNIIGDIRQAHC | 35 |
| EU786680 | B | CCR5 | CTRPNNNTRKSIHIGPGRALYAQDIIGDIRQAHC | 34 |
| EF657898 | B | CCR5 | CIRPNNNTRKSIPMGPGKAFYATGSIIGNIRQAHC | 35 |
| HM179924 | C | CCR5 | CTRPNNNTRKSIRIGPGQTFYATGDIIGDIRQAYC | 35 |
| M26727 | B | CCR5 | CTRPNNNTRNRISIGPGRAFHTTKQIIGDIRQAHC | 35 |
| AF543910 | C | CCR5 | CIRPNNNTRKSIRIGPGQTFYATNDIIGDIRQAHC | 35 |
| FJ376028 | C | CCR5 | CTRPGNNTRESIRIGPGQTFYATGDIIGNTRQAYC | 35 |
| EU577210 | B | CCR5 | CTRPNNNTRKSVHIGPGKVFYAGEIIGDIRQAHC | 34 |
| EF688438 | B | CCR5 | CTRPNNNTRKAIHLGPGRAFYTTGGIIGDIRQAHC | 35 |
| DQ904344 | C | CCR5 | CTRPNNNTRKSVRIGPGQTFYATGDIIGDIREAHC | 35 |
| AJ418520 | B | CCR5 | CTRPDNNTRKSIHIGPGRAFYATGDIIGNIRQAHC | 35 |
| AY887878 | C | CCR5 | CTRPNNNTRQSMRIGPGQTFYATGDIIGDIRQAHC | 35 |
| EU272290 | B | CCR5 | CTRPSNNTRKSINLGPGRAWYTTGQIIGDIRQAHC | 35 |
| EU908220 | C | CCR5 | CTRPNNNTRQSIRIGPGQVFYATGAIIGDIRQAYC | 35 |
| AY842792 | B | CCR5 | CTRPNNNTRKSIHMGPGSVWYATGEIIGDIRQAHC | 35 |
| EU744005 | B | CCR5 | CTRPNNNTRKSISIGPGRAFYATGDIVGDIRKAHC | 35 |
| AY124978 | B | CCR5 | CTRPSNNTRKSINIGAGRAIYATGDIIGDIRQAYC | 35 |
| KC312586 | B | CCR5 | CVRPNNNTRKSIHLGPGSAFYATGKIIGNIRQAHC | 35 |
| HQ678282 | B | CCR5 | CTRLNNNTRKSIRIGPGSTFYTSNIIGDIRQAYC | 34 |
| DQ002340 | B | CCR5 | CTRPNNNTRKSIHIAPGRAFYATGDIIGNIRQAYC | 35 |
| DQ002281 | B | CCR5 | CTRPNNNTKRSVNIVPGRAIYTTDIIGDIRQAHC | 34 |
| KC156120 | C | CCR5 | CTRPNNNTRKSIRIGPGQAFFATGDIIGDIREAHC | 35 |
| U52953 | C | CCR5 | CTRPNNNTRKSIRIGPGQAFYATGEIIGDIRQAHC | 35 |
| AY842817 | B | CCR5 | CTRPNNNTRKSIPMGPGRAMYATGDIIGDIRKANC | 35 |
| AF258977 | B | CCR5 | CTRPNNNTRKGIHLGPGRAFYATGGIVGDIRQAHC | 35 |
| KC156437 | C | CCR5 | CTRPNNNTRKSVRIGPGQTFYATGEIIGDIREAYC | 35 |
| DQ435683 | C | CCR5 | CIRPGNNTRRSIRIGPGQAFYAMGDIIGNIREAHC | 35 |
| EF688450 | B | CCR5 | CTRPNNNTRKSIHIGPGRAFYGTDIIGDIRQAHC | 34 |
| HQ708025 | C | CCR5 | CTRPNNNTRKSVRIGPGQTFYATGDIIGDIRQAYC | 35 |
| JF896853 | B | CCR5 | CTRPNNNTRKSIHMGPGKTFFATGDIIGDIRQAHC | 35 |
| DQ516304 | B | CCR5 | CTRPNNNTRKSIHIGPGRALYTTGEIIGNIRQAHC | 35 |
| EF688457 | B | CCR5 | CTRPNNNTRKSIPMGPGRAFYTTGEIIGDIRQAHC | 35 |
| EF643662 | B | CCR5 | CTRPNNNTRRSISIGPGRAFYATGEVIGDIRQAHC | 35 |
| EU744065 | B | CCR5 | CTRPNNNTRKSIHMGPGKTFYATGDIIGDIRQAHC | 35 |
| JF507805 | B | CCR5 | CMRPGNNTRKSIAIGPGRAFYAGEIIGDIRKAHC | 34 |
| AJ418534 | B | CCR5 | CARPHNNTRKSIHIGPGRAFYATGQITGDIRQAYC | 35 |
| AF541039 | B | CCR5 | CTRPSNNTRKSIPMGPGKAFYATGDIIGDIRQAHC | 35 |
| AF384294 | B | CCR5 | CTRPNNNTRKSIHIGPGRAVYATGQIIGDIRQAHC | 35 |
| AY842805 | B | CCR5 | CTRPNNNTRRSIHMGPGSVLYATGEIIGDIRQAHC | 35 |
| DQ002149 | B | CCR5 | CTRPGNNTRRGIHIGPGRAFYTTGEIIGDIRQAHC | 35 |
| HM239579 | B | CCR5 | CTRPNNNTRKSIIGPGRAYFATGIIGDIRKAHC | 33 |
| AF391245 | C | CCR5 | CTRPNNNTRRSIRIGPGQAFYTNDIIGDIRQAHC | 34 |
| AY887865 | C | CCR5 | CIRPGINKRKRIRIGLRYAFFATDNRIGKAHC | 32 |
| FJ670531 | B | CCR5 | CIRPNNNTRKSIHMGPGRAFYATGDVIGDIRKAYC | 35 |
| EU744008 | B | CCR5 | CTRPNNNTRKSISIGPGRAFYATGDIIGDIRKAHC | 35 |
| DQ869022 | B | CCR5 | CTRPGNNTRKSIHIAPGRAFYATGDIIGDIRQAHC | 35 |
| DQ002231 | B | CCR5 | CTRPSNNTRKSIPIGSGRAFYATGDIIGDIRKAHC | 35 |
| HM239613 | B | CCR5 | CTRPNNNTRKGITMGPGKAFYVTGGIVGDIRQAHC | 35 |
| AF199036 | B | CCR5 | CTRPNNNTRKSIHIGPGRALYATGDIVGDIRQAHC | 35 |
| DQ904347 | C | CCR5 | CTRPNNNTRKSIRIGPGQTFYATGDIIGNIREAHC | 35 |
| AY842807 | B | CCR5 | CTRPNNNTRKSIHMGPGSVLYATGEIIGDIRQAHC | 35 |
| DQ002264 | B | CCR5 | CTRPNNNTRKSIPIGPGRAFYTTGEIIDIRQAHC | 34 |
| HQ644868 | B | CCR5 | CTRPGNNTRKSIPMGPGRAFYATGDIIGDIRKAHC | 35 |
| U08455 | C | CCR5 | CTRPNNNTRKSVRIGPGQTFYATGAIIGDIRQAHC | 35 |
| AY010794 | B | CCR5 | CIRPNNNTRKSIQMGPGKAFYATGDIIGDIRQAYC | 35 |
| HQ644921 | B | CCR5 | CTRPNNNTRKSIPIGPGRAFYTTGDIIGDIRQAHC | 35 |
| JN002053 | B | CCR5 | CTRPNNNTRKSIHMGPGKAFYTTGVIGDIRQAHC | 34 |
| U45299 | B | CCR5 | CTRPNNNTRRGVHIGPRRGFYTTEIIGNIRQAHC | 34 |
| U23487 | B | CCR5 | CTRPSNNSRKSIYIGPGRRFHVTRAVTGDIRQAHC | 35 |
| FJ653358 | B | CCR5 | CTRPNNNTRRGIHIGPGRAFYATDIIGDIRQAHC | 34 |
| HM246193 | B | CCR5 | CIRPNNNTRKSIHIAPGRAFYATGNIIGDIRKAYC | 35 |
| U08712 | B | CCR5 | CTRPNNNTRKSIHMGWGRTFYATGEIIGDIRQAHC | 35 |
| HM239588 | B | CCR5 | CTRPNNNTRKSIHIGPGRAFYATGDIIGEIRQAYC | 35 |
| DQ061678 | B | CCR5 | CIRPNNNTRKGIHLGPGGALYATGGIIGDIRRAYC | 35 |
| DQ002026 | B | CCR5 | CIRPHNNTRKSIPIGPGRAFYTTGEIIGDIRQAHC | 35 |
| AF355727 | B | CCR5 | CTRPNNNTRKSIPLGPGRAFYATGDIIGNIRKAHC | 35 |
| EU575305 | B | CCR5 | CTRPGNNTRKSITIGPGRAFYATGDIIGDIRQAHC | 35 |
| U79719 | B | CCR5 | CTRPNNNTRKGIHIGPGAAFYATGEIIGDIRQAHC | 35 |
| AF310127 | B | CCR5 | CTRPNNNTRKSIHLGAGRALYTGEIIGDIRQAHC | 34 |

| | | | | |
|---|---|---|---|---|
| AY887886 | C | CCR5 | CIRPGNNTRKSVRIGPGQTIYATGDIIGDIRQAHC | 35 |
| AF022258 | B | CCR5 | CTRPNNNTRKSISIGPGRAFYTTGEIIGDIRQAHC | 35 |
| AJ418505 | B | CCR5 | CTRPNNNTRKSIPIGPGRAFYTTGEIIGDIRKAHC | 35 |
| EF657905 | B | CCR5 | CIRPNNNTRKSIPMGPGRAFYATGDIIGNIRQAHC | 35 |
| KC156318 | C | CCR5 | CTRPNDNIRKSMRIGPGQTFYATGDIIGNIRQAHC | 35 |
| AF153172 | C | CCR5 | CVRPNNNTRKSIRIGPGQTFYATNGIIGDIREAHC | 35 |
| HM239595 | B | CCR5 | CTRPNNNTRRSIRIGPGSYFATGDIIGDIRKAYC | 34 |
| AJ418480 | B | CCR5 | CTRPNNNTRKSIPIGPGRALYATGDIIGEIRQAFC | 35 |
| KF770391 | C | CCR5 | CIRPGNNTRKSMRIGPGQTFYATGDIIGDIRKAHC | 35 |
| HM246188 | B | CCR5 | CTRPNNNTRKSIHIVPGGAFYATGDIIGDIRQAHC | 35 |
| EU744075 | B | CCR5 | CIRYNNNTRKSIPLGPGRAFYATGDIIGNIRQAQC | 35 |
| GU945313 | C | CCR5 | CTRPGNNTRKSIRIGPGQTFYATGGIIGNIRQAHC | 35 |
| HM246190 | B | CCR5 | CSRPNNNTKSIPIGPGAFYATGDIIGDIRQAHC | 33 |
| AF153164 | C | CCR5 | CIRPGNNTRKSIRIGPGQAFYATGDIIGDIRQAHC | 35 |
| DQ061463 | B | CCR5 | CTRPNNNTRKSIHIGPGRAFYTTGEIIGDIRKAHC | 35 |
| AB553911 | B | CCR5 | CERPNNNTRRSIQIGPGRAWFEAEDIIGDIRKAHC | 35 |
| DQ516149 | B | CCR5 | CTRPNNNTRRDIHIGPGRAFYATGGIIGNIRQAHC | 35 |
| DQ002342 | B | CCR5 | CTRPNNNTRKSIHIGPGSAFYATGDIIGNIRQAYC | 35 |
| AF355726 | B | CCR5 | CTRPSNNTRKSIHLGPGRAFYATGDIIGNIRKAHC | 35 |
| DQ061452 | B | CCR5 | CTRPNNNTRKSIHIGPGRAFYTTGEIMGDIRKAHC | 35 |
| AF258994 | B | CCR5 | CTRPNNNTRKGIHLGPGRAFYATGEIVGDIRKAHC | 35 |
| EU272217 | B | CCR5 | CIRPNNNTRKSIPLGPGKAWYTTGQIIGDIRQAHC | 35 |
| HQ708068 | C | CCR5 | CIRPSNNTRTSMRIGPGQTFFATGDVTGDIRQAYC | 35 |
| EU744015 | B | CCR5 | CTRPNNNTRKSIHIGPGRAFYTTGEIIGEIRQAHC | 35 |
| GU455522 | B | CCR5 | CIRPNNNTRKSIPMGPGKAFYTTGDIIGDIRQAHC | 35 |
| JQ779893 | C | CCR5 | CIRPGNNTRKSIRIGPGQTYFSTGEIIGNIRQAHC | 35 |
| KF766537 | C | CCR5 | CIRPNNNTRKSIRIGPGQTFYATGEIIGDIRQAHC | 35 |
| KF770295 | C | CCR5 | CIRPNNNTRKSVRIGPGQTFYATGEIIGDIRKAHC | 35 |
| DQ002148 | B | CCR5 | CTRPGNNTRRGIHIGPGRAFYTTGEIIGNIRQAHC | 35 |
| KC156326 | C | CCR5 | CNRPHNNTRKSMRIGPGQAFYATGDITGDIRKAHC | 35 |
| EU744078 | B | CCR5 | CTRYNNNTRKSIHLGPGRAFYATGDIIGDIRQAQC | 35 |
| EU293450 | C | CCR5 | CTRPSNNTRKSVRIGPGQAFFATGEIIGDIRQAHC | 35 |
| EU272222 | B | CCR5 | CIRPNNNTRKSIPIGPGKAWYTTGQIIGDIRQAHC | 35 |
| DQ516241 | B | CCR5 | CTRPNNNTRKGIHIGPGRAFYTTGEIIGDIRQAYR | 35 |
| DQ061524 | B | CCR5 | CTRPNNNTRRSIHIAPGSAFYATGDIIGDIRQAHC | 35 |
| HQ644904 | B | CCR5 | CTRPNNNTRKSIPIGPGRAFYTTGDIIGDIRKAHC | 35 |
| JN002050 | B | CCR5 | CTRPDNNTRKGIHLGPGGTFFATGAIIGDIRQAHC | 35 |
| EU576298 | B | CCR5 | CTRPNNNTRKGIHIGPGKTFYTTGGIIGDIRQAYC | 35 |
| DQ869026 | B | CCR5 | CTRPNNNTRKSIHIAPGRPFYATGDIIGDIRQAHC | 35 |
| AF153138 | C | CCR5 | CTRPGNNTRKSVRIGPGQTFYATGEIIGNIRQAHC | 35 |
| DQ411850 | C | CCR5 | CTRPNNNTRKSIRIGPGQTVYATNDIIGDIRQAHC | 35 |
| HQ644927 | B | CCR5 | CMRPNNNTRKSIPIGPGRAFYTTGDIIGDIRQAHC | 35 |
| KC473843 | B | CCR5 | CTRPNNNTRRGIQMGPGRAVYATGDIIGDIRQAHC | 35 |
| EF688452 | B | CCR5 | CTRPNDNTRKGINIGPGRAFYATGEIIGDIRQAHC | 35 |
| AF254783 | C | CCR5 | CTRPGNNTRKSIRIGPGQAFFATGAIIGDIRKAYC | 35 |
| HM239529 | B | CCR5 | CTRPSNNTRKSITIGPGRAFYTTGQIIGNIRQAHC | 35 |
| KC473841 | B | CCR5 | CTRHSNNTRRSIRIGPGQAFYATGEIIGDIRKAHC | 35 |
| FJ653126 | B | CCR5 | CTRFNNNTRKSIHIGPGRAFYATGDIIGNIRQASC | 35 |
| KC156124 | C | CCR5 | CTRPNNNTRTSIRIGPGQTFYATGDIIGDIRQAYC | 35 |
| HM246228 | B | CCR5 | CTRPNNNTRKSIHIAPGRAFYATGDIIGDIRQAYC | 35 |
| AY887879 | C | CCR5 | CMRPNNNTRKSVRIGPGQTFYATGDIIGNIRQAHC | 35 |
| HQ644887 | B | CCR5 | CTRPNNNTRKSIHIGPGRAFYTTGEIIGKIRQAHC | 35 |
| DQ061752 | B | CCR5 | CTRPNNNTRRSITLGPGRAYYATGDIIGDIRQAHC | 35 |
| JF508128 | B | CCR5 | CTRPNNNTRRSIYIGPGRTFYATGEIIGDIRQAHC | 35 |
| HM239566 | B | CCR5 | CTRPSNNTRKGIHMAVGRAFYTTGQIIGNIRQAHC | 35 |
| DQ388517 | C | CCR5 | CTRPNNNTRKSIRIGPGQSFYATGEIVGNIREAHC | 35 |
| AF153181 | C | CCR5 | CIRPSNNTRRSVRIGPGQTFYATGEITGDIRQAHC | 35 |
| JF508018 | B | CCR5 | CTRPNNNRRKSINIGPGRAFYTTGAIIGDIRQAHC | 35 |
| AY010831 | B | CCR5 | CTRPNSNTRKGIHIGPGSAIYATGDIIGDIRQAHC | 35 |
| JX140654 | B | CCR5 | CTRPGNNTRRSITMGPGKVFYTNEIIGNIRRAYC | 34 |
| JF507854 | B | CCR5 | CTRPGNNTRRSINIGPGRAFYTGEIIGDIRQAHC | 34 |
| HQ644992 | B | CCR5 | CTRPNNNTRKSIPIGPGRAFYATGEIIGDIRQAHC | 35 |
| HM179723 | C | CCR5 | CTRPNNNTRKSMRIGPGQTFYATGEIIGNIREAHC | 35 |
| HQ708049 | C | CCR5 | CTRPNNNTRKSIRIGPGQAFYATGAVTGDIRKAYC | 35 |
| AY736825 | C | CCR5 | CTRPGDNRRRSVRIGPGQTFYATGEIIGNIREAHC | 35 |
| AY255825 | C | CCR5 | CTRPNNNTRRSIRIGPGQTFYATGEIIGNIREAHC | 35 |
| KC156396 | C | CCR5 | CIRPGNNTRRSVRIGPGQTYFSTGEIIGNIRQAHC | 35 |
| JF507916 | B | CCR5 | CTRPNNNTRKSISIGPGRAFYATGEIIGDIRQAHC | 35 |
| AF384291 | B | CCR5 | CTRPNNNTRKSIPIGPGRAFYARGDIIGDIRQAHC | 35 |
| EU272220 | B | CCR5 | CIKPNNNTRKSIPIGPGKAWYTTGQIIRDIRRAHC | 35 |

| | | | | |
|---|---|---|---|---|
| JN002015 | B | CCR5 | CTRPNNNTRKGIHIGPGRAFYATGDIVGDIKQAHC | 35 |
| HQ645006 | B | CCR5 | CTRPNNNTRRSISIGPGRAFYATGDIIGDIRRAQC | 35 |
| DQ235634 | C | CCR5 | CTRPNNNTRKSVRIGPGQAFYATNDIIGDIRQAHC | 35 |
| AF259010 | B | CCR5 | CTRPNNNTRKGIHMGPGRAFYATGEIVGDIRKAHC | 35 |
| EU161645 | C | CCR5 | CTRPNNNTRTSTRIGPGQAFYATGDIIGDIRQAHC | 35 |
| DQ177198 | B | CCR5 | CTRPNNNTRKSINIGPGRAFYSTGAIIGNIRQAHC | 35 |
| EF117268 | C | CCR5 | CTRPNNNTRKSIRIGPGQTFYATGDIIGNIRQAYC | 35 |
| HQ699981 | C | CCR5 | CTRPNNNTRKSVRIGPGQAFYATGDIIGNIREAHC | 35 |
| EF643664 | B | CCR5 | CTRPNNNTRKGISIGPGRAFYTTGEVIGDIRQAHC | 35 |
| KF770347 | C | CCR5 | CTRPNNNTRRSIRIGPGQAIYTTGGIIGDIRQAHC | 35 |
| AF021502 | B | CCR5 | CTRPNNNTRKSIPIGPGRAFYTTGQIIGDIRQAYC | 35 |
| AF310109 | B | CCR5 | CTRPNNNTRKSIHIGAGKALYTGEIIGDIRQAHC | 34 |
| AF153173 | C | CCR5 | CTRPNNNTRKSIRIGPGQTFYANDIIGDIRQAHC | 34 |
| JN002045 | B | CCR5 | CTRPNNNTRKSIPIGPGGAFYATGDIIGDIRQAHC | 35 |
| DQ110002 | B | CCR5 | CIRPNNNTRKSIPMGPGRAFYATGDIIGDIRQAHC | 35 |
| JN002019 | B | CCR5 | CTRPNNNTRKGIHIGPGRAFYATGDIVGDMRQAHC | 35 |
| HQ644936 | B | CCR5 | CTRPNNNTRKSINIGPGRALYTTGEIIGDIRQAHC | 35 |
| JQ779245 | C | CCR5 | CIRPGNNTRKSIRIGPGQVLFATTDIIGNIREAHC | 35 |
| DQ435682 | C | CCR5 | CRRPNNNTRKSIRIGPGQAFYATNDIIGDIRQAHC | 35 |
| AF258959 | B | CCR5 | CIRPSNNTRTSIHLGPGQAVYATGEIIGNIRQAHC | 35 |
| EU786676 | B | CCR5 | CTRPNNNTRKSITIGPGRAFYATGDVIGDIRQAHC | 35 |
| DQ382376 | C | CCR5 | CTRPGNNTRKSVRIGPGQVFYATGDIIGDIRQAHC | 35 |
| AF022281 | B | CCR5 | CTRPNNNTRKSISIGPGRAFYATGDIIGDIRQAHC | 35 |
| U08699 | B | CCR5 | CTRPNNSTRKSIHMGWGRAFYATGEIIGNIRQAHC | 35 |
| KC156458 | C | CCR5 | CTRPNNNTRKSVRIGPGQTFYATGEIIGNIREAYC | 35 |
| DQ002312 | B | CCR5 | CTRPNNNTRKSINIGPGRAWYTTGDIIGDIRKAHC | 35 |
| HQ644915 | B | CCR5 | CTRPNNNTRKSIHIGPGRAFYTTEIIGDIRKAYC | 34 |
| HQ644803 | B | CCR5 | CTRPNNNTRRSINIGPGRAFFATGEIIGDIRQAHC | 35 |
| EU786677 | B | CCR5 | CTRPNNNTRKSIRIGPGQAFYTTGEIIGDIRQAHC | 35 |
| AF259020 | B | CCR5 | CTRPGNNTRKSIHIGPGRAFYTTGQIIGKIRQAHC | 35 |
| HQ708020 | C | CCR5 | CIRPDNNTRKSIRIGPGQTFYATGDIIGNIRKAYC | 35 |
| HQ644793 | B | CCR5 | CTRPNNNTRRSIHIGPGRAFYTGEIIGNIRQAHC | 34 |
| JX140655 | B | CCR5 | CTRPNNNTRKGIHIGPGKTFFATDIIGDIRQAHC | 34 |
| KF770439 | C | CCR5 | CIRPGNNTSRSIRIGPGQTFYATGRIVGDIRQAHC | 35 |
| HM239515 | B | CCR5 | CTRPSNNTRKSIHMGPGRVLFATGEIIGNIRQAHC | 35 |
| DQ222217 | B | CCR5 | CTRPNNNTRKSIHVGPGKAIYTTEIIGNIRQAHC | 34 |
| U08450 | B | CCR5 | CIRPNNNTRRSIHMGLGRAFYATGDIIGDIRQAHC | 35 |
| HQ644979 | B | CCR5 | CTRPNNNTRRSINIGPGRAIYATGEIIGDIRQAHC | 35 |
| HQ644805 | B | CCR5 | CIRPNNNTRRSINIGPGRAIFTTGEIIGDIRQAHC | 35 |
| DQ061742 | B | CCR5 | CTRPNNNTRRSISLGPGRAYYATGDIIGDIRQAHC | 35 |
| JX140668 | C | CCR5 | CIRPNNNTRKSIRIGPGQTFYANNIIGDIRQAYC | 34 |
| AY887867 | C | CCR5 | CTRPNNNTRRSVRIGPGQAFYGTDIIGDIRQAYC | 34 |
| EU272262 | B | CCR5 | CTRPNNNTRKSINLGPGAWYATGQTIGDIRQAHC | 34 |
| HM239577 | B | CCR5 | CTRPNNNTRRSIHIAPGKTFYATGAIVGDIRQAYC | 35 |
| JQ779230 | C | CCR5 | CTRPNNNTRKSIRIGPGQAFFATTGIIGNIRQASC | 35 |
| HQ644882 | B | CCR5 | CTRPNNNTRKSINIGPGRAFYATGDIIGNIRQAHC | 35 |
| DQ002137 | B | CCR5 | CTRPNNNTRKSIHIRPGKAFYATGDIIGDIRQAHC | 35 |
| DQ061685 | B | CCR5 | CIRPNNNTRKGIHLGPGGALYATGGIIGDIRQAYC | 35 |
| JF507888 | B | CCR5 | CTRPSNNTRKSIHIGPGRALYATDIIGDIRKAYC | 34 |
| EF657900 | B | CCR5 | CIRPNNNTRKSIPMGPGKAFYATGDIIGNIRQAHC | 35 |
| AY010808 | B | CCR5 | CTRPNNNTRKGILLGPGSAFYATGDIIGDIRQAHC | 35 |
| AY887866 | C | CCR5 | CTRPGNNTRKSMRIGPGQTFYATGDIIGDIRQAHC | 35 |
| AY010886 | B | CCR5 | CTRPNNNTRTRIHMGPGRAMYATGDIIGNIRQAYC | 35 |
| DQ235620 | C | CCR5 | CTRPNNNTRKSMRIGPGQTFFATGEIIGDIRQAHC | 35 |
| EU578364 | B | CCR5 | CTRPNNNTRKSITIGPGSVFYTGEIIGDIRRAHC | 34 |
| AF391231 | C | CCR5 | CTRPNNNTRKSVRIGPGQAFYATNDVIGNIRQAHC | 35 |
| DQ061817 | B | CCR5 | CTRPNNNTRKGIHMGSGKVFYATGQIIGDIGQAHC | 35 |
| AF384312 | B | CCR5 | CTRPYEKKKSRRIRIGPGRTFHTTGSIGDIRRAHC | 35 |
| HQ645010 | B | CCR5 | CTRPNNNTRKSIPIGPGRAIYTTGQIIGDIRQAHC | 35 |
| EU575924 | B | CCR5 | CTRPGNNTRKSIHIAPGRTFYATGEIIGDIRRAHC | 35 |
| AF384274 | B | CCR5 | CTRPNNNTRKSIHMGPGGSFYATGDIIGNIRQAHC | 35 |
| GU945317 | C | CCR5 | CIRPNNNTRQSIRIGPGQVFYATGDIIGDIRQAYC | 35 |
| DQ061561 | B | CCR5 | CVRPNNNTRKSIPIGPGRAFYTTGEIIGDIRQAHC | 35 |
| DQ382380 | C | CCR5 | CTRPNNNTRTSVRIGPGQAFYATNGIIGDIRQAHC | 35 |
| DQ002203 | B | CCR5 | CTRPSNNTRKSIPIGPGRAFYATGEIIGDIRQAHC | 35 |
| HQ644926 | B | CCR5 | CTRPNNNTRKSIHIGPGKALYTTEIIGNIRQAHC | 34 |
| AY669741 | C | CCR5 | CTRPNNNTRKSMRIGPGQTFYATGDIIGDIRQAYC | 35 |
| AJ888842 | B | CCR5 | CTRPNNNTRKSITIGPGRAFYTTGEIIGDIRKAHC | 35 |
| AY835434 | B | CCR5 | CTRPNNNTRKSIHMGPGAAFYARGEVIGDIRQAHC | 35 |
| AY043175 | C | CCR5 | CTRPNNNTRKSVRIGPGQTFYATGEIIGDIREAHC | 35 |

| | | | | |
|---|---|---|---|---|
| JF507936 | B | CCR5 | CTRPNNNTAKGIHIGPGRAMYATERIVGNIRRAHC | 35 |
| EF688446 | B | CCR5 | CTRPNNNTRKSIRIGPGSAFYATGDIIGDIRQAHC | 35 |
| AM156920 | B | CCR5 | CTRPNNNTRRSIHIGPGRAFYTRDIIGNIRQAHC | 34 |
| DQ002165 | B | CCR5 | CTRPNNNTRKSISMGPGRAFYATGAIIGDIRQAHC | 35 |
| DQ061780 | B | CCR5 | CTRPNNNTRKGIHIGPGRAFYAAGEIIGNIRQAHC | 35 |
| JQ779243 | C | CCR5 | CIRPGNNTRKSIRIGPGQVFFATDIIGDIRKAHC | 34 |
| AJ418484 | B | CCR5 | CTRPNNNTRKGIHIGPGRALYATGDIIGKIRQAHC | 35 |
| EF579976 | B | CCR5 | CERPGNNTSKGIHIGPGRAFYATVNIIGDIRKAHC | 35 |
| DQ235623 | C | CCR5 | CTRPNNNTRKSVRIGPGQTFYATGDIIGNIRQAHC | 35 |
| JF896841 | B | CCR5 | CTRPNNNTRKSIPMGPGKAFYATGIIGDIRQAHC | 34 |
| JF507878 | B | CCR5 | CTRPSNNTRQGIHIGPGRAIYTTDIIGDIRKAYC | 34 |
| HM246187 | B | CCR5 | CTRPNNNTRKSISIGPGRAFYTTGDIIGNIRQAYC | 35 |
| HQ678292 | B | CCR5 | CTRPNNNTRKSIRIGRGRAFYAPGEIIGDIRKAYC | 35 |
| AF021523 | B | CCR5 | CTRPNNNTRKSITIGPGRAFYTTGQIIGDIRKAYC | 35 |
| KF770371 | C | CCR5 | CIRPGNNTRRSVRIGPGQTFYATGDIIGDIRQAHC | 35 |
| EU575170 | B | CCR5 | CVRPNNNTRRSITIGPGRAFYTGEIIGNIRKAYC | 34 |
| JN687739 | B | CCR5 | CTRPSDNTRKSIHMGWGRAFYATGEITGDIRQAHC | 35 |
| HM215421 | C | CCR5 | CTRPNNNTRKSIRIGPGQTFYATGEIIGDIRQAHC | 35 |
| DQ869025 | B | CCR5 | CTRPNNNTRKGIHIGPGRAFYATGEIIGNIRQAYC | 35 |
| EU272207 | B | CCR5 | CTRPNNNTRKSIHIGPGRAWYATGEIIGDIRQAHC | 35 |
| HQ708021 | C | CCR5 | CIRPNNNTRKSVRIGPGQTFYATGDIIGDIRQAYC | 35 |
| AY842793 | B | CCR5 | CTRPNNNTRKSIHMGPGRVWYTTGGIIGDIRQAHC | 35 |
| DQ002156 | B | CCR5 | CTRPNNNTRKSIPMGPGRAFYATGAIIGDIRQAHC | 35 |
| DQ061496 | B | CCR5 | CTRPSNNTRKSIHIGPGRAFYTTGEIIGNIRQAHC | 35 |
| AF153139 | C | CCR5 | CVRPNNNTRKSVRIGPGQTFYATGDIIGDIRQAHC | 35 |
| DQ235641 | C | CCR5 | CIRPNNNTRKSIRIGPGQAFYATGDIIGDIRKAYC | 35 |
| HQ644922 | B | CCR5 | CIRPNNNTRKSIHIGPGRAFYTTEIIGNIRKAYC | 34 |
| HM368253 | B | CCR5 | CTRPNNNTRKSIHIQPGRAFYATDIIGDIRQAHC | 34 |
| AF384251 | B | CCR5 | CTRPNNNTRKSIHMGPGRAFYTTGNIIGDIRKAHC | 35 |
| FJ375982 | C | CCR5 | CTRPNNNTRKSVGMGPGRAIYATGDIIGDIRQAHC | 35 |
| JF507901 | B | CCR5 | CTRPSNNTRQGIHIGPGRAIYTTNIIGDIRKAYC | 34 |
| FJ376013 | C | CCR5 | CTRNNNTRKSIRIGPGQYATGDIIGDIRQAHC | 32 |
| FJ853622 | B | CCR5 | CTRPNNNTRKSIHLGAGKAIYTTGAIIGNIRQAHC | 35 |
| EU744025 | B | CCR5 | CTRPNNNTRKSIHIGPGRALYATGEIIGNIRQAHC | 35 |
| AF082354 | B | CCR5 | CTRPNNNTRRSINIGPGRAFYTTGDIIGDIRQAHC | 35 |
| DQ002041 | B | CCR5 | CIRPHNNTRKSIHIGPGRAFYTTGEIIGDIRQAHC | 35 |
| KF770329 | C | CCR5 | CTRPNNNTRKSVRIGPGQTFYATGEIIGDIRRAHC | 35 |
| HQ644855 | B | CCR5 | CTRPGNNTRKSINIGPGRAIYATGDIIGNIRQAHC | 35 |
| EU575457 | B | CCR5 | CTRPSNNTRRSINIGPGKAFYATGEIIGNIRQAHC | 35 |
| HQ377469 | B | CCR5 | CIRPNNNTRKSISMGPGRAFFATGEIIGNIRQAHC | 35 |
| AY124975 | B | CCR5 | CTRPNNNTRKGIHMGPGGALYATGAIIGNIRQAHC | 35 |
| AY158533 | C | CCR5 | CTRPNNNTRKSMRIGPGQTFFATGDIIGNIRQAHC | 35 |
| EF643657 | B | CCR5 | CTRLNNNTRRSINIGPGRAWYTTGEVIGDIRKAHC | 35 |
| DQ002228 | B | CCR5 | CTRPSNNTRRSITIGPGRAFYATGEIIGDIRKAHC | 35 |
| HM239594 | B | CCR5 | CTRPNNNTRKSIPIGPGSAFYATGDIIGDIRQAHC | 35 |
| HM246197 | B | CCR5 | CTRLNNNTRKSIHIGPGQAFYATGAIIGIRQAHC | 34 |
| FJ653220 | B | CCR5 | CTRPGNNTSKSIHIGPGRAFDATKTITGDIRQAHC | 35 |
| EF117265 | C | CCR5 | CTRPNNNTRKSIRIGPGQTFYATGEIIGNIRQAHC | 35 |
| JN002044 | B | CCR5 | CTRPNNNTRKSIPMGPGKAFYATGDIIGDMRQAHC | 35 |
| AF258982 | B | CCR5 | CTRPNNNTRKGIHLGPGRAFYATGEIIGNIRQAYC | 35 |
| DQ109995 | B | CCR5 | CTRPNNNTRKSINIGPGRAWYTTGQIIGDIRQAHC | 35 |
| HM179818 | C | CCR5 | CTRPGNNTRQSMKIGPGQTFYATGDIIRDIRQAHC | 35 |
| HM239558 | B | CCR5 | CTRPNNNTRKSIHIGPGRAFYATGEIGDIRQAHC | 34 |
| AF310112 | B | CCR5 | CTRPNNNTRKSIHIGPGKALYTGEIIGDIRQAHC | 34 |
| AY426112 | B | CCR5 | CTRPNNNTRKSIHIGPGRALYTTGKIIGDIRQAHC | 35 |
| DQ869024 | B | CCR5 | CTRPNNNTRKGIHMGPGKTLYATGEIIGDIRQAHC | 35 |
| DQ235633 | C | CCR5 | CTRPNNNTRQSIRIGPGQAFFAAKDIIGDIREAHC | 35 |
| DQ002179 | B | CCR5 | CTRPNNNTRRSITIGPGRAFYATDIIGDIRQAQC | 34 |
| KC312595 | B | CCR5 | CVRPHNNTRKSIHIGPGRTFYATGEVIGDIRQAHC | 35 |
| AY173955 | B | CCR5 | CIRPNNNTRKSIHIGPGRAFYATGDIIGDIRQAHC | 35 |
| FJ653210 | B | CCR5 | CTRPNNNTRKGIHIGPGRAYYATEKITGDIRQAHC | 35 |
| AY043173 | C | CCR5 | CTRPNNNTRKSIRIGPGQTFYATGEIIGNIREAHC | 35 |
| KC312481 | B | CCR5 | CTRPNNNTRKGIHIGPGRTFYATGQIIGNIRQAHC | 35 |
| AF384329 | B | CCR5 | CTRPNNNTRKGIAIGPGRAVYATEKIVGDIRQAHC | 35 |
| HM215397 | B | CCR5 | CTRPNNNTRKSISLGPGRAWYATGQIIGNIRQAHC | 35 |
| FJ653122 | B | CCR5 | CTRFNNNTRKGIHIGPGRAFYATGDIIGNIRQASC | 35 |
| HQ678284 | B | CCR5 | CTRPNNNTRKSIHIAPGRAFYATGAIIGDIRKAYC | 35 |
| HQ708079 | C | CCR5 | CTRPNNNIRKGYTIGPGRAFFATGDIIGDIRQADC | 35 |
| AF384270 | B | CCR5 | CTRPGNNTRKSIHIAPGSAFYATGVIIGDIRKAHC | 35 |
| HQ644814 | B | CCR5 | CIRPNNNTRRSINIGPGRAIFRTGEVIGDIRQAHC | 35 |

| | | | | |
|---|---|---|---|---|
| AY124979 | B | CCR5 | CTRPSNNTRKSINIGAGRAIYAAGDIIGNTRQAYC | 35 |
| HQ644989 | B | CCR5 | CTRPNNNTRKSIPIGPGSAFYATGEIIGDIRQAHC | 35 |
| HQ708064 | C | CCR5 | CTRPHNNTRKSMRIGPGQTFYATEEVIGDIRQAYC | 35 |
| DQ235629 | C | CCR5 | CTRPSNNTRTSVRIGPGQTFYATGDIIGDIRQAHC | 35 |
| KC156284 | C | CCR5 | CTRPNNNTRQSMRIGPGQTFYATGAIIGNIRQAHC | 35 |
| AJ418536 | B | CCR5 | CTRPHNNTRKSIHIGPGRAFYATGQITGDIRQAYC | 35 |
| AF258962 | B | CCR5 | CTRPNNNTRKSIHLGPGRAFFTTGEIIGNIRQAYC | 35 |
| FJ376029 | C | CCR5 | TRGNNTRSIRIGPGQTFYATGDIIGDRQAHC | 31 |
| AF543920 | C | CCR5 | CTRPNNNTRKSVRIGPGQTFYATGAIIGDIREAHC | 35 |
| DQ516181 | B | CCR5 | CTRPNNNTRKGIHIGPGRAFYTTGAIIGNIRHAHC | 35 |
| EU575508 | B | CCR5 | CTRPNNNTRKSIHIGPGQAFYTTGAIIGDIRQAYC | 35 |
| JF507977 | B | CCR5 | CTRPNNNTKRDIPIGPGRAFYATDIVGDIRQAHC | 34 |
| JF896845 | B | CCR5 | CTRPNNNTRKSIGIGPGRAFYATGDIIGDIRQAHC | 35 |
| JF896873 | B | CCR5 | CTRPNNNTRKSIPLGPGSAFYATETIIGDIRQAHC | 35 |
| AF153130 | C | CCR5 | CARPNNNTRKSIRIGPGQAFYATGDIIGDIRQAHC | 35 |
| HQ644974 | B | CCR5 | CTRPNNNTRKSITIGPGRAFYATGNIIGDIRQAHC | 35 |
| DQ516289 | B | CCR5 | CTRPNNNTRKGIHIGPGRAFYTTGGIIGDIRQAHC | 35 |
| FJ376002 | C | CCR5 | CTRPNNNTRRVRIGPGQAFYATGDIIGDIRQAHC | 34 |
| EU293449 | C | CCR5 | CSRIGNNTRTSIGIGPGQAFFATGDIIGNIRKAHC | 35 |
| EU272293 | B | CCR5 | GTRPSNNTRKSINLGPGRAWYTTGQIIGDIRQAHC | 35 |
| EU604600 | B | CCR5 | CTRPSNNTRKSINVGPGRAWYATGQIIGDIRQAYC | 35 |
| EU744133 | B | CCR5 | CTRPNNNTRKSIHIGPGRAFYATGEIVGNIRQAHC | 35 |
| EU744080 | B | CCR5 | CTRYNNNTRRSIHMGPGRTFYATGDIIGDIRQAQC | 35 |
| JF896848 | B | CCR5 | CTRPNNNTRKSIHIAPGRAFYATENIVGDIRKAYC | 35 |
| HM246236 | B | CCR5 | CTRPNNNTRKSIHIGPGRAFYTTGDIIGDIRKAHC | 35 |
| EU744074 | B | CCR5 | CTRYNNNTRRSIPLGPGRAFYATGDIIGDIRQAQC | 35 |
| KF716498 | B | CCR5 | CTRPNNNTRRSIHLGPGQTLYATGDIIGDIRQAHC | 35 |
| KC312497 | B | CCR5 | CTRPNNNTRKGIHIGPGRTFYATGQIIGDIRQAHC | 35 |
| DQ002313 | B | CCR5 | CTRPNNNTRKGINIGPGRAWYTTGDIIGDIRKAHC | 35 |
| AY010784 | B | CCR5 | CIRPNNNTRKSIHIGPGKAFYATGDIIGDIRQAYC | 35 |
| DQ061429 | B | CCR5 | CTRPSNNTGKSITIGPGRAFYTTGEIIGDIRQAHC | 35 |
| EU272312 | B | CCR5 | YTRPNNNTRKSIPLGPGRAWYTTGQIIGDIRQAHC | 35 |
| AY228556 | C | CCR5 | CTRPNNNTRKSIRIGPGQTFYATNIIGDIRQAYC | 34 |
| U08784 | B | CCR5 | CTRPNNNTRKSIPLGPGQAWYTTGQILGDIRQAHC | 35 |
| GQ372990 | B | CCR5 | CLRPNNNTRKGIHIGPGRAFYTTGEIIGDIRQAHC | 35 |
| KC473830 | B | CCR5 | CTRPNNNTRKSIHIAPGRAFYATGAITGDIRQAHC | 35 |
| DQ110000 | B | CCR5 | CTRPNNNTRRSIHIAPGRAFYATGKIIGDIRQAHC | 35 |
| HM246227 | B | CCR5 | CTRPNNNTRKVQVGPGKALYITGSIIGDIRQAHC | 34 |
| DQ061776 | B | CCR5 | CTRPNNNTRKGIHIGPGRAFYATGEVIGNIRQAHC | 35 |
| HQ708091 | C | CCR5 | CTRPNNNTRRSYTIGPGRVLYATGSIIGDIRQAYC | 35 |
| HQ377420 | B | CCR5 | CTRPNNNTRKSIHIGPGSAFYATGEIIGDMRQAHC | 35 |
| FJ376004 | C | CCR5 | CTRPNNTKSVRIPGQTFYATGDIIGDIRQAHC | 32 |
| AF082372 | B | CCR5 | CTGPNNNTRRSISIGPGGAFYTTGDIIGDIRQAYC | 35 |
| AF153142 | C | CCR5 | CTRPNNNTRKSIRIGPGQTFYATGVIIGDIRQAHC | 35 |
| DQ061447 | B | CCR5 | CTRPNNNTRKSIHIGPGRAFYTTGEIIGDIRKAYC | 35 |
| JN002038 | B | CCR5 | CTRPNNNTRKSIHMGPGKAFYATGDIIGDIRRAHC | 35 |
| AF384255 | B | CCR5 | CIRPSNNTRKSIHMGPGRAWYATGSITGDIRQAHC | 35 |
| EU744132 | B | CCR5 | CTRPTNNTRKSIHIGPGRAFYATGDIIGDIRQAHC | 35 |
| AY585271 | C | CCR5 | CTRPHNNTRQSVRIGPGQAFFATREIIGDIRQAHC | 35 |
| FJ977094 | C | CCR5 | CARPGNNTRKSVRIGPGQAFYATGDIIGDIRQAHC | 35 |
| AF541062 | B | CCR5 | CVRPNNNTRRSIHIGLGRRFYTTEIVGDIRKAYC | 34 |
| AF384324 | B | CCR5 | CTRPNNNTRKGIHIGPGGAIYATGAIIEDIRQAHC | 35 |
| EU786673 | C | CCR5 | CTRPGNNTRKSMRIGPGQTFFATGDIIGDIRQAHC | 35 |
| HM246209 | B | CCR5 | CTRPSNNTRKSIPIGPGRAFYATGDIIGNIRQAHC | 35 |
| AY010842 | B | CCR5 | CTRPNNNTRKGIHIGPGSAFYATGDIIGDIRKAHC | 35 |
| DQ002115 | B | CCR5 | CTRPNNNTRKSITIGPGRAFYATGDIIGNIRQAHC | 35 |
| AF153136 | C | CCR5 | CTRPNNNTRKSIRIGPGQAFYTTGEIIGDIKQAHC | 35 |
| HQ644914 | B | CCR5 | CIRPNNNTRKSIHIGPGRAFYTTEIIGDIRKAYC | 34 |
| EU743981 | B | CCR5 | CTRPNNNTRKSIHIGPGRAFYATGEIIGDIRQAYC | 35 |
| HQ708065 | C | CCR5 | CIRPYNNTRKSMRIGPGQTFYATEEVIGNIRQAYC | 35 |
| AY170662 | C | CCR5 | CARPNNNTRKSVRIGPGQTFYATGGIIGDIRQAHC | 35 |
| FJ376014 | C | CCR5 | CTRPGNNTRRSVRIGPGQVFYATGEIIGDIRQAHC | 35 |
| HM179942 | C | CCR5 | CTRPGNNTRKGMWIGPGQAFYATGDIIGDIRQAYC | 35 |
| HQ708071 | C | CCR5 | CTRPNNNTRTSVRIGPGQTFFATGDVTGDIRQAFC | 35 |
| HQ644809 | B | CCR5 | CIRPNNNTRRSINIGPGRAIFRTGEIIGDIRQAHC | 35 |
| DQ061751 | B | CCR5 | CTKPNNNTRRSISLGPGRAYYATGDIIGDIRQAHC | 35 |
| HQ678272 | B | CCR5 | CTRPNNNTRKSIPMGPGKAFFTTDIIGEIRRAYC | 34 |
| GU204942 | B | CCR5 | CIRPNNNTRKSIHIAPGRAFYATGDIIGDIRQAHC | 35 |
| KF770290 | C | CCR5 | CIRPNNNTRRSVRIGPGQTFYATGAIIGDIRKAYC | 35 |
| AF491741 | B | CCR5 | CSRPNNNTRKSIPMGPGKAFYATGDIIGDIRQAHC | 35 |

| | | | | |
|---|---|---|---|---|
| DQ002096 | B | CCR5 | CTRPSNNTRRGIHIGPGRAFYTTGEVIGNIRAANC | 35 |
| HM246204 | B | CCR5 | CRPNNNTRKSIPMGPGQALYATGEIIGDIRQAHC | 34 |
| AY010860 | B | CCR5 | CTRPNNNTGTSIHMGPGRAVYATGDIIGNIRQAYC | 35 |
| AF391238 | C | CCR5 | CPRPNHNTRRSIRIGPGQAFYATGDIIGDIRQAHC | 35 |
| DQ002308 | B | CCR5 | CTRPNNNTRKSINIGPGRAWYTTGAIIGDIRKAHC | 35 |
| DQ002159 | B | CCR5 | CTRPNNNTRKSIHMGPGRAFYATGQIIGDIRQAHC | 35 |
| HQ644910 | B | CCR5 | CTRPNNNTRKSIHIGPGRAFYATGEIIGDIRRAYC | 35 |
| AF391237 | C | CCR5 | CTRPGNNTRQSIRIGPGQTFYATGDIIGDIRQAHC | 35 |
| EF657916 | B | CCR5 | CIRPNNNTMKSIHMGPGRAFYATGEIIGNIRQAHC | 35 |
| GU204920 | B | CCR5 | CARPNNNTRKSIHIAPGRAFHTTGSIIGDIRKAYC | 35 |
| KC156328 | C | CCR5 | CTRPNNNIRKSMRIGPGQTFYATGGIIGDIRQAHC | 35 |
| AF153183 | C | CCR5 | CTRPHNNTRKSVRIGPGQTFYATGDIIGDIRQAHC | 35 |
| FJ376012 | C | CCR5 | CIRPNNNTRKSMRIGPGQTFYATGDIIGDIRQAYC | 35 |
| KF770328 | C | CCR5 | CIRPNNNTRRSMRIGPGQAYFTTGEIIGNIRQAHC | 35 |
| KF384810 | B | CCR5 | CTRHNNNTRKSIHIGPGSAFYATGAIIGDIRQAHC | 35 |
| HM239534 | B | CCR5 | CTRLNNNTRKSIHIGPGQAFATGAIIGIRQAHC | 33 |
| EU272317 | B | CCR5 | CTRPNNNTRKSIPLKPGRAWYTTGQIIGDIRQAHC | 35 |
| AY736822 | C | CCR5 | CTRPGNNTRKSVRIGPGQAFYATNDIIGDIRQAYC | 35 |
| DQ235645 | C | CCR5 | CIRPQNNTVKSVRIGPGQTFYTTGQVVGDIRKAHC | 35 |
| JN687737 | C | CCR5 | CTRPNNNTRKSIRIGPGQAFYATGDIVGDIRQAHC | 35 |
| HM239592 | B | CCR5 | CTRPNNNTMKSITIGPGRAFYATGSIIGDIRQAHC | 35 |
| EU578309 | B | CCR5 | CTRPNNNTRKSIAIGPGRAFYTGEIIGDIRQAHC | 34 |
| EU576240 | B | CCR5 | CTRPNNNTRRSIPMGPGKVFYTADIIGDIRQAHC | 34 |
| AF067154 | C | CCR5 | CVRPNNNTRESIRIGPGQTFYATGEIIGDIRQAHC | 35 |
| FJ375969 | C | CCR5 | CTRPGTRKSIRIGPGQSFYATGIIGDIRQAHC | 32 |
| JF508129 | B | CCR5 | CTRPNNNTRKGIHIGPGRTFYATGEIVGDIRQAHC | 35 |
| JF507835 | B | CCR5 | CMRPGNNTRRSITIGPGRAFYAGEIIGDIRKAHC | 34 |
| EU578351 | B | CCR5 | CTRPNNNTRKSIHMGAGRAFYTGEIIGDIRQAHC | 34 |
| KC156335 | C | CCR5 | CNRPHNNTRKSMRIGPGQTFYATGDVTGDIRKAHC | 35 |
| EU744041 | B | CCR5 | CTRPSNNTRRGIHIGPGRAFYATGEIIGDIRQAHC | 35 |
| AF259023 | B | CCR5 | CTRPGNNTRKSIHIGPGRAFYTTGEIIGKIRQAHC | 35 |
| HM239635 | B | CCR5 | CTRPNNNTRKGIHLGPGKAFYATGDIIGNIRQAHC | 35 |
| AY887852 | C | CCR5 | CIRPGNNTRKSIRIGPGQTFYATGDIIGDIRQAHC | 35 |
| HM179915 | C | CCR5 | CTRPNNNTRKSIRIGPGQTFYATGDIIGDIRQAYR | 35 |
| JF896868 | B | CCR5 | CTRLNNNTRKSIFAPGRTFYAMGDIIGDIRKASC | 34 |
| HQ377470 | B | CCR5 | CTRPNNNTRKSIHMGPGGAVYATGAIIGNIRQAHC | 35 |
| AY887854 | C | CCR5 | CTRPGNNTRKSVRIGPGQTFFATGDIIGDIRQAHC | 35 |
| AY887872 | C | CCR5 | CTRPGNNTRRSVRIGPGQTFYATGDIIGDIRQAHC | 35 |
| JN687705 | C | CCR5 | CTRPNNNTRKSVRIGPGQTFYATGGIIGDIKQAHC | 35 |
| AY842816 | B | CCR5 | CTRPNNNTRKSIPMGPGRAMYATGDIIGDIRQAHC | 35 |
| EU576299 | B | CCR5 | CTRPNNNTRKSIHIGPGRTFYTTGDIIGDIRQAYC | 35 |
| AF254773 | C | CCR5 | CTRPNNNTRKSIRIGPGQTFYATNDIIGDIRQAYC | 35 |
| KC156240 | C | CCR5 | CIRPGNNTRRSVRIGPGQTFYATGDITGDIRKAHC | 35 |
| AF022266 | B | CCR5 | CTRPNNNTRKSISIGPGRAFYTTGEIIGNIRQAHC | 35 |
| FJ998107 | B | CCR5 | CTRPNNNTRRGIRIGPGRAFYATAIIGDIRQAHC | 34 |
| DQ061518 | B | CCR5 | CTRPNNNTRKSIHITPGRTFYATGEIIGDIRQAHC | 35 |
| EU577329 | B | CCR5 | CSRPNNNTRRSVHIGPGRAWYTGEIIGDIRQAHC | 34 |
| DQ061749 | B | CCR5 | CTRPNNNTRRSISLGPGRAYYATGDIIGDIRRAHC | 35 |
| AY842789 | B | CCR5 | CTRPSNNTRKSIHMGPGRVLYTTGGIIGDIRQAHC | 35 |
| DQ061531 | B | CCR5 | CTRPNNNTRRSIYIAPGSAFYATGEIIGDIRQAHC | 35 |
| AJ418521 | B | CCR5 | CTRPSNNTRKSIPIGPGRAFYTTGEIIGDIRQAHC | 35 |
| DQ002152 | B | CCR5 | CTRPNNNTRRNIHIGPGKAIYTTGEIIGDIRQAHC | 35 |
| DQ516124 | B | CCR5 | CTRPNNNTIKSIHIGPGRAFYTTGQIIGDIRQAHC | 35 |
| JF507907 | B | CCR5 | CTRPSNNTRQSVHIGPGRALYTTTKIIGDIRKAYC | 35 |
| AF022274 | B | CCR5 | CTRPNNNTRKSISIGPGRAFYATGEIIGDIRQAYC | 35 |
| KF770330 | C | CCR5 | CIRPNNNTRRSMRIGPGQALFTTGEIIGNIRQAHC | 35 |
| JX973084 | C | CCR5 | CIRPGNNMRKSVRIGPGQAFYATGDIIGDIRQAHC | 35 |
| HM239590 | B | CCR5 | CTRPNNNTRKSIPIGPGKAFYTTGDIIGDIRQAHC | 35 |
| EU744070 | B | CCR5 | CIRPNNNTRKSIHLGPGRAFYATGDIIGDIRQAQC | 35 |
| AY887853 | C | CCR5 | CIRPGNNTRKSVRIGPGQTFFATGDIIGDIRKAHC | 35 |
| AF199041 | B | CCR5 | CTRPNNNTRKSINMGQGRAWYTTGQIIGDIRQAHC | 35 |
| HQ644865 | B | CCR5 | CTRPGNNTRKSIPIGPGRAFYATGDIIGDIRQAHC | 35 |
| JF507846 | B | CCR5 | CVRPGNNTRRSITIGPGRAFYAGEIIGDIRKAHC | 34 |
| JF507941 | B | CCR5 | CTRPNNNTRKDIHIGPGRAFYATDIVGDIRQAHC | 34 |
| DQ110003 | B | CCR5 | CIRPNNNTRSIMGPGAFYATGDIIGDIRQAHC | 32 |
| JF507788 | B | CCR5 | CERPNNNTRKGIHIGPGRTFFATGEIIGDIRRAYC | 35 |
| EU744056 | B | CCR5 | CTRPNNNTRKSIHVGPGKTLYATGDIIGNIRQAHC | 35 |
| AF541018 | B | CCR5 | CIRPNNNTRKSIHIGPGRAFYTTGEIIGDIRQAHC | 35 |
| FJ653282 | C | CCR5 | CTRPNNNTRKSIRIGPGQTFFATEIVGNIREAHC | 34 |
| HM179735 | C | CCR5 | CTRPSNNTRKSMRIGPGQAFYATGEIIGNIREAHC | 35 |

| | | | | |
|---|---|---|---|---|
| KC473846 | B | CCR5 | CTRPSNNTRRGIHIGPGQAFYTTGQIIGDIRQAYC | 35 |
| AF153149 | C | CCR5 | CTRPINNTRWSVRIGPGQTFYATGDIIGDIRQAHC | 35 |
| AF153174 | C | CCR5 | CTRPNNNTRKSVRIGPGQAFYATDGIIGDIRQAHC | 35 |
| HQ708077 | C | CCR5 | CTRPNNNTRTSMRIGPGQTFYATGKVTGDIRQAYC | 35 |
| JF507950 | B | CCR5 | CTRPNNNTAKGIHIGPGRAMYATERIVGDIRRAHC | 35 |
| HM239570 | B | CCR5 | CERPNNNTRKSIHIAPGRTFYATGEIIGNIRQAHC | 35 |
| AB553913 | B | CCR5 | CTRPNNNTRKGIHMGPGRAIYTTDIIGDIRQAHC | 34 |
| AY887863 | C | CCR5 | CIRPGNNTRKSIRIGPGQAFYATGSIIGDIRQAHC | 35 |
| EU744021 | B | CCR5 | CTRPNNNTRKGIHIGPGRAFYTTGEIVGDIRQAHC | 35 |
| HM239617 | B | CCR5 | CTRPNNNTRRSVHIAPGAAFYATGAIIGQIRQAHC | 35 |
| AF541063 | B | CCR5 | CIRPNNNTRRGIHIGLGRRFYTTEIVGDIRKAYC | 34 |
| EU575330 | B | CCR5 | CIRPNNNTRKRITMGPGKVYYTTGQIIGDIRQAHC | 35 |
| AF391240 | C | CCR5 | CTRPNNNTRKSIRIGPGQTFFATNDIIGDIRQAYC | 35 |
| FJ798403 | B | CCR5 | CTRPNNNTRKGISIGPGRAFYATGDIIGDIRKAHC | 35 |
| FJ977091 | C | CCR5 | CARPGNNTKKSVRIGPGQTFYATGDIIGDIRQAHC | 35 |
| AF541031 | B | CCR5 | CTRPSNNTRKSIPIGPGRAFYATGEITGDIRQAHC | 35 |
| JN687717 | C | CCR5 | CTRPNNNTRKSIRIGPGQSFYATGDIIGDIRQAHC | 35 |
| DQ061843 | B | CCR5 | CTRPNNNTRKGIHVGPGAVFYATGQIIGDIRQAYC | 35 |
| HQ644913 | B | CCR5 | CTRPNNNTRKSIPIGPGRAFYTTEIIGNIRQAHC | 34 |
| AY253322 | C | CCR5 | CTRPGNNTRKSIRIGPGQAFFATGEVIGDIRQAHC | 35 |
| AY010803 | B | CCR5 | YKTQQLYKKSIHMGPGRAFYTTGDIIGDIRQAHC | 34 |
| EU272301 | B | CCR5 | CIRPNNNTRKSIHLGPGRAWYATGEIIGNIRQAHC | 35 |
| HQ644980 | B | CCR5 | CTRPNNNTRRSIHIGPGRAFYATGEIIGDIRQAHC | 35 |
| DQ061420 | B | CCR5 | CTRPNNNTRKSITIGPGRAFYTTGEIIGNIRQAHC | 35 |
| AF082360 | B | CCR5 | CTRPNNNTRKSIHIGPGRAFYTTGDIVGDIRQAHC | 35 |
| FJ653231 | B | CCR5 | CTRPSNNTSGSIHIGPGRAFDATKTITGDIRQVHC | 35 |
| DQ002279 | B | CCR5 | CTRPNNDTKRSVNIVPGRAIYTTDIIGDIRQAHC | 34 |
| AF021499 | B | CCR5 | CTRPNNNTRKSIHIGPGGAFYTTGQIIGDIRQAYC | 35 |
| HM239552 | B | CCR5 | CTRPNNNTRKSIIGPGRAFYATDIIGDIRQAHC | 33 |
| HQ708031 | C | CCR5 | CTRPNNNTRKSTRIGPGQSFYATGDIIGNIRQAHC | 35 |
| DQ061844 | B | CCR5 | CTRPNNNTRKGIHMGPGAVFYATGQIIGDIRQAYC | 35 |
| DQ002237 | B | CCR5 | CTRPSNNTRKSIHIAPGRAFYATGEIIGDIRQAHC | 35 |
| AF541060 | B | CCR5 | RVRPNNNTRRSIHIGLGRRFYTTEIVGDIRKAYC | 34 |
| HQ644789 | B | CCR5 | CTRPNNNTRRGIHIGPGRAFYTTGEIVGDIRQAHC | 35 |
| AY124977 | B | CCR5 | CTRLNNNTRKSINIGPGRAFYATGDIIGDIRQAHC | 35 |
| GU204943 | B | CCR5 | CIRPNNNTRKSIHIPGRAFATGDIIGDIRQAHC | 33 |
| AY835447 | B | CCR5 | CTRHNNNTRKSINIGPGRAFYATGKIIGDIRQAHC | 35 |
| KF770377 | C | CCR5 | CIRPGNNTSRSIRIGPGQTFYATGRIVGDIRLAHC | 35 |
| HQ644985 | B | CCR5 | CTRPNNNTRKSIRIGPGSAFYATGEIIGDIRQAHC | 35 |
| EF580005 | B | CCR5 | CERPGNNTSKGIHIGPGRAFYATEDIIGDIRKAHC | 35 |
| DQ645382 | B | CCR5 | CTRPNNNTRKSIHLGPGKTLYATDIIGNIRQAHC | 34 |
| GU362881 | B | CCR5 | CTRPNNNTRKGIHIGPGRTLYATGEIIGDIRKAHC | 35 |
| HQ644838 | B | CCR5 | CTRPNNNTRKSITIGPGRAFYTTGDIIGDIRQAYC | 35 |
| AF254782 | C | CCR5 | CTRPNNNTRRSMRIGPGQTFYATGEIIGDIRQAYC | 35 |
| HQ644917 | B | CCR5 | CIRPNNNTRKSIHIGPGRALYATEIIGDIRKAYC | 34 |
| JF896834 | B | CCR5 | CTRLNNNTRKGIHIGPGRAFYATTDIIGDVRQAHC | 35 |
| AY010861 | B | CCR5 | CTRPNNNTRTSIHMGPGRALYATGDIIGNIRQAYC | 35 |
| JF507905 | B | CCR5 | CTRPSNNTRQSVHIGPGRALYTTNIIGDIRKAYC | 34 |
| JF896832 | B | CCR5 | CTRPNNNTRKSIPIGPGRAFYATGHIIGNIRQAHC | 35 |
| AF153184 | C | CCR5 | CTRPNNNTRKSIRIGPGQAFYATNAIIGNIRQAHC | 35 |
| EU576418 | B | CCR5 | CTRPNNNTRKSIPMGPGKAFYARGDITGDIRKAYC | 35 |
| AY010785 | B | CCR5 | CIRPNNNTRKSIHMGPGKAFYATGDMIGDIRQAYC | 35 |
| JF507966 | B | CCR5 | CTRPNNNTRKDIHIGPGRAMYATGEIVGDIRRAHC | 35 |
| FJ375986 | C | CCR5 | CTRPNNNTRTSMRIGGQTFYATGDIIGDIRQAHC | 34 |
| AF384265 | B | CCR5 | CTRPNNNTRKSINIGPGRAFFATGDIIGDIRQAHC | 35 |
| JF507870 | B | CCR5 | CTRPGNNTRRSINIGPGRAFYTGEIIGNIRQAHC | 34 |
| DQ177201 | B | CCR5 | CTRPNNNTRKSIPMGPGKAFYATGEIIGDIRQAHC | 35 |
| DQ002272 | B | CCR5 | CTRPNINTKRSVNIVPGRAIYTTDIIGDIRQAHC | 34 |
| EU743975 | B | CCR5 | CTRPNNNTRKGIHIGPGKAFYATGEIIGDIRQAHC | 35 |
| DQ061803 | B | CCR5 | CIRPNNNTRKGIHIGPGRAFYATGATIGNIRQARC | 35 |
| EU744174 | B | CCR5 | CTRPNNNTRRSIHMGPGGALYTTGAIIGNIRQAHC | 35 |
| DQ869021 | B | CCR5 | CTRPGNNTRRSIPMGPGRAWYAIGEITGNIRKAHC | 35 |
| HM368242 | B | CCR5 | CTRPNNNTRRGIHIGPGGAFYSTGDIIGNIRQAYC | 35 |
| KF770393 | C | CCR5 | CTRPSNNTRRSVRIGPGQAFYTTGEIIGDIRQAHC | 35 |
| GU362883 | B | CCR5 | CTRPNNNTRRSIHIAPGRAFYATGEIIGDIRQAHC | 35 |
| HQ644799 | B | CCR5 | CTRPNNNTRKSINIGPGRAIYTTGEIIGDIRQAHC | 35 |
| JF507918 | B | CCR5 | CIRPNNNTRKDIHIGPGRAFYATGIVGDIRQAHC | 34 |
| FJ376022 | C | CCR5 | CTRPGNNTKSFRIGPGQTFFATGEIIGNIRQAHC | 34 |
| AY713408 | B | CCR5 | CTRPNNNTRKSIQLGPGRAWYTTGQIIGDIRQAHC | 35 |
| JX508874 | C | CCR5 | CTRPNNNTRRSIRIGPGQTFYATGEIIGNIREAYC | 35 |

| | | | | |
|---|---|---|---|---|
| KC312596 | B | CCR5 | CVRPNNNTRKSIHLGPGSAFYATGEIIGNIRQAHC | 35 |
| EU744071 | B | CCR5 | CTRPHNNTRKSIHLGPGRAFYATGDVIGNIRQAHC | 35 |
| DQ002168 | B | CCR5 | CTRPNNNTRKSIHMGPGRALYATGAIIGDIRQAHC | 35 |
| EU576838 | B | CCR5 | CTRPNNNTRRDITIGPGRVFYTGEIIGDIRQAHC | 34 |
| AF384334 | B | CCR5 | CTRPNNNTRKSIPLGAGRAWYATGDIIGDIRQAHC | 35 |
| GU945319 | C | CCR5 | CTRPNNNTRQSIRIGPGQVFYATGDIIGDIRQAYC | 35 |
| DQ177199 | B | CCR5 | CIRPNNNTRRSIPIGPGRAFYATGGIIGDIRKAYC | 35 |
| EU744138 | B | CCR5 | CTRPTNNTRKSIHIGPGRAFYATGDIVGNIRQAHC | 35 |
| KC312394 | B | CCR5 | CTRPNNNTRKSIHIGPGSAFYTTGAIIGDIRQAHC | 35 |
| HQ708029 | C | CCR5 | CTRPNNNTRKSTRIGPGQSFYATGDIIGEIRQAHC | 35 |
| HM239559 | B | CCR5 | CTRPSNNTRKGIHMGPGKAFYATGEIRGDIRQAHC | 35 |
| AF153153 | C | CCR5 | CPRPINNTRWSVRIGPGQTFYATGDIIGDIRQAHC | 35 |
| HQ708030 | C | CCR5 | CTRPNNNTRKSTRIGPGQSFYATGEIIGNIRQAHC | 35 |
| JF507730 | B | CCR5 | CIRPNNNTRKSIHIGPGRTFFATGEIIGDIRRAHC | 35 |
| AF355728 | B | CCR5 | CTRPSNNTRKSIPLGPGRAFYATGDIIGNIRKAHC | 35 |
| EF688453 | B | CCR5 | CTRPNNNTRKGIHIGPGRAWYTTGEIIGDIRQAHC | 35 |
| HQ645012 | B | CCR5 | CTRPNNNTRKSINLGPGRALYTTGNIIGNIRQAHC | 35 |
| AF259012 | B | CCR5 | CTRPSNNTRKSIHIGPGRAFYTTGEVTGDIRQAHC | 35 |
| AJ418517 | B | CCR5 | CTRPNNNTRKSIHIGPGRAFYTTGDIIGNIRQAHC | 35 |
| HM179948 | C | CCR5 | CTRPGNNTRESMWIGPGQAFYATGDIIGDIRQAYC | 35 |
| EU744017 | B | CCR5 | CTRPNNNTRKSIHMGPGRAFYTTGEIIGDIRQAHC | 35 |
| HM239562 | B | CCR5 | CIRPNNNTRKGIHIGPGKAFYTTGSIIGDIRQAHC | 35 |
| HM239556 | B | CCR5 | CTRPSNNTRKGIHMAVGRAFYATGQIIGDIRQAYC | 35 |
| JN687824 | B | CCR5 | CTRPNNNTRKSINMEPGRAWYATGEIIGNIRQAHC | 35 |
| DQ235625 | C | CCR5 | CTRPNNNTRRSMRIGPGQAFYATGDIIGDIRQAHC | 35 |
| AY887881 | C | CCR5 | CVRPNNNTRKSIRIGPGQTFYATGNIIRDIRQAHC | 35 |
| EU272251 | B | CCR5 | CIRPNNNTRKSIPLRPGKAWYTTGQIIGDIRQAHC | 35 |
| FJ375988 | C | CCR5 | CTRPNNNTRQSIRIGPGQAFYATQGIIGDIRKAYC | 35 |
| KC473842 | B | CCR5 | CTRPGNNTRKSISLGPGRVFYATGDIIGDIRQAYC | 35 |
| AX455929 | C | CCR5 | CTRPGNNTRKSVRIGPGQAFYATGDIIGDIRQAHC | 35 |
| KF770319 | C | CCR5 | CTRPSNNTRKSIRIGPGQTFFATDDIIGDIRKAYC | 35 |
| AF541013 | B | CCR5 | CTRPNNNTRKSIPIGPGRALYATGDIIGDIRKAHC | 35 |
| EU744137 | B | CCR5 | CTRPTNNTRKSIHLGPGRAFYATGDIVGNIRQAHC | 35 |
| DQ061810 | B | CCR5 | CIRPNNNTRKGIHIGPGRAFYATGAIIGNIRQAHC | 35 |
| HQ377468 | B | CCR5 | CTRPNNNTRKGIHIGPGGAVYATGAIIGNRRQAHC | 35 |
| AF384249 | B | CCR5 | CTRPNNNTRKSIHMGPGKAFYTTGNIIGDIRKAHC | 35 |
| EF657926 | B | CCR5 | CIRPNNNTRKSIHMGPGRAFYTTGEIIGNIRQAHC | 35 |
| HM215428 | B | CCR5 | CTRPSNNTRKSIHLGQGRAWYTTGKIIGDIRQAHC | 35 |
| AF310131 | B | CCR5 | CTRPNNYTRKHIHLGARKAFYTGEIIGDIRQAHC | 34 |
| EF688447 | B | CCR5 | CTRPNNNTRKSIPIGPGRAFFTTGDIIGDIRQAHC | 35 |
| HQ644820 | B | CCR5 | CKRPNNNTRRSIHIGPGRAFFATGDIIGDIRQAHC | 35 |
| GU455519 | B | CCR5 | CTRPNNNTKKSISIGPGRAFYTTGQIIGDIRQAHC | 35 |
| EU576991 | B | CCR5 | CTRPNNNTRKSIPIGPGRAFYATGEIIGNIRQAHC | 35 |
| EU578539 | B | CCR5 | CTRPNNNTRRSIHLGPGAAFITGEIIGDIRQAYC | 34 |
| AY510063 | C | CCR5 | CTRPSNNTRKSVRIGPGQTFYATGIIIGDIRQARC | 35 |
| HM239550 | B | CCR5 | CTRSNNTRKSIHIGPGRAFYATGEIIGNIRQAHC | 34 |
| KC473828 | B | CCR5 | CTRPNNNTRKSIHIAPGGAFFATGDIIGDIRKAHC | 35 |
| DQ869020 | B | CCR5 | CTRPSNNTRKSITIGPGRAFYATGDIIGDIRQAHC | 35 |
| AF541065 | B | CCR5 | CIRPNNNTKRSIHIGLGRRFYTTEIVGDIRKAYC | 34 |
| AF384318 | B | CCR5 | CTRPNNNTRKGIHIGPGGAFYATGAITGDIRQAHC | 35 |
| DQ002255 | B | CCR5 | CTRPNNNTRKSIPIGPGRAFYTTGEIIGNIRQAHC | 35 |
| JN687762 | B | CCR5 | CTRPNNNTRKSIHMGPGRAFYTTGDIIGDIRQANC | 35 |
| EF657930 | B | CCR5 | CIRPNNNTRKSIPMGPGRAFYATGGIIGDIRQAHC | 35 |
| AY173952 | B | CCR5 | CTRPSNNTRKSIHIGPGRAFYTTGNIIGDIRQAHC | 35 |
| EU576936 | B | CCR5 | CTRPGNNTRKSIPIGPGRAFYTTGDIIGDIRKAHC | 35 |
| HQ377459 | B | CCR5 | CTRPNNNTRKGIHIGPGGAVYATGAIIGNIRQAHC | 35 |
| AF384278 | B | CCR5 | CIRPNNNTRTSMPMGPGRAFYAMGDIIGDIRQAHC | 35 |
| DQ061506 | B | CCR5 | CTRPNNNTRKSITIGPGRAFYTTGEIIGEIRQAHC | 35 |
| AJ418530 | B | CCR5 | CTRPSNNTRKSIPIGPGRAFYTTGEIIGDIRKAHC | 35 |
| U16217 | B | CCR5 | CTRPNNNTRKGIHMGCGRTFYATGEIIGDIRQAHC | 35 |
| FJ375977 | C | CCR5 | CTRPNNNTRKSVRIGPGQTFYATGDIIGREAHC | 33 |
| EF657911 | B | CCR5 | CIRPNNNTKKSIPMGPGRAFYATGDIIGNIRQAHC | 35 |
| DQ061807 | B | CCR5 | CIRPNNNTRKGIHIGLGRAFYATGEIIGNIRQAHC | 35 |
| HM246202 | B | CCR5 | CRPNNNTRKGIHMGFGKTLYATGEIVGNIRQAHC | 34 |
| HQ678289 | B | CCR5 | CTRPNNNTRKSIPMGPGRAFYTTGQIIGDIRKAYC | 35 |
| AF180902 | B | CCR5 | CTRPNNNTRRSIHIGPGRALYTTGQIIGDIRQAYC | 35 |
| FJ376016 | C | CCR5 | CTRPSNNTRTSIRIGPGQAFYATGAITGDIRQAHC | 35 |
| JN687726 | C | CCR5 | CTRPGNNTRKSTRIGPGQAFFTAEIIGDIRQAHC | 34 |
| DQ904348 | C | CCR5 | CIRPNNNTRKSIRIGPGQTFYATNAIIGDIRQAHC | 35 |
| FJ653132 | B | CCR5 | CTRFNNNTRKSVHIGPGRAFYATGDIIGNIRQASC | 35 |

| | | | | |
|---|---|---|---|---|
| JF508037 | B | CCR5 | CARPNNNTRKGIHIGPGRAFYTTGEIIGDIRQAHC | 35 |
| KC312499 | B | CCR5 | CTRPNNNTRKGIHIGPGKTFYATGQIIGDIRQAHC | 35 |
| AY835449 | B | CCR5 | CTRPNNNTRKSIHIAPGRAFYATGEIIGDIRKAYC | 35 |
| AY887892 | C | CCR5 | CIRPGNNTRKSVRIGPGQAFYATGEIIGDIRQAHC | 35 |
| EU578432 | B | CCR5 | CTRPNNNTRKSIHLGPGSVFYTGEIIGDIRQAHC | 34 |
| DQ002100 | B | CCR5 | CTRPNNNTRRGIHIGPGRAFYTTGEVIGNIRAANC | 35 |
| HQ644837 | B | CCR5 | CTRPNNNTRRSITIGPGRAFYTTGDIIGDIRQAYC | 35 |
| HQ644998 | B | CCR5 | CIRPNNNTRKSIPIGPGRAFYATGDIIGDIRQAHC | 35 |
| EU744136 | B | CCR5 | CTRPNNNTRKGIHIGPGRAFYATGDIIGNIRQAHC | 35 |
| HM239585 | B | CCR5 | CTRPNNNTRTSIHMGPGKAFYTTGDIIGDIRKAYC | 35 |
| AF384283 | B | CCR5 | CTRPNNNTRKSIHIGPGRAFYTTGQIVGDVRKAHC | 35 |
| FJ375998 | C | CCR5 | CRPNNTRKMRIGPGQTYATGDIIGIRAHC | 29 |
| HQ678251 | B | CCR5 | CTRPNNNTRKRINLGPGKTFYATGEIIGDIRQAHC | 35 |
| AY835448 | B | CCR5 | CTRPNNNTSKSIHMGPGGAFFATGRIIGDIRKAYC | 35 |
| L48068 | C | CCR5 | CIRPGNNTRKSVRIGPGQTFFATGEIIGKIREAHC | 35 |
| AF082361 | B | CCR5 | CIRPNNNTRKSIHIGPGRAFYAAGDIIGDIRQAHC | 35 |
| FJ376031 | C | CCR5 | CTRGNNTRKSIRIGPGQTLYATGEIIGDIRAHC | 33 |
| HQ708081 | C | CCR5 | CTRPNNNTRRSYTIGPGRAFYATGAIIGDIRQAYC | 35 |
| AY887859 | C | CCR5 | CTRPGNNTRQSIRIGPGQTFYATGEVIGDIRQAHC | 35 |
| AF194975 | B | CCR5 | CTRPNNNTRRSINIGPGRAFYATGDIIGDIRQAHC | 35 |
| HM239612 | B | CCR5 | CTRPNNNTRKSIPIGPGRAYVATGDIIGDIRKAHC | 35 |
| AF541075 | B | CCR5 | CVRPSNNTRRGIHIGLGRRFYTTEIVGDIRKAYC | 34 |
| JF896854 | B | CCR5 | CTRPNNNTRKSIHMGPGAFYATGDIIGDIRQAHC | 34 |
| FJ376030 | C | CCR5 | CTRPYNNTRRSVRIGPGQAFYTTGEVIGDIRQAYC | 35 |
| AY123278 | C | CCR5 | CVRPHNNTRKSIRIGPGQAFYATESIIGDIRQAHC | 35 |
| KC473832 | B | CCR5 | CTRPGNNTRKSISIGPGRAFYATGDIIGDIRQAHC | 35 |
| HM239545 | B | CCR5 | CTRPSNNTRGIHMAVGRAFYATGQIIGNIRQAHC | 34 |
| AY426110 | B | CCR5 | CTRPNNNTRKSIHIGPGRALYTTGEIIGDIRQAHC | 35 |
| EU744139 | B | CCR5 | CTRPTNNTRKSIHIGPGRAFYATGDIIGNIRQAHC | 35 |
| DQ061740 | B | CCR5 | CTRPNNNTRRSISLGPGRAYYATGDIIGDIRQARC | 35 |
| JF507922 | B | CCR5 | CTRPNNNTRKSISIGPGRAFYATGGIIGDIRQAHC | 35 |
| AF541007 | B | CCR5 | CTRPSNNTRKSIPMGPGRAFYTTGEIIGNIRQAHC | 35 |
| JF896839 | B | CCR5 | CTRLSNNTRKGIHIGPGRAFYTTGEIIGDIRQAHC | 35 |
| EU578625 | B | CCR5 | CTRPNNNTRKSVHIGPGKVFYTGEIIGDIRQAHC | 34 |
| U79720 | B | CCR5 | CTRPNNNTRKSVSLGPGSAWYATGDIIGDIRQAHC | 35 |
| JF507748 | B | CCR5 | CTRPNNNTRKSIHIGPGRTFFATGEIIGDIRRAYC | 35 |
| HM239599 | B | CCR5 | CTRPSNNTRKGIHAWGRAFYATGQIIGDIRQAHC | 34 |
| AF153187 | C | CCR5 | CTRPGNNTRKSIRIGPGQTFYATNDIIGDIRSAHC | 35 |
| U04909 | B | CCR5 | CTRPNNNTRKSIPMGPGKAMYATGEIIGDIRKAYC | 35 |
| AJ418485 | B | CCR5 | CTRPNNNTRKGIHIGPGKALYATGDIIGKIRQAHC | 35 |
| HQ644856 | B | CCR5 | CTRPGNNTRKSIHIGPGRAFYTTGDIIGDIRQAHC | 35 |
| EF579975 | B | CCR5 | CGRPGNNTSKGIHMGPGRAFYATENIIGDIRKAHC | 35 |
| U08700 | B | CCR5 | CTRPNNNTRKSIHMGWGRAFFSTGELIGNIRQAHC | 35 |
| AY170661 | C | CCR5 | CTRPSNNTRKSIRIGPGQAFYATNEIIGDIRQAHC | 35 |
| HM239520 | B | CCR5 | CSRPNNNTRKSIPIGPGRAFYATGDIIGDIRQAHC | 35 |
| DQ002173 | B | CCR5 | CTRPNNNTRRSITIGPGRAFYGTDITGDIRQAHC | 34 |
| FJ376020 | C | CCR5 | CTRNNNTRKSIRIGPGQTFYATGDIIGNIRQAHC | 34 |
| EF688445 | B | CCR5 | CTRPNNNTSKGIHMGPGKAFYATGKIIGDIRQAHC | 35 |
| EU272322 | B | CCR5 | CTRPNNNTRKSIPLGPGRAWYTTGQIIGDIRQAHC | 35 |
| AF384253 | B | CCR5 | CTRPNNNTRKSIHMGPGRAFYTTGEIIENIRQAHC | 35 |
| KC473829 | B | CCR5 | CIRPNNNTRKGIHIGPGRAFYATGDIIGNIRKAHC | 35 |
| HM239630 | B | CCR5 | CTRPNNNTRKSISIGPGRAFYATGEVIGIRQAC | 33 |
| HQ708074 | C | CCR5 | CTRPNNNTRTSMRIGPGQTFYATGEVTGDIRQAYC | 35 |
| DQ235616 | C | CCR5 | CTRPNNNTRQSIRIGPGQTFFATKGIIGDIRQAYC | 35 |
| EF688458 | B | CCR5 | CTRPNNNTRKGIPIGPGKAFYATGEIIGDIRQAHC | 35 |
| FJ375975 | C | CCR5 | CTRPNNNTRKSRIGPGQAFYATDIIGDIREAHC | 33 |
| KF770359 | C | CCR5 | CTRPNNNTRESIRIGPGQTYYAMGDIIGDIRQAHC | 35 |
| HQ678262 | B | CCR5 | CIRPNNNTRKSIPMGPGKAFFTTNIIGDIRQAHC | 34 |
| DQ382364 | C | CCR5 | CARPGNNTRKSVRIGPGQTFFATGDIIGDIRKAHC | 35 |
| KC312582 | B | CCR5 | CVRPNNNTRKSIRIGPGSAFYAAGEIIGNIRQAHC | 35 |
| U79721 | B | CCR5 | CTRPNNNTRRSINIGPGRALYATGEIIGDIRQAHC | 35 |
| AY010802 | B | CCR5 | CTRPNNNTRTSIPMGPGRALYATGDIIGNIRQAYC | 35 |
| HM239553 | B | CCR5 | CIRPNNNTRKGIHMGFGKTLYATGEIVGNIRQAHC | 35 |
| AY835436 | B | CCR5 | CTRPNNNTRKSINIGPGRAFVATGAIIGDIRQAHC | 35 |
| EF643659 | B | CCR5 | CTRPNNNTRKSIHIGPGRAFYATGAIIGNIRQAHC | 35 |
| FJ653221 | B | CCR5 | CTRPSNNTSTSIHIGPGRAFDATKTIIGDIRQAHC | 35 |
| HM368227 | B | CCR5 | CTRPNNNTRKSIHIAPGRTFYTTGEIIGDIRQAHC | 35 |
| DQ061809 | B | CCR5 | CTRPSNNTRKGIHIGPGRAFYATGEITGDIRQAHC | 35 |
| HQ644800 | B | CCR5 | CTRPNNNTRKSINIGPGRAFFATGEIIGDIRQAHC | 35 |
| HM239582 | B | CCR5 | CTRPNNNTRKSIPLGPGRAFYTTGDIIGDIRKAYC | 35 |

| | | | | |
|---|---|---|---|---|
| AF259013 | B | CCR5 | CTRPSNNTRKSINIGPGRAWYTTGQITGDIRQAHC | 35 |
| HQ644919 | B | CCR5 | CIRPNNNTRKSIHIGPGRALYTTEIIGDIRKAYC | 34 |
| AY887862 | C | CCR5 | CIRPNNNTRQSIRIGPGQAFFATGDIIGDIRQAYC | 35 |
| HM179830 | C | CCR5 | CTRPGNNTRQSMRIGPGQTFYATGDIIGDIRQTHC | 35 |
| KC596067 | B | CCR5 | CTRPGNNTRRSISIGPGRAFYTMGQIIGNIREAHC | 35 |
| HQ644961 | B | CCR5 | CTRPNNNTRKSITIGPGRAFYATGEIIGDIRKAHC | 35 |
| KF770399 | C | CCR5 | CTRPGNNTRKSMRIGPGQTFYATGDIVGDIRKAHC | 35 |
| EF657929 | B | CCR5 | CMRPNNNTRKSIHMGPGRAFYTTGEIIGNIRQAHC | 35 |
| JN687741 | B | CCR5 | CTRPSNNIRKSIHMGWGRAFYATGEITGDIRQAHC | 35 |
| AF153167 | C | CCR5 | CVRPNNNTRRSVRIGPGQTFYATGDIIGDIRQAHC | 35 |
| EF600088 | B | CCR5 | CIRPNNNTRQGMHIGPGKALYTTNIIGNIRQAHC | 34 |
| HQ645004 | B | CCR5 | CTRPNNNTRRSITIGPGRAFYATGDIIGDIRRAQC | 35 |
| EF175209 | B | CCR5 | CIRPNNNTRKSIPMGPGKAFYATGDIIGNIRLAYC | 35 |
| AY426116 | B | CCR5 | CTRPNNNTRKSIHIGPGRAIYTTGKIIGDIRQAHC | 35 |
| EU272323 | B | CCR5 | CTRPNNNTRKSIPLGPGRAWYTTGIIGDIRQAHC | 34 |
| HQ644863 | B | CCR5 | CTRPNNNTRKSIHMGPGRAFYATEDVIGDIRQAHC | 35 |
| HM215400 | B | CCR5 | CIRPNNNTRKSIPLGQGRAWYTTGQIIGDIRQAHC | 35 |
| EF600084 | B | CCR5 | CIRPNNNSRQGIHIGPGKALYTTKIIGNIRQAHC | 34 |
| EU576214 | B | CCR5 | CTRPSNNTRKGIHIGPGRAFYATDIIGEIRQAHC | 34 |
| KC473825 | B | CCR5 | CTRPGNNTRKSIHIGPGRAFYATGDIIGDIRKAHC | 35 |
| GU455525 | B | CCR5 | CIRPNNNTRKGIHIGPGRAFYTTGQIIGDIRQAHC | 35 |
| AY510064 | C | CCR5 | CTRPNNNTRKSARIGPGQTFYAMGDIIGDIRQAHC | 35 |
| HM246225 | B | CCR5 | CTRPGNNTRKGIHIGPGRAFYATGDIIGDIRQAHC | 35 |
| AF199032 | B | CCR5 | CTRPNNNTRKSIPIGPGRAFYTTGQIIGDIRQAHC | 35 |
| EF600093 | B | CCR5 | CTRPNNNTRKSINIGPGRALYTTGDIIGDIRQAHC | 35 |
| FJ670521 | C | CCR5 | CTRPGNNTRKSMRIGPGQTFFATGDIIGNIRQAHC | 35 |
| DQ061480 | B | CCR5 | CIRPNNNTRKSITIGPGRAFYTTGEIIGEIRQAHC | 35 |
| DQ516122 | B | CCR5 | CIRPNNNTIKSIHIGPGRAFYTTGQIIGDIRQAHC | 35 |
| EU575728 | B | CCR5 | CMRPNNNTRKGIHIGPGGAFYATGDIIGNIRQAHC | 35 |
| KC156340 | C | CCR5 | CTRPHNNTRKSMRIGPGQAFYATGDVTGDIRKAHC | 35 |
| AF541073 | B | CCR5 | CIRPNNNTRRGIHIGLGRRFYATEIVGDIRKAYC | 34 |
| AY842811 | B | CCR5 | CTRPNNNTRKSIHMGPGRAMYATGDIIGDIRQAHC | 35 |
| EF657903 | B | CCR5 | CIRPNNNTRKSIPMGPGRAFYATGEIIGNIRQAHC | 35 |
| AJ418498 | B | CCR5 | CTRPSNNTRKSINIGPGRAFYTTGEIIGDIRKAHC | 35 |
| DQ516259 | B | CCR5 | CTRPNNNTRKGIHMGPGRALYTTGAIIGDIRQAHC | 35 |
| JF507945 | B | CCR5 | CTRPNNNTRRNIHIGPGRAMYATGQIIGNIRQAHC | 35 |
| AY010801 | B | CCR5 | YKTQQQYKTSIPMGPGRAMYATGDIIGNIRQAYC | 34 |
| AY713412 | B | CCR5 | CTRPNNNTRKSIHMGPGRAFYTTGDIIGDIRKAHC | 35 |
| AF112548 | B | CCR5 | CTRPNNNTRKGIHIGPGRTFYTTGEIIGDIRQAHC | 35 |
| AF194976 | B | CCR5 | CTRPNNNTRRSISIGPGRAFYATGDIIGDIRQAHC | 35 |
| U08698 | B | CCR5 | YKTQQQYKKKYTYGMGESIYATGEIIGNIRQAHC | 34 |
| EU744083 | B | CCR5 | CTRHHNNTRRSIHLGPGRAFYATGDVIGDIRQAQC | 35 |
| EF600089 | B | CCR5 | CIRPNNNTRQGIHIGPGRALYTTNIIGNIRQAHC | 34 |
| AY010843 | B | CCR5 | CTRPNNNTRKGIHIGPGSAFYATGDIIGDIRQAHC | 35 |
| JQ779253 | C | CCR5 | CTRPGNNTRKSIRIGPGQVFFTATDIIGNIREAHC | 35 |
| AF543911 | C | CCR5 | CTRPNNNTRKSIRIGPGQTFYATDAIIGNIREAHC | 35 |
| HM239571 | B | CCR5 | CIRPNNNTRRSIQMGPGKTFFTTGDIIGDIRRAHC | 35 |
| AY835442 | B | CCR5 | CTRPNNNTRKSIHIGPGRAFYATGGVIGDIRQAHC | 35 |
| JF507986 | B | CCR5 | CTRPNNNTRKSINIGPGRAFYTTGAIIGDIRQAHC | 35 |
| HQ644900 | B | CCR5 | CTRPNNNTRKSIHIGPGRAFYTTGDITGDIRKAYC | 35 |
| HQ377464 | B | CCR5 | CTRPNNNTRKSISMGPGRAFFATGEIIGNIRQAHC | 35 |
| DQ869031 | B | CCR5 | CTRPNNNTRKGIHMGPGRTFYATGEIIGDIRQAHC | 35 |
| HM246208 | B | CCR5 | CTRPNNNTRRSISIGPGRAFFATDIIGDIRQAHC | 34 |
| KC156272 | C | CCR5 | CTRPNNNTRKSMRIGPGQTFYATGAIIGDIRQAHC | 35 |
| DQ110004 | B | CCR5 | CRPNNNTRKSIMGPGRAFYATGDIIGDIRQAHC | 33 |
| KF766538 | C | CCR5 | CIRPNNRKSIRIGPGQTFYATGEIIGDIRQAHC | 33 |
| AF254771 | C | CCR5 | CTRPGNNTRKSVRIGPGQAFFATNDIIGDIRQAHC | 35 |
| U08454 | C | CCR5 | CTRPNNNTRKSIRIGPGQTFYATNEIIGNIREAHC | 35 |
| AF541078 | B | CCR5 | CIRPNNNTKRSIHLGLGRRFYTTEIIGDIRKAYC | 34 |
| KC156223 | C | CCR5 | CIRPGNNTRRSMRIGPGQTFYATGDITGDIRKAHC | 35 |
| AY010883 | B | CCR5 | CTRPNNNTRTSIPMGPGRAMYATGDIIGNIRHAYC | 35 |
| GU204924 | B | CCR5 | CIRPNNNTRKSIHIGPGRAFYTTGEIIGDIRQAYC | 35 |
| AY713417 | C | CCR5 | CTRPNNNTRQSIRIGPGQTFYATGEIIGDIRQAHC | 35 |
| HQ678269 | B | CCR5 | CTRPSNNTRKGIHMGPGRAFYATTNIAGDIRKAHC | 35 |
| JF896840 | B | CCR5 | CIRPGNNTRKSIQMGFGRAIYTTGDIIGIRQAHC | 34 |
| EU744040 | B | CCR5 | CTRPSNNTRKGIHMGPGRAFYATGEIIGDIRQAHC | 35 |
| DQ002143 | B | CCR5 | CTRPNNNTRRGIHIGPGRAFYTTGEIIGNIRQAYC | 35 |
| AF153190 | C | CCR5 | CTRPNNNTRKSIRIGPGQAFFATNEIIGDIRQAHC | 35 |
| DQ358757 | C | CCR5 | CTRPNNNTRKSIRIGPGRTFYATGDIIGNIRKAYC | 35 |
| DQ235618 | C | CCR5 | CIRTGNNTRKSVRIGPGQTFYATGAIIGDIRQAHC | 35 |

| | | | | |
|---|---|---|---|---|
| JF896828 | B | CCR5 | CTRPNNNTRKSIHIAPGRAFYATGEIIGNIRAHC | 34 |
| EU744162 | B | CCR5 | CTRPNNNTRRSIHIGPGRAFYTSGGIIGDIRQAHC | 35 |
| EF657924 | B | CCR5 | CIRPNNNTRKSIHMGPGRAFYATGEIIGNIRQAHC | 35 |
| DQ235619 | C | CCR5 | CTRPGNNTRKSVRIGPGQTFFATGDIIGNIRLAHC | 35 |
| FJ375974 | C | CCR5 | CTRPNNNRKSIRIGPGQAFYATGDIIGDIRQAHC | 34 |
| EU575892 | B | CCR5 | CSRPNNNTRRSIHIGPGRAFYATGDITGDIRKAHC | 35 |
| HQ644908 | B | CCR5 | CIRPNNNTRKGIHIGPGRAFYTTGEIIGNIRQAYC | 35 |
| JF896851 | B | CCR5 | CIRPNNNTRKSIHIGPGRAFYATGQIIGNIRQAHC | 35 |
| GU455515 | B | CCR5 | CTRPNNNTRKSIPMGPGKAFYTTGEIIGDIRQAHC | 35 |
| HQ708070 | C | CCR5 | CIRPSNNTRTSMRIGPGQTFYATGDVTGDIRQAYC | 35 |
| DQ222214 | B | CCR5 | CTRPNNNTSKSIPLGPGRAFHTTGRIIGDIRQAHC | 35 |
| DQ002226 | B | CCR5 | CTRPSNNTRKSITIGPGRAFYATGEIIGDVRKAHW | 35 |
| FJ375971 | C | CCR5 | CRGNNTRKSIRIGPGQAFHATGAIIGDIRAHC | 32 |
| AY158535 | C | CCR5 | CIRPGNNTRQSIRIGPGQTFFATGDIIGDIRQALC | 35 |
| AY123262 | C | CCR5 | CTRPNNNTRSVRIGPGQAFYATKDIIGDIRQAHC | 34 |
| DQ061765 | B | CCR5 | CTRPSNNTRKGIHIGPGRAFYATGEIIGDIRKAHC | 35 |
| JF896867 | B | CCR5 | CVRPNNNTRKIHMGPGKAFATGDIIGDIRQAHC | 33 |
| EU578352 | B | CCR5 | CTRPNNNTRRSIHLGAGKALYTGEIIGDIRQAHC | 34 |
| EF117272 | C | CCR5 | CIRPNNNTRKSIRIGPGQTFYATGDIVGDIRQAYC | 35 |
| AF384303 | B | CCR5 | CTRPNNNTRKSIPIGPGRAVYATGQMIGDIRQAHC | 35 |
| KC473827 | B | CCR5 | CTRPGNNTRKSIHLGQGRAWYATGDIIGDIRQAHC | 35 |
| AF286234 | C | CCR5 | CTRPGNNTRKSVRIGPGQTFYTNDIIGDIRQAYC | 34 |
| HQ644797 | B | CCR5 | CTRPNNNTRRSINIGPGRAIYTTGEIIGDIRQAHC | 35 |
| HQ708059 | C | CCR5 | CTRPNNNTRKSVRIGPGQTFYATGAVTGDIRKAYC | 35 |
| EU744141 | B | CCR5 | CTRPNNNTRKSIHIGPGKTFYATGEIIGNIRQAHC | 35 |
| FJ376003 | C | CCR5 | CARPGNNTRKSRIGPGQSFHATGEIIGNIRAHC | 33 |
| HM239618 | B | CCR5 | CIRPGNNTRKSIHIGPGRAFYATGNIIGDIRQAHC | 35 |
| AF384269 | B | CCR5 | CTRPGNNTRRSIHIAPGRAFYATGDIIGDIRKAHC | 35 |
| EU744094 | B | CCR5 | CTRPHNNTRKSIHLGPGRAFYATGDIIGNPRQAYC | 35 |
| HQ645000 | B | CCR5 | CTRPNNNTRRSIRIGPGSAFYTTGDIIGDIRRAHC | 35 |
| FJ376034 | C | CCR5 | CTRPGNNTRSIRIGPGQTFYATIIGIRQAHC | 31 |
| DQ061758 | B | CCR5 | CTRPNNNTRKSISLGPGRAYYATGDIIRNIQAHC | 34 |
| AY010882 | B | CCR5 | CTRPNNNTRTSIPMGLGRAMYATGDIMGNIRQAYC | 35 |
| JF508116 | B | CCR5 | CTRPNNNTRRSIHIGPGRAFFTAGEIIGNIRQAHC | 35 |
| EF117271 | C | CCR5 | CARPSNNTRTSIRIGPGQTFYATGAITGDIRQAHC | 35 |
| AY835451 | B | CCR5 | CTRPGNNTRRSINIGPGRAFYATGAIIGDIRKAHC | 35 |
| EF657902 | B | CCR5 | CIRPNNNTRKSIPMGPGKAFYATGGIIGDIRQAHC | 35 |
| AY170665 | C | CCR5 | CTRPNNNTTTSVRIGPGQTFYATGDIIGNIRAAHC | 35 |
| FJ653219 | B | CCR5 | CTRPSNNTSGSIHIGPGKAFDATKTITGDIRQAHC | 35 |
| AY835446 | B | CCR5 | CIRPNNNTRKGIHIGPGRAFYTTGDIIGDIRQAHC | 35 |
| DQ382367 | C | CCR5 | CTRPSNNTRKSVRIGPGQTFFATGEIIGDIRQAHC | 35 |
| EU575786 | B | CCR5 | CMRPNNNTRKSINIGPGRAFYATGDIIGDIRQAHC | 35 |
| DQ061479 | B | CCR5 | CTGPHNNTRKSIHIGPRRAFYTTEEIIRNIRQAHC | 35 |
| KF770356 | C | CCR5 | CTRPNNNTSQSIRIGPGQTYYAMGRIIGDIRQAHC | 35 |
| EU272334 | B | CCR5 | CTRPNNNTRKSIHIGPGQAWYATGEIIGDIRQAHC | 35 |
| AF384257 | B | CCR5 | CIRPGNNTGKSIPMGPGRAWYATGSIIGDIRQAHC | 35 |
| FJ977095 | C | CCR5 | CTRPNNNTRQSIRIGPGQAFYATGDIIGDIRQAHC | 35 |
| EU272325 | B | CCR5 | CTRPNNNTRKSIHIGPGQAWYTTGQIIGDIRQAHC | 35 |
| U08772 | B | CCR5 | STRPGNNTRKGIPIGPGGSFYATERIIGDIRQAHC | 35 |
| EU744073 | B | CCR5 | CTRYNNNTRKSIPLGPGRAFYATGDIIGDIRQAQC | 35 |
| JF896865 | B | CCR5 | CTRNNNTRRSIPMGPGAFYTTIIGDIRQAHC | 31 |
| EU272269 | B | CCR5 | CIRPNNNTRKSIHLGLGRAWYATGEIIGNIRQAHC | 35 |
| HM368228 | B | CCR5 | CTRPNNNTRRSIHIAPGRTFYTTGEIIGDIRQAHC | 35 |
| DQ061527 | B | CCR5 | CTRPNNNTRKSIHIAPGRAFYATGEIVGDIRQAHC | 35 |
| EU578657 | B | CCR5 | CTRLSNNTRKGVHLGPGSAMYATGEIIGDIRQAHC | 35 |
| AY170659 | C | CCR5 | CIRPNNNTSKSIRIGPGQTFYATGRIIGDIRQAHC | 35 |
| AY510066 | C | CCR5 | CTRPGNNTRKSVRIGPGQAFYATGAIIGDIRQAHC | 35 |
| JX140669 | C | CCR5 | CTRHNNNTRKSIRIGPGQTFYATGDIIGDIRQAYC | 35 |
| AY887857 | C | CCR5 | CTRPGNNTRNSIRIGPGQTFFATGEIIGDIRQAHC | 35 |
| JN687761 | B | CCR5 | CIRPSNNTRKSIHMGPGRVLYATGEIIGDIRQAHC | 35 |
| JF508108 | B | CCR5 | CTRPNNNTRRSIHIGPGRALFTAGEIIGNIRQAHC | 35 |
| AY010884 | B | CCR5 | CTRPNNNTRTSIPIGPGRAIYATGDIIGNIRQAYC | 35 |
| DQ002333 | B | CCR5 | CTRPNNNTRKSIHIAPGSAFYATGDIIGNIRQAYC | 35 |
| AF384264 | B | CCR5 | CTRPNNNTRKGIHVGPGRAFYATGEIIGDIRQAHC | 35 |
| JX140658 | B | CCR5 | CTRPSNNTRKGISLGQGGVFYTTGDIIGNIRQAHC | 35 |
| DQ002103 | B | CCR5 | CTRPSNNTRKSITIGPGRAFYTTEVIGNIRAAYC | 34 |
| AY570012 | B | CCR5 | CTRPNNNTRKSINLGPGRALYTTGEIIGDIRQAHC | 35 |
| JF896823 | B | CCR5 | CTRLNNNTRQSIHMGPGRALYTTDIVGDIRAHC | 33 |
| JF507942 | B | CCR5 | CTRPNNNTRKDIHIGPGRAMYATGQIIGNIRQAHC | 35 |
| AY010774 | B | CCR5 | CLRPNNNTRKSIHMGPGKAFYATGDIIGDIRQAYC | 35 |

| | | | | |
|---|---|---|---|---|
| DQ904338 | C | CCR5 | CTRPNNNTRKSMRIGPGQSFYATGEIIGDIRQAHC | 35 |
| FJ853620 | B | CCR5 | CIRPNNNTRKSINIGPGRAFYAAGEIIGDIRQAHC | 35 |
| KC596066 | B | CCR5 | CVRPNNNTRKSIHLAAGKALYATGDIIGDIRQAHC | 35 |
| DQ904349 | C | CCR5 | CTRPNNNTRKGIGIGPGQTFYATNAIIGDIRQAHC | 35 |
| HQ699979 | B | CCR5 | CTRPNNNTRKSIHLGPGQAWYTTGEIIGDIRQAHC | 35 |
| DQ516159 | B | CCR5 | CTRPNNNTRKDIHIGPGRAFYATGDIIEDIRQAHC | 35 |
| JN687723 | C | CCR5 | CTRPNNNTRKSVRIGPGQAFYATNGIVGNIRQAHC | 35 |
| DQ002059 | B | CCR5 | RIRPNNNTRKSIHIGPGRAFYTTGDVIGDIRQAHC | 35 |
| DQ516121 | B | CCR5 | CIRPNNNTIKSIHIGPGRAFYTTGQIIGNIRQAHC | 35 |
| HM246196 | B | CCR5 | CTRPNNNTRKSIHIAPGRTFYATGDIIGDIRQAHC | 35 |
| HQ644787 | B | CCR5 | CTRPNNNTRRSIHIGPGRALYTTGEIIGDIRQAYC | 35 |
| HQ644903 | B | CCR5 | CTRPGNNTRKSIHIGPGRAFYTTGDITGDIRKAHC | 35 |
| JX140665 | C | CCR5 | CIRPGNNTRKSVRIGPGQAFYATGEIIGDIRKAHC | 35 |
| HQ708022 | C | CCR5 | CTRPNNNTRKSTRIGPGQTFYATGGIIGDIRQAHC | 35 |
| AF259047 | B | CCR5 | CTRPGNNTRKGIHIGPGRAFYTTGQIIGDIRQAHC | 35 |
| HM239564 | B | CCR5 | CTRPNNNTRKSINIGPGRAFYATGAIIGDIRQAHC | 35 |
| AY158534 | C | CCR5 | CTRPHNNTRKSVRIGPGQAFYATGDIIGNIRQAHC | 35 |
| DQ002178 | B | CCR5 | CTRPNNNTRRSITIGPGRAFYGTDIIGDIRQAHC | 34 |
| JN002060 | B | CCR5 | CTRPNNNTRKSIPIGPGRAIYTTGGIIGDMRQAHC | 35 |
| AF153175 | C | CCR5 | CTRPNNNTRKSVRIGPGQAFYATNEIIGDIRQAHC | 35 |
| FJ653109 | B | CCR5 | CTRPNNNTRKSINIGPGGAFYAATDIIGDIRQAHC | 35 |
| HM239565 | B | CCR5 | CTRPNNNTRKGVHIGPGRTFFYTGDIIGDIRQAHC | 35 |
| HQ644834 | B | CCR5 | CTRPNNNTRRSINIGPGRAIYTTGQIIGDIRQAHC | 35 |
| EF600087 | B | CCR5 | CIRPNNNTRQGIHIGPGKALYTTNIIGNIRQAHC | 34 |
| DQ904337 | C | CCR5 | CTRPNNNTRKSIRIGPGQTFYATGAIIGNIREAHC | 35 |
| JF507850 | B | CCR5 | CTRPGNNTRRSINIGPGRAFYTGEIVGDIRQAHC | 34 |
| AY835452 | B | CCR5 | CTRPNNNTSKSITIGPGRAFYATGRIIGDIRKAHC | 35 |
| AY887870 | C | CCR5 | CIRPGNNTRKSVRIGPGQAFYATGDIIGDIRKAHC | 35 |
| DQ002066 | B | CCR5 | CIRPNNNTRKSIHIGPGRVFYTTGDIIGDIRQAHC | 35 |
| HM239575 | B | CCR5 | CTRPNNNTRKSIPMGPGKAWYATGEIIGDIRQAYC | 35 |
| HM179917 | C | CCR5 | CTRPNNNTRKSIRIGPGQTFYATGDIMGDIRQAYC | 35 |
| DQ002326 | B | CCR5 | CTRPSNNTRKSIHMGPGRAFYTTGEIIGDIRQTHC | 35 |
| HQ678274 | B | CCR5 | CTRPNNNTRKGIHIGPGRVFYATGEIIGDIRQAHC | 35 |
| HQ644864 | B | CCR5 | CTRPGNNTRKSIHMGPGRAFYVTEDVIGDIRQAHC | 35 |
| DQ235615 | C | CCR5 | CIRPNNNTRQSVRIGPGQTFFANDIIGDIRQAHC | 34 |
| HM246240 | B | CCR5 | CTRPNNNTRKGVHIGPGRAFYATGIIGDIRKAYC | 34 |
| AY887885 | C | CCR5 | CTRPGNNTRKSVRIGPGATFYATGDIIGDIRQAHC | 35 |
| EU575529 | B | CCR5 | CTRPNNNTRKGIHIGLGRALYATGDIIGDIRQAHC | 35 |
| DQ061416 | B | CCR5 | CTRPNNNIRKSINIGPERAFYTTGEIIGDIRQAHC | 35 |
| AY887861 | C | CCR5 | CTRPGNNTRKSIRIGPGQAFYATGDIIGDIRKAHC | 35 |
| EU578397 | B | CCR5 | CTRPNNNTRRSIHMGAGKALYTGEIIGDIRQAHC | 34 |
| EU578272 | B | CCR5 | CTRPNNNTRKSIHMGWGRAFYATGQIIGDIRQAHC | 35 |
| EF688454 | B | CCR5 | CTRPNNNTRRSIHIGPGRAFHATGEIVGNIRQAHC | 35 |
| FJ653100 | B | CCR5 | CTRPNNNTRKSINIGPGRAFYAATGIIGDIRQAHC | 35 |
| DQ061684 | B | CCR5 | CIRPNNNTRKGIHLGPGGAFYATGGIIGGIRQAYC | 35 |
| AY736823 | C | CCR5 | CTRPNNNTRRSIRIGPGQTFYATGEIIGDIRQAHC | 35 |
| AJ418514 | B | CCR5 | CTRPNNNTRKSIHIGPGRAFYTTGGIIGNIRQAHC | 35 |
| DQ382365 | C | CCR5 | CTRPGNNTRKSVRFGPGQAFYATGDIIGDIRQAHC | 35 |
| JQ779237 | C | CCR5 | CTRPNNNTRKSIRIGPGQAFFATTGIIGNIRQAYC | 35 |
| FJ376035 | C | CCR5 | CIRPNNNTKSRIGPGQFATDIIGNIRQAHC | 30 |
| AY170666 | C | CCR5 | CARPNNNTRTSVRIGPGQAFYATNDIIGKIRQAHC | 35 |
| AF153165 | C | CCR5 | CIRPGNNTRKGMRIGPGQTFYATGDIIGDIRQAHC | 35 |
| DQ382363 | C | CCR5 | CTRPNNNTRKSVRIGPGQTFYATGEIIGNIRQAHC | 35 |
| DQ061805 | B | CCR5 | CTRPSNNTRKGIHIGPGRAFYATGAIIGNIRQAHC | 35 |
| KF384807 | B | CCR5 | CTRPSNNTIKGIHMGPGRAFYATEQVIGDIRQAHC | 35 |
| AY505002 | C | CCR5 | CTRPGNNPRKSVRIGPGQAFYATGDIIGDIRQAYC | 35 |
| DQ002211 | B | CCR5 | CTRPNNNTRKSIHIGPGRAFYTTGEIIGDIRQAYC | 35 |
| AY010857 | B | CCR5 | CTRPNNNTGTSIPMGPGRAVYATGDIIGNIRHAYC | 35 |
| JF507914 | B | CCR5 | CTRPNNNTRKSISIGPGRAFYATGQIIGDIRQAHC | 35 |
| DQ358770 | C | CCR5 | CTRPNNNTRKSIRIGPGQTFYATRGIIGDIREAHC | 35 |
| HQ644888 | B | CCR5 | CTRPNNNTRKGIHIGPGRAFYTTGQIIGNIRLAHC | 35 |
| KF770311 | C | CCR5 | CTRPDNNTRRSVRIGPGQVFYTNDIIGDIRQAYC | 34 |
| AF153188 | C | CCR5 | CTRPGNNTRTSIRIGPGQTFFANNIIGDIRQAHC | 34 |
| DQ516305 | B | CCR5 | CTGPNNNTRKSIHIGPGRAFYTTGGIIGDIRQAHC | 35 |
| FJ376025 | C | CCR5 | CTRPGNNKRKSMRIGPGQTFYATGDVIGDIRKAQC | 35 |
| AF384321 | B | CCR5 | CTRPNNNTRKGIHIGPGGAIYATGAITGDIRQAHC | 35 |
| DQ002311 | B | CCR5 | CTRPNNITRKSINIGPGRAWYTTGAIIGDIRKAHC | 35 |
| AY842837 | B | CCR5 | CIRPNNYTRKSINIGPGRAMYATEQITGDIRQAHC | 35 |
| HM239603 | B | CCR5 | CTRPNNNTRKGVQVGPGKALYITGSIIGDIRQAHC | 35 |
| DQ235639 | C | CCR5 | CIRPNNNTRKSIRIGPGQAFYATNDIIGDIRQAHC | 35 |

| | | | | |
|---|---|---|---|---|
| AY835443 | B | CCR5 | CERPNNNTIKSIHLGPGRAWHATGQIIGDIRKAFC | 35 |
| AF384289 | B | CCR5 | CTRPNNNTRKSIIGPGRAFYATGDIIGDIRQAHC | 34 |
| EU744063 | B | CCR5 | CTRPHNNTRRSIHLGPGKTFYATGDIIGNIRQAHC | 35 |
| EU272196 | B | CCR5 | YKTPQQSRKSINLGPGRAWYTTGQIIGDIRQAHC | 34 |
| EF688428 | B | CCR5 | CTRPNNNTRKGIHIGPGRAFYTTGEIIGNIRQAHC | 35 |
| AF384293 | B | CCR5 | YTRPNNNTRKSIPIGPGRAFYARGDIIGDIRQAHC | 35 |
| DQ061821 | B | CCR5 | CTRPNNNTRKGIHMGPGAVFYATGQIIGDIRKAHC | 35 |
| AF153150 | C | CCR5 | CTRPGNNTRKSVRIGPGQAFYATGEIIGDIRQAHC | 35 |
| DQ177200 | B | CCR5 | CTRPNNNTRKGIHMGPGRAYYATGDIIGNIRQAHC | 35 |
| DQ002101 | B | CCR5 | CTRPSNNTRRGIHIGPGKAFYTTGGVIGDIRKANC | 35 |
| EU293446 | C | CCR5 | CIRPNNNTRKSIRIGPGQSFHATGEIIGNIRQAHC | 35 |
| HM239627 | B | CCR5 | CTRPNNNTRKIHIGPGRAFYATGDIGDIRQAHC | 33 |
| AF384245 | B | CCR5 | CTRPNNNTRKSIHIAPGRAYYATGDIIGDIRQAHC | 35 |
| DQ061750 | B | CCR5 | CTRPNNNTRRSISLGPGRAYYATGDIIGNIRQAHC | 35 |
| JF896849 | B | CCR5 | CTRPNNTRKSIPMGPGKVFYATEIIGDIRQAHC | 33 |
| KF384813 | B | CCR5 | CTRPNNNTRKSISIGPGRAFYTTGEVIGDIRQAHC | 35 |
| FJ376027 | C | CCR5 | CTRPNNNTRKSVRIGPGQTFYATGDIIGDRQAYC | 34 |
| EU578380 | B | CCR5 | CTRPGNNTRKGITIGPGSVFYTGEIIGDIRQAHC | 34 |
| AF180901 | B | CCR5 | CTRPNNNTRKSIPIGPGRAFYATGQIIGDIRQAYC | 35 |
| DQ061431 | B | CCR5 | CTRPNNNTRKSIHIGPGRAFYTAGEIIGDIRQAHC | 35 |
| AF254775 | C | CCR5 | CTRPNNNTRKSVRIGPGQTFYATGEIIGNIREAHC | 35 |
| DQ002194 | B | CCR5 | CTRPGNNTRKSIRIGPGRAFYATDIIGDIRQAYC | 34 |
| HM179921 | C | CCR5 | CTRPNNNTRKSIRIGPGQAFYATGDIIGDIRQAYC | 35 |
| JF507899 | B | CCR5 | CTRPNNNTRQSVHIGPGRALYTTNIIGDIRKAYC | 34 |
| AY887871 | C | CCR5 | CTRPANNTRRSIRIGPGQTFYATGEIIGDIRQAHC | 35 |
| JN687759 | B | CCR5 | CTRPNNNTRRSIPMGPGRMFTTKIVGDIRQAHC | 33 |
| HQ644835 | B | CCR5 | CTRPNNNTRRSITIGPGRAFYTTGDIIGDIRQAHC | 35 |
| JN002057 | B | CCR5 | CTRPDNNTRKGIHLGPGGTFFATGAKIGDIRQAHC | 35 |
| AF384323 | B | CCR5 | CTRPNNNTRKGIHTGPGGAIYATGAIIGDIRQAHC | 35 |
| EU272266 | B | CCR5 | CTRPNNNTRKSINLGPGRAWYATGQIIGDIRRAHC | 35 |
| HM239633 | B | CCR5 | CIRPNNNTIKSIPIGPGRAFYATGKIVGDIRKAYC | 35 |
| AM156921 | B | CCR5 | CIRPNNNTRRSIHIGPGRAFYTTDIIGNIRQAHC | 34 |
| AF541009 | B | CCR5 | CTRPSNNTRKSIPIGPGRAFYATGEITGDIRKAHC | 35 |
| EF657925 | B | CCR5 | CMRPNNNTRKSIPMGPGRAFYATGEVIGNIRQAHC | 35 |
| EU786679 | B | CCR5 | CTRPNNNTRRSITIGPGRAFYATGEIIGDIRKAYC | 35 |
| AF355729 | B | CCR5 | CTRPCNNTRKSIPLGPGRAFYATGDIIGNIRKAHC | 35 |
| HQ644791 | B | CCR5 | CTRPNNNTRRSIHIGPGRAFYTTGEIIGDIRQAHC | 35 |
| DQ061400 | B | CCR5 | CTGPNNNTRKSIHIGPGRAFYTTGEIIGDIRQAHC | 35 |
| HM246237 | B | CCR5 | CTRPNNNTRKSINLGPGRTIYATGDIIGDIRQAHC | 35 |
| EU786678 | B | CCR5 | CIRPNNNTRKSINIGPGRAFYTTGAITGDIRQAHC | 35 |
| JF896817 | B | CCR5 | CIRPNNNTRKSIHMGPGRAYYTTDIIGDIRKAYC | 34 |
| AF384252 | B | CCR5 | CTRPNNNTRKSIHMGPGRAFYTTGEIIGNIRQAHC | 35 |
| U08714 | B | CCR5 | CTRPSNNTRKGIHIGPGRAFYATGDIIGDIRQAHC | 35 |
| AF199033 | B | CCR5 | CTRPSNNTKSIHIGPGRAFYTTGEITGDIRQAHC | 34 |
| EU744118 | B | CCR5 | CTRPNNNTRKSINIGPGRAIYTTGEIIGNIRQAHC | 35 |
| AF082384 | B | CCR5 | CTRLNNYTKEVVSMGPGRAFFTTGDIIGAIRRAHC | 35 |
| DQ061523 | B | CCR5 | CTRPNNNTRKSIPIGPGRAFYATGETIGDIRQAHC | 35 |
| AF384254 | B | CCR5 | CTRPSNNTRKSIHMGPGRAWYATGSITGDIRQAHC | 35 |
| EF688437 | B | CCR5 | CTRPNNNTRKSIYIGPGRAFHTTGRIIGDIRQAHC | 35 |
| AF153154 | C | CCR5 | CTRPNNNTRKSIRIGPGQAFYATGDIIGDIRQAHC | 35 |
| EF688448 | B | CCR5 | CTRPNNNTRKSIPIGPGRAFYTTGDIIGNIRQAHC | 35 |
| GU945314 | C | CCR5 | CTRPGNNTRKSIRIGPGQTFYATGGIIGDIRQAHC | 35 |
| AF153132 | C | CCR5 | CTRPGNNTRKSVRIGPGQTFYATDDIIGDIRKAHC | 35 |
| AF153179 | C | CCR5 | CTRPNNNTRQSMRIGPGQTFYATGDIIGNIRQAHC | 35 |
| DQ061398 | B | CCR5 | CTRPNNNTRKSIHIGPGSAFYTTGEIIGNIRQAHC | 35 |
| DQ002301 | B | CCR5 | CTRPSNNTRKGIHIGPGRAFYATGDIIGDIRKAHC | 35 |
| JF508035 | B | CCR5 | CTRPNNNTRKGIHIGPGRAFYTTGEIIGNIRQASC | 35 |
| KF770342 | C | CCR5 | CTRPGNNTRRSIRIGPGQSFYATGGIIGDIRQAHC | 35 |
| AY170657 | C | CCR5 | CTRPSNNTRKSIRVGPGQSFHATGEIIGDIRQAHC | 35 |
| AY010894 | B | CCR5 | CIRPNNNTRKSIPMGPGKAFYATGDIIGDMRDACW | 35 |
| AY842786 | B | CCR5 | CTRPNNNTRKSIHMGPGRVLYTTGGITGDIRQAHC | 35 |
| AF199040 | B | CCR5 | CTRPNNNTRKSINMGPGRAWYTTGQIIGDIRQAHC | 35 |
| FJ977083 | C | CCR5 | CIRPGNNTRKSIRIGPGQTFFATGDIIGDIRRAHC | 35 |
| DQ177191 | B | CCR5 | CTRPNNNTRKSIHIAPGRAFHATGDIIGNIRQAHC | 35 |
| DQ002302 | B | CCR5 | CTRPNNNTRKGIHIGPGKAFYATGDIIGDIRKAHC | 35 |
| AF355732 | B | CCR5 | CTRPNNNTRKSIHLGPGRAFYATGDIIGNIRKAHC | 35 |
| KF384811 | B | CCR5 | CERPNNNTRRSIPIGPGRVFFTSEIIGDIRQAYC | 34 |
| JF896820 | B | CCR5 | CTRPNNNTRKSIHIAPGRSFYATGDIIGDIRQAHC | 35 |
| FJ375978 | C | CCR5 | CTRPNNNTRKSVRIGPGQTFYAGGIIGDIRQAYC | 34 |
| HM239546 | B | CCR5 | CTRPNNNTRKSIHITPGRAFYATGDIIGDIRQAHC | 35 |

| | | | | |
|---|---|---|---|---|
| FJ375967 | C | CCR5 | CTPNNNTSVRIGPGQTFYATNDIIGNIRQAC | 31 |
| DQ235647 | C | CCR5 | CTRPNNNTRQGIGIGPGQTFYAHTNIIGDIRQAHC | 35 |
| HQ644794 | B | CCR5 | CTRPNNNTRRSIHIGPGSAFYTGEIIGNIRQAHC | 34 |
| HQ678273 | B | CCR5 | CTRPSNNTSKSIPIGPGRAFYATGRIIGDIRQAHC | 35 |
| U45485 | C | CCR5 | CTRPSNNTRKSMRIGPGQTFYATGDIIGDIRQAHC | 35 |
| DQ235638 | C | CCR5 | CTRPGNNTRKSIRIGPGQTFYANDIIGDIRQAYC | 34 |
| DQ061439 | B | CCR5 | CTRPSNNTRKSIHIGPGRAFYTTGEVIGDIRQAHC | 35 |
| HQ644925 | B | CCR5 | CVRPNNNTRKSIPIGPGRAFYTTGDIIGDIRQAHC | 35 |
| HM239637 | B | CCR5 | CTRPNNNTRKSISIGPGRAFFATGDIIGDIRQAHC | 35 |
| DQ061802 | B | CCR5 | CIRPNNNTRKGIHIGPGRAFYATGAIIGNIIQAHC | 35 |
| DQ516111 | B | CCR5 | CTRPNNNTVKSIHIGPGRAFYTTGQIIGNIRQAHC | 35 |
| HM246221 | B | CCR5 | CTRPNNNTRKSIIGPGRAYFATGDIIGDIRKAHC | 34 |
| AY529663 | C | CCR5 | CPRPNNNTRKSIRIGPGQTFYATNDIIGDIRQAHC | 35 |
| AF310108 | B | CCR5 | CTRPNNNTRKSIHIEAGKALYTGEIIGDIRQAHC | 34 |
| DQ061436 | B | CCR5 | CARPNNNTRKSIHIGPGRAFYTTGEIIGDIRQAHC | 35 |
| AY713410 | B | CCR5 | CTRPNNNTRKGIHMGPGRAFYATGTIIGDIRQAHC | 35 |
| U08671 | B | CCR5 | CTRPGNNTRKGIPIGPGGSFYATERIIGDIRQAHC | 35 |
| DQ002219 | B | CCR5 | CTRPSNNTRKSITIGPGRAFYATGDIIGDIRKAHC | 35 |
| DQ235648 | C | CCR5 | CTRPGSNTRRSIRIGPGQAFYTQDIIGDIRQAHC | 34 |
| AF541036 | B | CCR5 | CTRPSNNTRKSIPMGPGKAFYTTGEIIGNIRQAHC | 35 |
| DQ002227 | B | CCR5 | CARPSNNTRKSITIGPGRAFYATGDIIEDIRKAHC | 35 |
| KF770455 | C | CCR5 | CTRPGNNTRRSVRIGPGQAFYTTGEIIGDIRVAHC | 35 |
| AF153170 | C | CCR5 | CTRPNNNTRQSIRIGPGQVFYATGDIIGDIRQAHC | 35 |
| AF153129 | C | CCR5 | CTRPNNNTRKSIRIGPGQTFYATGGIIGNIREAHC | 35 |
| KF716495 | B | CCR5 | CTRPSNNTMKSIHIGPGRAFYTTEQVIGDIRKAHC | 35 |
| AY170667 | C | CCR5 | CTRPGNNTRKSVRIGPGQAFYATGDIIGDIRKAYC | 35 |
| U08688 | B | CCR5 | CTRPNNNTKSIHIGPGSAFYATGDIIGDIRQAHC | 34 |
| HM239527 | B | CCR5 | CTRPNNNTRKSIQGPGKAIYATGEIIGDIRKAHC | 34 |
| AY842809 | B | CCR5 | CTRPNNNTRKSIHMGPGRAMYATGDIIGDMRQAHC | 35 |
| DQ002104 | B | CCR5 | CTRPNNNTRRSITIGPGKAFYTTGGVIGDIRKANC | 35 |
| EU575611 | B | CCR5 | CTRPNNNTRKSIHIGPGGAFYAAGGIIGNIRQAHC | 35 |
| KC312573 | B | CCR5 | CERPNNNTRKSIRIGPGSAFYAAGEIIGNIRQAHC | 35 |
| EU575279 | B | CCR5 | CTRPNNNTIKGIHIGPGRAFYTTGQVIGDIRKAYC | 35 |
| AF082379 | B | CCR5 | CTRPNNNTRKSIPMGPGRAFYATGQIIGDIRQAYC | 35 |
| HM239614 | B | CCR5 | CTRPNNNTRRSVHFAPGRAFYATGDIIGDIRQAHC | 35 |
| JF507766 | B | CCR5 | CTRPNNNTRKGIHIGPGRVFYATEGIIGDIRRAYC | 35 |
| JQ779247 | C | CCR5 | CIRPNNNTRKSIRIGPGQAFFATTGIIGNIRQAYC | 35 |
| HM246194 | B | CCR5 | CTRPSNNTRRGIHMAGRAFYATGQIIGDIRQAHC | 34 |
| DQ002327 | B | CCR5 | CTRPNNNTRRSIHMGPGKAFYTTGEIVGNIRQAYC | 35 |
| AY887877 | C | CCR5 | CTRPGNNTRKSIRIGPGQTFYATGDIIGDIRRAYC | 35 |
| DQ382366 | C | CCR5 | CTRHNNNTRKSVRIGPGQTFYATGDIIGDIRQAHC | 35 |
| AF180906 | B | CCR5 | CTRPNNNTRKSINIGPGRAFYTTGQVIGDIRKAHC | 35 |
| DQ061753 | B | CCR5 | CTRPNNNTRRSISLGPGRAYYATGDIVGDIRQAHC | 35 |
| FJ375996 | C | CCR5 | CTRPNNNTRRGIGIGPGQTFFATDAIIGDIRQAHC | 35 |
| JF507980 | B | CCR5 | CTRPNNNTRKSINIGPGRAFYTTGAIIGNIRQAHC | 35 |
| AY010876 | B | CCR5 | CTRPNNNTRTSIPMGSGRAMYATGDIIGNIRQAYC | 35 |
| DQ869016 | B | CCR5 | CTRPNNNTRKSISIAPGRAWYATGDIIGDIRQAHC | 35 |
| FJ798398 | B | CCR5 | CTRPNNNTRKGISIGPGRAFYATGGIIGDIRKAHC | 35 |
| JF508097 | B | CCR5 | CTRPNNNTRKGIHIGPGRTFHVTGEIIGDIRQAHC | 35 |
| HQ678266 | B | CCR5 | CSRPNNNTRKSIHIAPGRTFYATGDIIGDIRQAHC | 35 |
| HM239535 | B | CCR5 | CVRPNNNTRRSIPLGPGKTFYAGEVIGDIRQAHC | 34 |
| DQ235631 | C | CCR5 | CIRPGNNTSKSIRIGPGQTFYATGDVIGNIRQAHC | 35 |
| HM246195 | B | CCR5 | CTRPNNNTRKSIPMGPGAAIYATGAIIGDIRQAHC | 35 |
| FJ977086 | C | CCR5 | CTRPGNNTRKSVRIGPGQTFYATGDIIGNIRQAHC | 35 |
| EU272248 | B | CCR5 | CIRPNNNTRKSIPIGPGRAWYTTGQIIGDIRQAHC | 35 |
| JF507750 | B | CCR5 | CTRPNNNTRKSIPIGPGRVFFATGGIIGDIRRAHC | 35 |
| JX140656 | B | CCR5 | CTRPNNNTRKGIQMGPGRAFYATGDIIGDIRQAHC | 35 |
| HQ644790 | B | CCR5 | CTRPNNNTRRSIHIGPGRAFYTTGEIVGDIRQAHC | 35 |
| GU204921 | B | CCR5 | CRPNNNTRKSIHLTPGAFHTTGSIIGDIRAC | 31 |
| JF507983 | B | CCR5 | CTRPNNNIRKSINIGPGRAFYTTGAIIGDIRQAHC | 35 |
| DQ516307 | B | CCR5 | CTRPNNNTRKSIHIGPGRALYTTGGIIGDIRQAHC | 35 |
| JN002043 | B | CCR5 | CTRPNNNTRKSIPIGPGRAFYATGDIIGNIRQAHC | 35 |
| HM215423 | B | CCR5 | CTRPNNNTLKSIQLGLGRAWHATGQIIGDIRQAHC | 35 |
| AY426113 | B | CCR5 | CTRPNNNTRKSIHIGPGRKIYTTGKIIGDIRQAHC | 35 |
| DQ002241 | B | CCR5 | CTRPNNNTRKSIHIAPGRAFYATGEIIGDIGQAHC | 35 |
| HQ644897 | B | CCR5 | CTRPNNNTRKSIHIGPGRAFYTTGDIIGDIRKAYC | 35 |
| HM239593 | B | CCR5 | CTRPNNNTRRSVPLGPGRAVYATGAIIGDIRQAYC | 35 |
| DQ002087 | B | CCR5 | CTRPNNNTRKSIHIGPGRAFYATGNIIGNIRQAHC | 35 |
| U08689 | B | CCR5 | CTRPGNNTRRSIHMGWGRAFYATGDIIGDIRQAHC | 35 |
| AY887864 | C | CCR5 | CIRPGNNTRKSIRIGPGQTFYATGDIIGDIRKAHC | 35 |

| | | | | |
|---|---|---|---|---|
| DQ382375 | C | CCR5 | CTRPNNNTRRSIRIGPGQTFYTNDIIGDIRQAYC | 34 |
| AF541084 | B | CCR5 | CVRPNNNTRRGIHIGLGRRFYTTEIVGDIRRAYC | 34 |
| DQ388515 | C | CCR5 | CVRPNNNTRKSVRIGPGQTFFATGEIIGDIRQAHC | 35 |
| JN002040 | B | CCR5 | CTRPNNNTRKSIHMGPGKAFYATGDIIGDMRRAHC | 35 |
| HM239605 | B | CCR5 | CTRPNNNTRRSIHIAPGKAFYATGDVIGDIRQAHC | 35 |
| DQ516215 | B | CCR5 | CTRPNNNTRKGIHIGPGSAFYTTGAIIGNIRQAHC | 35 |
| KC156330 | C | CCR5 | CARPHNNTRKSMRIGPGQAFYATGDVTGDIRQAHC | 35 |
| DQ516225 | B | CCR5 | CTRPNNNTRKGIHIGPGRAFYTTGAIIGSIRQAHC | 35 |
| AY669749 | C | CCR5 | CTRPNNTRKSIRIGPGQTFYATNEIIGNREAHC | 33 |
| DQ061848 | B | CCR5 | CTRPNNNTRKGIHMGPGKMFYATGQIIGDIRQAYC | 35 |
| EU576774 | B | CCR5 | CTRPNNNTRKSIPIGPGSVFYTGEIIGDIRQAHC | 34 |
| EF117274 | C | CCR5 | CTRPGNNTRKSIRIGPGQTFYATGEIIGNIRQAHC | 35 |
| JQ779236 | C | CCR5 | CTRPNNNTRQSIRIGPGQAFFATTGIIGNIRQASC | 35 |
| EU744170 | B | CCR5 | CTRPNNNTRRSIHIGPGRAIYATGDIIGDIRKAHC | 35 |
| HQ708057 | C | CCR5 | RIRPNNNTRKSVRIGPGQTFYATGGIIGDIRRAYC | 35 |
| JX140659 | B | CCR5 | CTRPNNNTRKGIHIGPGKTFFATEVIGDIRKAHC | 34 |
| JF507793 | B | CCR5 | CMRPGNNTRKSITIGPGRAFYAGEIIGNIRQAHC | 34 |
| JN188292 | C | CCR5 | CTRPGNNTRKSVRIGPGQTFFATGEIIGKIREAHC | 35 |
| EU577152 | B | CCR5 | CTRPNNNTRRSITFGPGAAFYTGEIIGDIRQAYC | 34 |
| JF896831 | B | CCR5 | CTRPGNNTRKSIHMGPGRAFYTTGEIIGDIRLAHC | 35 |
| DQ177210 | B | CCR5 | CTRPNNNTRRSITIGPGRAFYATGDIIGDIRQAHC | 35 |
| JX140667 | C | CCR5 | CTRPHNNTRKSVRIGPGQTFYATGEIIGNIRQAHC | 35 |
| JF507803 | B | CCR5 | CMRPGNNTKKSITIGPGKAFYAGEIIGDIRKAHC | 34 |
| KF770389 | C | CCR5 | CTRPGNNTRKSMRIGPGQTFYATGDIIGNIRQAHC | 35 |
| HM239524 | B | CCR5 | CTRPSNNTRKSIPMGPGKAFYATGEIVGDIRQAHC | 35 |
| AF153180 | C | CCR5 | CTRPNNNTRRSYGIGPGQAFRATTNIIGDIRKAHC | 35 |
| HM239576 | B | CCR5 | CTRPNNNTRKGIHMGPGGAFYATGEIIGNIRQAHC | 35 |
| EU744066 | B | CCR5 | CIRPNNNTRKSIHLGPGRAFYATGDIIGDIRQAHC | 35 |
| AF384332 | B | CCR5 | CIRPNNNTRKGIHIGPGRTFYGTGEIIGNIRQAHC | 35 |
| AB553914 | B | CCR5 | CTRPNNNTRKGIHFGPGQALYTTGAIIGDIREAHC | 35 |
| EF657907 | B | CCR5 | CIRPSNNTRKSIPMGPGRAFYATGDITGNIRQAHC | 35 |
| HQ644867 | B | CCR5 | CTRPGNNTRKSIHMGPGRAFFVTDDVIGDIRQAHC | 35 |
| FJ977079 | C | CCR5 | CTRPGNNTRTSIRIGPGQTFYATGDIIGDIRQAHC | 35 |
| KF716466 | C | CCR5 | CTRPNNNTRKNVRIGPGQAFYATNGIIGDIRQAYC | 35 |
| AF541024 | B | CCR5 | CTRPNNNTRKGINIGPGRAFYTTGEIIGDIRQAHC | 35 |
| KC473831 | B | CCR5 | CTRPNNNTRRSIHLGPGKAIYTTGEIIGDIRRAHC | 35 |
| JF896842 | B | CCR5 | CVRPNNNTRKGIHIGPGRAFYATGEIIGNIRQAHC | 35 |
| HQ645001 | B | CCR5 | CTRPNNNTRRSITMGPGKAFYTTGDIIGDIRRAHC | 35 |
| U04925 | B | CCR5 | CTRPSNNTRKSVHIGPGRAFYTTGEIIGDIRQAHC | 35 |
| EF688440 | B | CCR5 | CTRPHNNTRKGIHIGPGRAFYTAEKIVGDIRQAHC | 35 |
| DQ061683 | B | CCR5 | CIRPNNNTRKGIHLGPGGAFYATGGIIGDIRQAYC | 35 |
| AF391233 | C | CCR5 | CTRPNNNTRKSIRIGPGQTFYATNGIIGNIRQAHC | 35 |
| EU575091 | B | CCR5 | CTRPNNNTRKSIGMGPGRAFYATGDIIGDIRQAHC | 35 |
| AF254766 | C | CCR5 | CTRPNNNTRKSVRIGPGQAFYATNDVIGDIRQAHC | 35 |
| HQ644852 | B | CCR5 | CTRPGNNTRKSITIGPGRAFYTTGDIIGDIRQAHC | 35 |
| DQ061478 | B | CCR5 | CTRPNNNTGKSISIGPGRAFYTTGEIIGEIRQAYC | 35 |
| AF254778 | B | CCR5 | CTRPNNNTRKSINLGQGRAWYATGAIIGDIRQAHC | 35 |
| DQ002092 | B | CCR5 | CTRPSNNTRRGIHIGPGRAFYTTGEVIGDIRAANC | 35 |
| AF384296 | B | CCR5 | CTRPNNNTKKSIHIGPGRAVYATGQIIGDIRQAHC | 35 |
| HQ678286 | B | CCR5 | CTRPNNNTRKSIHMGPGGAFYATGDIIGDIRQAHC | 35 |
| AY842824 | B | CCR5 | CIRPNNNTRKSINIGPGRAMYATEQITGDIRQAHC | 35 |
| EU744082 | B | CCR5 | CTRLNNNTRKSIPLGPGRAFYATGDIIGDIRKAQC | 35 |
| EF579978 | B | CCR5 | CERPGNNTSKGIHIGPGRAFCATENIIGDIRKAHC | 35 |
| U04908 | B | CCR5 | CTRPNNNTRKGIHIGPGRAFYTTGEVIGNIRQAHC | 35 |
| U04917 | B | CCR5 | CIRPNNNTRKSIHIGPGRAFYTTGDIIGDIRKAHC | 35 |
| AF384258 | B | CCR5 | CIRPSNNTRKSIPMGPGRAWYATGSIIGDIRQAHC | 35 |
| AF384282 | B | CCR5 | CTRPNNNTRKSIHIGPGRAFYTTGQIVGDIRKAHC | 35 |
| EU578561 | B | CCR5 | CTRPNNNTRRSIHMGPGKALYTGDIIGDIRQAHC | 34 |
| AB553912 | B | CCR5 | CTRPNDNTRKSINIAPGRAFYATGDIIGDIRQAHC | 35 |
| JF896824 | B | CCR5 | CTRPNNNTRRSISIGPGRAFFATGEVIGDIRKAYC | 35 |
| KC156277 | C | CCR5 | CTRPNNNTRKSMRIGPGQTFYATGDIIGNIRQAHC | 35 |
| DQ869019 | B | CCR5 | CTRPSNNTRKGIHIGPGRAFYTTGDIIGDIRQAHC | 35 |
| AY887891 | C | CCR5 | CTRPSNNTRTSIRIGPGQTFYATGEIIGDIRKAHC | 35 |
| AF153145 | C | CCR5 | CTRPNNNTRKSIRIGPGQTFFATNAIIGDIRQAHC | 35 |
| DQ061540 | B | CCR5 | CIRPNNNTRKSIHIAPGRAFYATGEIIGDIRQAHC | 35 |
| KC312477 | B | CCR5 | YKTQQQYKERYTYRTRENIYATGEIIGDIRQAHC | 34 |
| EU744166 | B | CCR5 | CTRPNNNTRKGIHIGPGRAFYTSGGIIGDIRQAHC | 35 |
| AF384273 | B | CCR5 | CTRPNNNTRKSIPMGPGKFYATGEIIGNIRQAHC | 34 |
| JF896844 | B | CCR5 | CTRPNNNTRKGIHMGPGRAFYATEIIGNIRQAHC | 34 |
| JN002009 | B | CCR5 | CTRPHNTRKSIHIGPGRAFYATGDIVGDIREAHC | 34 |

| | | | | |
|---|---|---|---|---|
| EU575538 | B | CCR5 | CTRPSNNTRKSIHMGPGGAFYATGSIIGDIRQAHC | 35 |
| FJ375981 | C | CCR5 | CTRPNNNTRKSRIGPGQTFYATGDIIGDIRQAHC | 34 |
| KF770313 | C | CCR5 | CIRPDNNTRRSIRIGPGQVFYANDIIGDIREASC | 34 |
| AY887855 | C | CCR5 | CTRPSNNTRKGIRIGPGQTFYATGDVIGDIRQAHC | 35 |
| KC156320 | C | CCR5 | CNRPHNNTRKSMRIGPGQAFYATGDVTGDIRKAHC | 35 |
| AY835435 | B | CCR5 | CTRPNNNTRKSIHMGPGKVFYTTGEIIGDIRQAHC | 35 |
| DQ002118 | B | CCR5 | CTRPNNNTRKSITIGPGGAFYATGDIIGDIRQAHC | 35 |
| AY887868 | C | CCR5 | CTRPNNNTRQSVRIGPGQTFYATNIIGDIRQAYC | 34 |
| JF507855 | B | CCR5 | CTRPGNNTRRSINIGPGRAFYTGEMIGDIRQAHC | 34 |
| HQ644839 | B | CCR5 | CIRPNNNTRRSINIGPGRAFYTTGDIIGDIRQAHC | 35 |
| HM239539 | B | CCR5 | CTRPNNNTRRSIPIGPGRAFWATGDIIGDIRQAHC | 35 |
| EU575025 | B | CCR5 | CTRPNNNTRKGIHIGPGRAFYATGQIIGDIKRAYC | 35 |
| AY835438 | B | CCR5 | CTRPNNNTRKSINLGPGRAFYATGDIIGDIRQAHC | 35 |
| JF508106 | B | CCR5 | CTGPNNNTRKSIHIGPGRTFYATGEIIGDIRQAHC | 35 |
| DQ002170 | B | CCR5 | CTRPNNNTRRSITIGPGGAFYATDIIGDIRQAHC | 34 |
| FJ670524 | B | CCR5 | CIRPNNNTRKSIHIGPGRAFYAQDIIGDIRQAHC | 34 |
| EU272300 | B | CCR5 | CTRPNNNTRKSIHLGLGRAWYATGEIIGNIRQAHC | 35 |
| AF153160 | C | CCR5 | CTRPNNNTRKSVRIGPGQTFYATGEVIGNIRQAHC | 35 |
| HQ644817 | B | CCR5 | CIRPNNNTRRSIHIGPGRAFFATGDIIGDIRQAHC | 35 |
| HM215413 | B | CCR5 | CTRPNNNTRKSMTLGPGRAWYTTGQIIGDIRKAHC | 35 |
| DQ061815 | B | CCR5 | CTRPNNNTRKGMHMGPGKVFYATGQIIGDIRQAHC | 35 |
| KC312590 | B | CCR5 | CERPNNNTRKSIRIGPGSAFYATGEIIGNIRQAHC | 35 |
| JQ779233 | C | CCR5 | CIRPGNNTRKSIRIGPGQVFYATNIIGDIREAHC | 34 |
| AY010787 | B | CCR5 | CIRPNNNTRKSIHVGPGKAFYATGDIIGDIRQAYC | 35 |
| DQ061422 | B | CCR5 | CTRPNNNTRKSMHIGPGRAFYTTGEIIGDIRQAHC | 35 |
| AF384308 | B | CCR5 | CTRPNNNTRKSIHIGPGRAFYATGDIIGDIIQAHC | 35 |
| DQ061425 | B | CCR5 | RTRPNNNTRKSIHIGPGRAFYATGEIIGDIKQAHC | 35 |
| DQ235617 | C | CCR5 | CTRPSNNTRKSIRIGPGQAFFATGEIIGDIRQAHC | 35 |
| EU908221 | C | CCR5 | CTRPDNNTRKSIRIGPGQTFYATGDIIGDIRQAHC | 35 |
| JF896837 | B | CCR5 | CTRPNNTRRSINIGPGRAWYTTGEIVGDIRQAHC | 34 |
| DQ388516 | C | CCR5 | CMRPGNNTRRSVRIGPGQTFYATGEIIGDIRQAHC | 35 |
| FJ375985 | C | CCR5 | CTRPGNNTRRSVRIGPGQAFYATGDIIGDPRQAHC | 35 |
| EU908225 | C | CCR5 | CTRPNNNTRRSIRIGPGQTFYATGDIIGNIRQAYC | 35 |
| DQ002337 | B | CCR5 | CTRPSSNTRKSIHLGLGRAFYATGEIIGDIRQAHC | 35 |
| FJ375984 | C | CCR5 | TRPNNNTRKSIRIGPGQAFYATKDIIGDIRKAHC | 34 |
| DQ061819 | B | CCR5 | CTRPNNNTRKGIHMGPGKVFYATGQIIGNIRQAHC | 35 |
| AY510060 | C | CCR5 | CTRPNNNTRRGIRIGPGQTFYATGGIIGDIRQAHC | 35 |
| JX140666 | C | CCR5 | CTRPNNNTRKSIRIGPGQTFYATNDIIGDIREAHC | 35 |
| AF541028 | B | CCR5 | CTRPNNNTRKSIHIGPGRAFYTTGEIIGDIKQAHC | 35 |
| AY159664 | B | CCR5 | CTRPNNNTRKGIHIGPGAALYATGAIIGNIRQAHC | 35 |
| DQ869017 | B | CCR5 | CIRPNNNTRRSIPIGPGRAFYATGDIIGDIRQAHC | 35 |
| AF153140 | C | CCR5 | CTRPGNNTRKSMRIGPGQTFFATGEIIGDIRQAHC | 35 |
| AY010798 | B | CCR5 | CMRPSNNTRKSISIGPGRAFYTTGEIIGDIRQAHC | 35 |
| HM368243 | B | CCR5 | CTRPNNNTRRGIHIGPGGAFYSTGDIIGNIRQAHC | 35 |
| HQ644951 | B | CCR5 | CIRPNNNTRKSIHIGPGRAFYATGEIIGDIRQAYC | 35 |
| EU578340 | B | CCR5 | CTRPNNNTRRSTHLGAGRALYTGEIIGDIRQAHC | 34 |
| EU744122 | B | CCR5 | CTRPNNNTRKSIHIGPGKAFYATGEIIGNIRQAHC | 35 |
| DQ061829 | B | CCR5 | CTRPDNNTRKGIHMGPGKVFYATGQIIGDIRQAHC | 35 |
| EU272264 | B | CCR5 | CTRPNNNTKKSINLGPGRAWYATGQIIGDIRQAHC | 35 |
| EU578358 | B | CCR5 | CTRPNNNTRKSIHIGPGSVFYTGEIIGNIRQAHC | 34 |
| HQ678276 | B | CCR5 | CTRPNNNTRKSIHIGPGRAFYATGEIIGDIRKAHC | 35 |
| AF153177 | C | CCR5 | CTRPNNNTRKSIRIGPGQTFYATNEIIGNIRQAHC | 35 |
| KC312395 | B | CCR5 | TRPNNNTRKSIHIGPGSAFYTTGEIIGDIRQAHC | 34 |
| EU744087 | B | CCR5 | CTRPHNNTRKSIHLGPGRAFYATGDIIGNIRQAYC | 35 |
| DQ235624 | C | CCR5 | CIRPNNNTRKSIRIGPGQTFYATGDIIGDIRKAHC | 35 |
| EU744069 | B | CCR5 | CTRPHNNTRRSIHMGPGKTFYATGDIIGDIRQAHC | 35 |
| HM239574 | B | CCR5 | CTRPSNNTRKSINIGPGRAFYATGDIIGDIRQAHC | 35 |
| HM239584 | B | CCR5 | CTRPSNNTRKDIHIGPGRAFYTTGEIIGDIRKAHC | 35 |
| AF112565 | B | CCR5 | CTRLNNNTRKSIHIGPGRAFYATGDIIGDIRQAHY | 35 |
| JF507871 | B | CCR5 | CTRPGNNTRRSINIGPGRAFYTGEIIGNLRQAHC | 34 |
| HQ708053 | C | CCR5 | CIRPNNNTRKSVRIGPGQTFYATGGIIGDIRRAYC | 35 |
| JF507737 | B | CCR5 | CTRPNNNTRKSIHIGPGRTFFATGEIIGDIRRAHC | 35 |
| EU272288 | B | CCR5 | CTRANNNTRKSIPLGPGKAWYTTGDIIGDIRQAHC | 35 |
| DQ235646 | C | CCR5 | CTRPGNNTRKSIRIGPGQTFYATGDIIGDIRKAYC | 35 |
| HQ644795 | B | CCR5 | CTRPNNNTRRSIHIGPGRAFYTGEIIGDIRKAHC | 34 |
| DQ235626 | C | CCR5 | CTRPNNNTRRSIRIGPGQTFYATGDIIGDIRQAHC | 35 |
| DQ177207 | B | CCR5 | CTRPNNNTRKSITLTPGRAFYATGDIIGDIRQAHC | 35 |
| JF896818 | B | CCR5 | CTRPNNNTRKGIHIGPGRAIYATGAIIGDIRQAHC | 35 |
| HM239607 | B | CCR5 | CRPNNNTRKSIHIAPGRAFYATGDIIGDIQAYC | 33 |
| HQ708061 | C | CCR5 | CTRPNNNTRKSVRIGPGQTFYATGGIIGNIRQAYC | 35 |

| | | | | |
|---|---|---|---|---|
| HQ377465 | B | CCR5 | CTRPNNNTRKSISMGPGRAFYATGEIIGNIRQAHC | 35 |
| DQ061757 | B | CCR5 | CTRPNDNTRRSISLGPGRAYYATGDIIGDIRQAHC | 35 |
| AF194979 | B | CCR5 | CTRPNNNTRRSIHIGPGKAFYTGRNNRNIRQAHC | 34 |
| JF507858 | B | CCR5 | CTRPGNNTGRSINIGPGRAFYTGEIIGDIRQAHC | 34 |
| AF310125 | B | CCR5 | CTRPNNNTRKSIHLGAGRALYTREIIGDIRQAHC | 34 |
| EF688456 | B | CCR5 | CTRPNNNTRKSINMGPGRAFYATGDIIGDIRQAHC | 35 |
| AF153163 | C | CCR5 | CTRPGNNTRRSVRIGPGQAFYATGEIIGNIRRAHC | 35 |
| AY510058 | C | CCR5 | CTRPGNNTRKGIWIGPGQAFYATGDIIGDIRQAHC | 35 |
| DQ869023 | B | CCR5 | CARPNNSTRKGIHIGPGRAFYAAADIIGDIRQAHC | 35 |
| EF117267 | C | CCR5 | CTRPNENRRKSIRIGPGQAFYATGDIIGDIRQARC | 35 |
| AY010881 | B | CCR5 | CTRPNNNTRTGIPMGPGRAMYATGDIIGNIRQAYC | 35 |
| AY669715 | B | CCR5 | CTRPNNNTRKSIHMGGRFYATGEIIGNIRQAHC | 33 |
| GU204932 | B | CCR5 | CTRPNNNTRRSISLGPGRSIYTTGQIIGDIRQAHC | 35 |
| EU577288 | B | CCR5 | CTRPNNNTRRGIHVGPGQALYTGDIIGDIRQAHC | 34 |
| HQ644866 | B | CCR5 | CTRPGNNTRKSIHMGPGRAFYATEDVIGDIRQAHC | 35 |
| EU604557 | B | CCR5 | CTRPSNNTRKSINMGPGRAFYTTGEIIGNIRQAHC | 35 |
| HQ678287 | B | CCR5 | CIRPNNNTRKSINIGPGRAFYATGDIIGDIRRAHC | 35 |
| JF508061 | B | CCR5 | CTRPNNNTRKSIHIGPGRAFYTTGEIIGNIRQAFC | 35 |
| DQ869014 | B | CCR5 | CTRPNNNTRKSIPIGPGRAFFATDIIGDIRQAHC | 34 |
| EU578424 | B | CCR5 | CTRPNNNTRRGVTIGPGRVFYTGQVIGDIRQAHC | 34 |
| KC473835 | B | CCR5 | CTRPNNNTRKGIHIAPGRAFYATGDIIGDIRQAHC | 35 |
| HM239538 | C | CCR5 | CTRPNNNTRESIRIGPGQTFYATGDIIGDIRQAYC | 35 |
| AJ418522 | B | CCR5 | CTRPNNNTRKSIPIGPGGAFYTTGEIIGDIRKAHC | 35 |
| HQ678293 | B | CCR5 | CIRPGNNTRKSIHIAPGRAFYATGDIIGDIRQAHC | 35 |
| FJ375972 | C | CCR5 | CTRNNTRKSRIGPGQTFYATGIIGIRQAHC | 30 |
| AF194978 | B | CCR5 | CTRPNNNTRRSIHIGPGKAFYTGEIIRNIRQAHC | 34 |
| HQ644885 | B | CCR5 | CTRPNNNTRKSINIGPGRAIYATGDIIGNIRQAHC | 35 |
| FJ375994 | C | CCR5 | CTRPGNNTRTSIRIGPGQTFYANNPIIGDIRQAYC | 35 |
| AF307750 | C | CCR5 | CTRPNNNTRKSIRIGPGQVFYATGDIIGDIRQAHC | 35 |
| HQ644895 | B | CCR5 | CTRPNNNTRKSIHIGPGRALYTTGDIIGDIRKAYC | 35 |
| DQ002128 | B | CCR5 | CTRSNNNTRKSIHIEPGRAFYATGDIIGDIRQAHC | 35 |
| U66221 | B | CCR5 | CTRPNNNTRRSVRIGPGGAMFRTGDIIGDIRQAHC | 35 |
| HQ708063 | C | CCR5 | CTRPNNNTRKSMRIGPGQTFYATEEVIGDIRQAYC | 35 |
| AY887884 | C | CCR5 | CTRPGNNTRKGIGIGPGQTFYAPRGIIGDIRQAHC | 35 |
| DQ061495 | B | CCR5 | CTRPNNNTRKSIHIGPGRAFYTTGGIIGKIRQAHC | 35 |
| HQ644909 | B | CCR5 | CTRPNNNTRKSIHIGPGRAFYATGEIIGDIRKAYC | 35 |
| AF384328 | B | CCR5 | CTRPNNNTRKSISIGPGRAFYAHGEIIGDIRQAHF | 35 |
| AF082840 | B | CCR5 | CTRPNNNTRKSIHIGPGKAFYATGDVIGDIRKAHC | 35 |
| DQ002089 | B | CCR5 | CTRPNNNTRKSIHIGPGRAFYATGNMIGNIRQAHC | 35 |
| AY510065 | C | CCR5 | CTRPNNSTRKSVRIGPGQAFYATGDIIGDIRQAHC | 35 |
| JN002026 | B | CCR5 | CTRPNNNTRKSIPMGPGKAFYATGDIIGNIRQAHC | 35 |
| HQ644972 | B | CCR5 | CTRPNNNTRRSIHIGPGRAFYATGDIIGDIRKAHC | 35 |
| AF153182 | C | CCR5 | CTRPGNNTRKSVRIGPGQTFFATGEIIGDIRQAHC | 35 |
| AF384286 | B | CCR5 | CTRPNNNTRKSIHIGPGRAFYARGDIIGDIRQAHC | 35 |
| DQ002315 | B | CCR5 | CTRPNNNTRRSIHMGPGKAFYTTGEIVGDIRQAYC | 35 |
| FJ375991 | C | CCR5 | CTRPNNNTRQSIRIGPGQTYATGDIIGDIRQAHC | 34 |
| AY835441 | B | CCR5 | CTRPNNNTRKGITIGPGRVFYTGEIVGDIRQVHC | 34 |
| DQ061584 | B | CCR5 | CIRPNNNTRKSIRIRPGSAFYTTGEIIGDIRQAHC | 35 |
| HM215403 | C | CCR5 | CTRPSNNTRKSIRIGPGQTFYATGDIIGDIRQAHC | 35 |
| AF384263 | B | CCR5 | CTRPNNNTRRGIHIGPGRAFYATGEIIGDIRQAHC | 35 |
| DQ516161 | B | CCR5 | CTRPNNNTRKDIRIGPGRAFYATGDIIGDIRQAHC | 35 |
| AY713415 | C | CCR5 | CTRYANNTRKSVRIGPGQTFYTNDIIGDIRQAHC | 34 |
| JN687734 | C | CCR5 | CTRPGNNTRKSIRIGPGQAFYATNDIIGDIRQAHC | 35 |
| DQ061482 | B | CCR5 | CTRPNNNTRKSISIGPGRAFYTTGEIIGETRQAYC | 35 |
| KF770339 | C | CCR5 | CIRPGNNTRKSMRIGPGQTFYATGEIIGDIRRAHC | 35 |
| HM239533 | B | CCR5 | CIRPNNNTRKGIHMGMGRAFYATGEVIGNIRQAHC | 35 |
| AF112563 | B | CCR5 | CTRPTNNTRKSIHIAPGSAFYATGDIIGDIRQAHC | 35 |
| DQ002074 | B | CCR5 | CTRPNNSTRKSIHIGPGRAFYATGEIIGNIRQAHC | 35 |
| EF688430 | B | CCR5 | CTRPNNNTRQGINIGPGRAFYTTGEVIGDIRQAHC | 35 |
| EU578323 | B | CCR5 | CTRPNNNTRKSISIGPGRAFYTGEIIGDIRQAHC | 34 |
| JF508118 | B | CCR5 | CTRPNNNTRKGIHIGPGRTFYATGEIIGDMRQAHC | 35 |
| HM239628 | B | CCR5 | CTRPNNTRKSINIGPGRAFYATGDIIGNIRQAHC | 34 |
| JX140653 | B | CCR5 | CIRPNNNTRKSIHMGPGGAFYATGDVIGDIRKAYC | 35 |
| HQ708033 | C | CCR5 | CTRPNNNIRKSVRIGPGQTFYATGDIIGDIRQAYC | 35 |
| JF896821 | B | CCR5 | CTRPNNNTRKSIHLTPGGAYATGDIIGDIRKAC | 33 |
| AY835450 | B | CCR5 | CTRPHNNTRKSIHIGPGRAWYATGDIIGDIRKAYC | 35 |
| AF153168 | C | CCR5 | CPRPINNTRRSVRIGPGQTFYATGEIIGNIRQAHC | 35 |
| FJ376015 | C | CCR5 | CTRPSNNTRESIRIGPGQTFYATGDIIGDIRQAYC | 35 |
| DQ869015 | B | CCR5 | CTRPNNNTRRSIPIGPGRAFYATDIIGDIRQAHC | 34 |
| KF716496 | B | CCR5 | CTRPSNNTRKSINIGPGRAFYTTGEIIGDIRQAHC | 35 |

| | | | | |
|---|---|---|---|---|
| EU578312 | B | CCR5 | CTRPNNNTRKSITIGPGRAFYTGEIIGDIRQAHC | 34 |
| GU204919 | B | CCR5 | CARPNNNTRKSIHIAPGRAFYTTGSIIGDIRQAHC | 35 |
| AY123275 | C | CCR5 | CRPGNNTRESRIGPGQAFFATGVIGDIRKAYC | 32 |
| AY887874 | C | CCR5 | CARPNNNTRKSIRIGPGQTFYATNDIIGNIRQAHC | 35 |
| U04918 | B | CCR5 | CIRPNNNTRRSIHMGPGRAFYATGDIIGDIRQAHC | 35 |
| FJ376024 | C | CCR5 | CTRPNNNTRKSIRIGPGQAFWATGDIIGDIRQAC | 34 |
| JF896872 | B | CCR5 | CTRPNNNTRKSIHGPGSSFETTGEIIGNIRQAHC | 34 |
| EU575201 | B | CCR5 | CTRPSNNTSKSIPIGPGRAFYTTDRIVGDIRQAHC | 35 |
| KC473826 | B | CCR5 | CTRPNNNTRKSIHIAPGKAFYATGDIIGNIRQAHC | 35 |
| HM179801 | C | CCR5 | CIRPNNNTRTSMRIGPGQAFFATNGIIGNIRQAYC | 35 |
| EU744062 | B | CCR5 | CTRPNNNTRKSIHVGPGKTLYATGDIIGDIGQAHC | 35 |
| HM179793 | C | CCR5 | CIRPNNNTRTSIRIGPGQALFATNGIIGNIRQAYC | 35 |
| FJ376017 | C | CCR5 | CIRPGNNTRKSVRIGPGQTFYVNNIIGDIRQAC | 33 |
| DQ061677 | B | CCR5 | CIRPNNNTRKGIHLGPGGALYATGGIIGAIRQAYC | 35 |
| AF153146 | C | CCR5 | CTRPNNNTRKSVRIGPGQTFYATGGIIGDIREAHC | 35 |
| HQ377416 | B | CCR5 | CTRPNNNTRRSIHIGPGSAFYATGEIIGDIRQAHC | 35 |
| EU575795 | B | CCR5 | CTRPGNNTRKSIHIAPGRAFYTTGDIIGDIRQAHC | 35 |
| FJ977090 | C | CCR5 | CTRPNNNTRKSVRIGPGQTFYATGDIIGNTRQAYC | 35 |
| JQ779251 | C | CCR5 | CIRPNNNTRKSIRIGPGQAFFATTGIIGNIRQAHC | 35 |
| EU576666 | B | CCR5 | CTRPNNNTRKGIHIGPGKTLYATGEIIGDIRQAHC | 35 |
| DQ061771 | B | CCR5 | CTRPNDNTRKGIHIGPGRAFYATGEIIGNIRQAHC | 35 |
| AY010858 | B | CCR5 | CTRPNNNTGTCIPMGPGRAVCATGDIIGNIRQAYC | 35 |
| KC473824 | B | CCR5 | CIRPNNNTRKSIHLGPGRAFYATGEIIGDIRKAHC | 35 |
| AY887882 | C | CCR5 | CVRPNNNTRKSMRIGPGQTFYATGEIIGDIRQAHC | 35 |
| JF508043 | B | CCR5 | CTRPNNNTRKSIPIGPGKAFYTTGEIIGDIRQAHC | 35 |
| AY010859 | B | CCR5 | CTRPNNNTGTSIHMGPGRAVYATGDIIGNIRHAYC | 35 |
| EU576296 | B | CCR5 | CIRPNNNTRKGIHIGPGRTFYTTGDIIGDIRQAYC | 35 |
| EU578541 | B | CCR5 | CTRPNNNTRRSIPLGPGAAFFTGEIIGDIRQAYC | 34 |
| DQ061451 | B | CCR5 | RTRPNNNTRKSIHIGPGRAFYTTGEIIGDIRKAYC | 35 |
| AF153159 | C | CCR5 | CTRPNNNTRTSIRIGPGQTFYATGDIIGDIRQAHC | 35 |
| JF896838 | B | CCR5 | CTRPNNNTRKSIPMGPGKAFYATGAIIGIRQAHC | 34 |
| DQ177195 | B | CCR5 | CTRPGNNTRKSIHIGPGRAFYTTGSIIGDIRKAHC | 35 |
| DQ235644 | C | CCR5 | CIRPNNNTRKSIRIGPGQVFYANNDIIGDIRQAHC | 35 |
| DQ516173 | B | CCR5 | CTRPNNNTRKGIHIGPGRAFFTTGAIIGNIRQAHC | 35 |
| HM239573 | B | CCR5 | CTRPNNNTRRSIHLGPGRTIFATGTVIGEIRRAHC | 35 |
| HQ678260 | B | CCR5 | CTRPNNNTRKRITIGPGKVFYATGDIIGDIRQAHC | 35 |
| DQ002183 | B | CCR5 | CARPNNNTRRSITIGPGRAFYATDIIGDIRKAHC | 34 |
| AF254779 | B | CCR5 | CTRPGNNTRKSISMGPGRAFYATGDIIGDIRQAYC | 35 |
| AF153169 | C | CCR5 | CTRPNNNTRKSVRIGPGQTFYATGSIIGDIRQAHC | 35 |
| DQ178989 | B | CCR5 | CTRPNNNTRRSIPIGPGRAFYATGDIIGDIRKAHC | 35 |
| DQ002307 | B | CCR5 | CTRPNSNTRKSINIGPGRAWYTTGDIIGDIRKAHC | 35 |
| AF384256 | B | CCR5 | CTRPSNNTRKGIHIGPGRAWYATGSITGDIRQAHC | 35 |
| JQ779177 | C | CCR5 | CTRPGNNTRKSVRIGPGQTFYATGQIIGNIREAHC | 35 |
| AF384275 | B | CCR5 | CTRPNNNTRKSIHMGPGGSFYATGDIIGNIRQEHC | 35 |
| HM239597 | B | CCR5 | CTRPNNNTRKSINIAPGRAFYATGDVIGDIRQAHC | 35 |
| EU575376 | B | CCR5 | CTRPNNNTRKSITFGPGRAFYTTGDIIGDIRKAYC | 35 |
| FJ670525 | B | CCR5 | CTRPNNNTRQGIHIGPGRAFYATTDIVGNIRKAHC | 35 |
| DQ002186 | B | CCR5 | CTRPGNNTRKSIHIGPGRAFYATDIIGDLRQAYC | 34 |
| AY010850 | B | CCR5 | CTRPNNNTRKSIHLGPGSAIYATGDIIGDIRQAHC | 35 |
| JQ779204 | C | CCR5 | CTRPGNNTRRSVRIGPGQTFYATEEIIGDIREAHC | 35 |
| AY010855 | B | CCR5 | CTRPNNNTRKRIRIGPGRAFYTTGAIIGDIRQAHC | 35 |
| AF153166 | C | CCR5 | CTRPINNTRRSVRIGPGQTFYATGGIIGNIRQAHC | 35 |
| AF384250 | B | CCR5 | CTRPNNNTRKSIHMGPGRAFYTTGNIIGDIRQAHC | 35 |
| HM239620 | B | CCR5 | CTRPNNNTRKSINIGPGRAFFTTGDIIGDIRQAHC | 35 |
| HQ644996 | B | CCR5 | CTRPNNNTRKSIPMGPGRAFYATGDIIGDIRQAHC | 35 |
| AJ418483 | B | CCR5 | CTRPNNNTRKSIHVGPGRALYATGDIIGEIRQAFC | 35 |
| AF021494 | B | CCR5 | CTRPNNNTRRSIHIGPGRAFYTTGRIIGDIRQAYC | 35 |
| HM368255 | B | CCR5 | CTRPNNNTRRGIHIQPGGAFYATDRIIGDIRQAHC | 35 |
| EF688443 | B | CCR5 | CTRPNNNTKRSIYMGPGRTVHTKGRIIGDIRQAHC | 35 |
| AF153137 | C | CCR5 | CIRPNNNTRKSVRIGPGQTFYATEIIGEIRQAHC | 34 |
| AY170663 | C | CCR5 | CTRPSNNTRTSVRIGPGQTFFATNDVIGDIRQAHC | 35 |
| DQ002254 | B | CCR5 | CTRPNNNTRKSIPIGPGRAFYTTGGIIGDIRQAHC | 35 |
| HQ644846 | B | CCR5 | CTRPNNNTRRSINIGPGRAFYTTGDIIGDIRQAYC | 35 |
| EU786674 | B | CCR5 | CTRPGNNTRRSISMGPGRAIYATGEIIGDIRQAYC | 35 |
| FJ376019 | C | CCR5 | CMRPGNNTRKSIIGPGRAFYAGDIIGDIRQAHC | 33 |
| JF508032 | B | CCR5 | CTRPNNNTRKSINIGPGRAFYTTGEIIGNIKQAHC | 35 |
| JN002013 | B | CCR5 | CTRPNNNTRKSIHIGPGRAFYATGEIVGDIREAHC | 35 |
| DQ235635 | C | CCR5 | CIRPGNNTRRSVRIGPGQTFFATGDIIGDIRQAHC | 35 |
| HQ678265 | B | CCR5 | CTRPNNNTRRSIPMGPSKAFYATGDIIGNIRQAHC | 35 |
| EU575148 | B | CCR5 | CERPNNNTRRSIHIGPGRAFYAGEIIGNIRKAYC | 34 |

| | | | | |
|---|---|---|---|---|
| HM239608 | B | CCR5 | CIRPNNNTRKSINIGPGKAFYATGGIIGDIRQAHC | 35 |
| KF384798 | B | CCR5 | CTKHSINKRKRVTIGPGRVYYSTKEIIGDIRKAHC | 35 |
| AF153178 | C | CCR5 | CTRPNNNTRKSVRIGPGQTFYATDIIGDIRQAHC | 34 |
| HM239555 | B | CCR5 | CTRPSNNTRKIPIGPGRAFYATGDIIGNIRQAHC | 34 |
| HQ708023 | C | CCR5 | CIRPNNNTRKSVRIGPGQTFYATGDIIGNIRKAYC | 35 |
| DQ061697 | B | CCR5 | CTRPNNNTRKGIHMGPGKAFYATGDIIGNIRQAHC | 35 |
| AY510061 | C | CCR5 | CTRPNNNTRKSVRIGPGQALYATGGIIGDIRQAHC | 35 |
| AF254772 | C | CCR5 | CTRPNNNTRRSIRIGPGQAFYATGDIIGDIRQAHC | 35 |
| FJ653125 | B | CCR5 | CTKFNNNTRKSIHIGPGRAFYATGDIIGNIRQASC | 35 |
| JF896816 | B | CCR5 | CTRPNNNTRKSIHMGPGRAYATGEIIGIRQAC | 32 |
| EU743997 | B | CCR5 | CTRPNNNTRKSISIGPGRAFYATGEIIGNIRQAHC | 35 |
| DQ516311 | B | CCR5 | CTRPDNNTRKSIHIGPGRAFYTTGGIIGDIRQAHC | 35 |
| FJ375968 | C | CCR5 | CTPNNTRKSIRIGPGQTFYTNIIGDIRKAHC | 31 |
| AF355644 | B | CCR5 | CTRPNNNTRKSINIGPGRAWYATGKIIGNIRQAHC | 35 |
| U08703 | B | CCR5 | CTRPNNNTRKGIHMGWGRAFYATGEIIGNIRQAHC | 35 |
| GU945312 | C | CCR5 | CIRPNNNTRKSIGIGPGQTFYATGDIIGDIRQASC | 35 |
| KF770331 | C | CCR5 | CIRPGNNTRKGMRIGPGQTFYATGEIIGDIRQAHC | 35 |
| HQ678257 | C | CCR5 | CIRPNNNTRRSVRIGPGQTFYATGDIIGDIRKAYC | 35 |
| AF384284 | B | CCR5 | CTRPNSNTRKGIHIGPGRAFYTTGEIVGDIRQAHC | 35 |
| JF507909 | B | CCR5 | CTRPNNNTRQSVHIGPGRALYTTDIIGDIRKAYC | 34 |
| AY736826 | C | CCR5 | CTRPNNNTRKSIRIGPGQTFYATDIIGDIRQAYC | 34 |
| KF716467 | C | CCR5 | CVRPNNNTRKSLRIGPGQTFYATGDIIGDIRQAHC | 35 |
| AY010853 | B | CCR5 | CTRPSNNTRKSIHIGPGRAFYTTGSIIGDIRQAHC | 35 |
| U08704 | B | CCR5 | CTRPNNNTRKSIHMGWGRAFYATGDIIGDIRQAHC | 35 |
| EU576909 | B | CCR5 | CTRPNNNTRKGIHIGPGKVFYTGEIVGDIRQAHC | 34 |
| AY887880 | C | CCR5 | CTRPANNTRKSVRIGPGQTFYATGAIIGDIRQAHC | 35 |
| DQ061536 | B | CCR5 | CTRPNNNTRKSIPIGPGRASYATGDIIGDIRQAHC | 35 |
| EF657904 | B | CCR5 | CIRPNNNTRKSIPMGPGRAFYATGAIIGNIRQAHC | 35 |
| AF384262 | B | CCR5 | CTRPNNNTRKGIHIGPGRAFYATGEIIGDIRQAHF | 35 |
| FJ846652 | C | CXCR4 | CTRPNNNTRKSMRIGIGRGHAFYTTGKVIGNIRQAH | 37 |
| FJ375993 | C | CXCR4 | CARPNYTRQRIGIGRGQALFTARRIIGNIKQAHC | 34 |
| AF021617 | B | CXCR4 | CTRPNNNTRRRIHIGPGRAVYTTGQIIGDIRKAYC | 35 |
| FJ653149 | B | CXCR4 | CKRPNNNARRRIHIGPGRAFYATDIIGNIRQAYC | 34 |
| EU604549 | B | CXCR4 | CTRPSNHTRKRVTLGPSRVYYTTGEITGDIRRAHC | 35 |
| KF770414 | C | CXCR4 | CMRPGNNTRRRVRIGPGQTFYATGNIIGDIRQAHC | 35 |
| DQ382378 | C | CXCR4 | CTRPGKGKRTRVRIGPGRTFYATGAVTGDIRKAHC | 35 |
| FJ846632 | C | CXCR4 | CTRPDNKISMKRIKIGPGRAFVATKGIKGDIRQAYC | 36 |
| AF146728 | B | CXCR4 | CMRPNNNTRKGIYVGPGRHIYATEKIVGDIRQAHC | 35 |
| FJ846648 | C | CXCR4 | CTRPNNNTRKSVRIGIGRGHAFYTTGKVIGNIRQAH | 37 |
| AF259007 | B | CXCR4 | CTRPNKTIRKGLRLGPGRAFYTMGRIEGYIRQAHC | 35 |
| X01762 | B | CXCR4 | CTRPNNNTRKSIRIQRGPGRAFVTIGKIGNMRQAHC | 36 |
| KF770412 | C | CXCR4 | CTRPNINNVRNVRIGPGRALFKTGKMTGDIRQASC | 35 |
| AY842799 | B | CXCR4 | CTRPNNNTRKRITAGPGRVLYTTGQIIGDIRRAHC | 35 |
| FJ798547 | B | CXCR4 | CTRPNNNTRKRITMGPGRVYYTTGQIIGNIRQAHC | 35 |
| FJ798429 | B | CXCR4 | CTRPNNNTRNRISIGPGRAFYTTRQVIGDIRQAHC | 35 |
| JN001990 | B | CXCR4 | CTRPNNNTKKGIYVGPGRKVYTTDRIIGDIRQAHC | 35 |
| KF770420 | C | CXCR4 | CTRPSNNTRRRVRIGRGQAFDATGQIIGDIRQAHC | 35 |
| AF355735 | B | CXCR4 | CTRPGNKTIKSISLGPGRAFSATRQIIGDIRKAYC | 35 |
| AY736819 | B | CXCR4 | CTRPNYNKRKRIHIGPGRAFYTTKNIIGTIRQAHC | 35 |
| FJ541293 | C | CXCR4 | CTRPNNNTRKSVRIGIGRGQAIYAKKAIIGDIRQAH | 37 |
| AY842828 | B | CXCR4 | CARPNNNTRKRIYMGTGRYMSATEKITGDIRQAHC | 35 |
| AF258981 | B | CXCR4 | CTRPNKNIRKGLRLGPGRAFYTMGGIVGYIRQAHC | 35 |
| KF770417 | C | CXCR4 | CVRPNNNTRKSVRIGRGQTFYANRIIGDIRQAHC | 34 |
| FJ846659 | C | CXCR4 | CIRPGNNTRKRVRLGIGPGQTFYATGRVIRDIRQAH | 37 |
| FJ798399 | B | CXCR4 | CIRPGNNTSKRISIGPGRAFRATKIIGDIRKAHC | 34 |
| AF180903 | B | CXCR4 | CTRPNNNTRRRIYIGQGRAVYTTKQIVGDIRKAYC | 35 |
| AF259052 | B | CXCR4 | CTRPGNNTRKRISIGPGRAFYTTEQIIGNIRQAHC | 35 |
| AJ810483 | B | CXCR4 | CTRHHEIIKRRKLHIGPGRPFYTAEIGDRRKAYC | 34 |
| FJ846645 | C | CXCR4 | CTRPNNNTRKSIRTGIGRGQTFYVTGQIIGDVRQAH | 37 |
| FJ798362 | B | CXCR4 | CTRPNNNTRKGIHIGLGRRVYYVTRQIIGDTKRAHC | 35 |
| L31963 | B | CXCR4 | CTRPNNNTRKKFRIQRGPGRAFVTIGKIGNMRQAHC | 36 |
| AF034384 | B | CXCR4 | CTRPNNYKRKRITTGPGRVLYTTGQIIGDIRRAYC | 35 |
| FJ798460 | B | CXCR4 | CTRPNNNTRRGVYIGPGKAFYTTDRIIGDIRQAHC | 35 |
| AF258999 | B | CXCR4 | CTRPNKTTRKGLRLGPGRAFYTLGGIVGYIRQAHC | 35 |
| JN001993 | B | CXCR4 | CTRPNNNTKRGIYVGPGRKVYTTDRIIGDIRQAHC | 35 |
| AY265949 | C | CXCR4 | CGRPNNHRIKGLRIGPGRAFFAMGAIGGGEIRQAHC | 36 |
| U48207 | B | CXCR4 | CTRPNNNTRRSIPIGPGRAFYATGDIIGDIRQAHC | 35 |
| FJ846634 | C | CXCR4 | CTRPNNNTRKSMRIGIGRGQTFYAMGRIIGDIRQAH | 37 |
| AF034378 | B | CXCR4 | CTRPNNYKKKRITIGPGRVLYTTGQIIGDIRRAYC | 35 |
| FJ846655 | C | CXCR4 | CTRPNNNTRKNVRIGIGRGQTFNAMGRIIGNIRQAH | 37 |

| | | | | |
|---|---|---|---|---|
| FJ798540 | B | CXCR4 | CTRPNNNTRGRLSIGPGRAFYATRDIIGDIRRAHC | 35 |
| FJ846640 | C | CXCR4 | CARPGNNTIKRIRIGPRYAFYAKETIIGDIRQAHC | 35 |
| AM156922 | B | CXCR4 | CTRPHNTIKRRIHIGPGRAFYTTKGIQGDLRQAHC | 35 |
| FJ846628 | C | CXCR4 | CTRPDNKINMKRIKIGPGRAFVATKGIRGDIRQAHC | 36 |
| FJ653155 | B | CXCR4 | CKRPNNNARRHIHIGPGRAFYATDIIGNIRQAYC | 34 |
| FJ541294 | C | CXCR4 | CTRPNNNTRKRVRIGHRHLVYAHGEIIGNIRQAHC | 35 |
| AY842819 | B | CXCR4 | CTRPYNLKKSITRGPGRVIYSTGDIMGDIRKAHC | 34 |
| DQ869028 | B | CXCR4 | CTRPGNSTRRGILVGTRRFYTTRNIIGDIRKAHC | 34 |
| EF688436 | B | CXCR4 | CTRPHNKAIRHIHIGQGRAFTTGSIEGNIRQAHC | 34 |
| AF033819 | B | CXCR4 | CTRPNNNTRKRIRIQRGPGRAFVTIGKIGNMRQAHC | 36 |
| AF075721 | B | CXCR4 | CTRPNYNKRKRIHIGPGRAFYTTKNIKGTIRQAHC | 35 |
| DQ904343 | C | CXCR4 | CTRPGNKTIRSIRIGPGRTFYTNKGDIRQAYC | 32 |
| AF021669 | B | CXCR4 | CTRPNNNTRNRIYIGQGRAVYTTKQIVGDIRKAYC | 35 |
| FJ653151 | B | CXCR4 | CKRPNNNMRRHIHIGPGRAFYTTDIIGNMRRAYC | 34 |
| KF770430 | C | CXCR4 | CTRPGNNTGRSVRIGLRTFYTTRKIIGDIRAAHC | 34 |
| U04904 | B | CXCR4 | CTRPNNNTRRSVHSGHIGGGRTLFTTHIVGDIRKAH | 37 |
| FJ846657 | C | CXCR4 | CTRPNNNTRKNVRIGIGRGQTFYANGRIIGNIRQAH | 37 |
| AF258985 | B | CXCR4 | CTRPNKNIRKGLRLGPGRAFYTMGGIVGNIRQTHC | 35 |
| AF258988 | B | CXCR4 | CTRPNKTTRKGLRLGPGRAFYTMGGIVGYIRQAHC | 35 |
| FJ798539 | B | CXCR4 | CTRPNNNTRRRLSIGPGRAFYATRDIIGDIRQAHC | 35 |
| AY736821 | B | CXCR4 | CTRPNNYTRKRITMGPGRVYYTTGEIIGDIRAHC | 34 |
| HQ678267 | B | CXCR4 | CTRPNTKTRKRIHIGPGRAFYTTKTVRDIRQAHC | 34 |
| AF021622 | B | CXCR4 | CTRPNNNTRRRIYIGQGRAVYTTRQIIGDIRKAYC | 35 |
| DQ382362 | C | CXCR4 | CSRPGNNTRKSVRIGIGRGQTFYATGKVIGDIRQAH | 37 |
| FJ798402 | B | CXCR4 | CIRPGNNTSKRVSIGPGRAFRATKVIGDIRKAHC | 34 |
| EU578431 | B | CXCR4 | CTRPNNYTRKHIHLGARKAFYTGEIVGDIRQAHC | 34 |
| AF021647 | B | CXCR4 | CTRPNNNTRKRIYIGQGRAVYTTRQIIGDIRKAYC | 35 |
| FJ846630 | C | CXCR4 | CTRPDNKINMKRIKIGPGRAFVATKGIKGDIRQAYC | 36 |
| GU647196 | B | CXCR4 | CTRPNNNTRKRVSIGPGRAWYTTKQIVGDIRQAHC | 35 |
| DQ286958 | B | CXCR4 | CTRPDNNIKRRIIHIGPGRAFHATKTGDIRQAYC | 34 |
| AF034377 | B | CXCR4 | CTRPNNYKKKRITVGPGRVLYTTGQIIGDIRRAHC | 35 |
| FJ798432 | B | CXCR4 | CTRPNNNTRKRISIGPGRAFYTTRQVIGDIRQAHC | 35 |
| AY173956 | B | CXCR4 | CTRPNNKARRRIRIGPGRTFYTGKIVGDIRQAYC | 34 |
| M14100 | B | CXCR4 | CTRPNNNTRKKIRIQRGPGRAFVTIGKIGNMRQAHC | 36 |
| AY230878 | C | CXCR4 | CMRPGNNTRKRVRIGIGPRQTFYAPGGINKDIRQAH | 37 |
| FJ798323 | B | CXCR4 | CTRPNNNTRKGIHIGLGRRFYVTQIIGDVKRAHC | 34 |
| AY842826 | B | CXCR4 | CARPNNNTRKGIHMGPGRAMYATEKITGDIRQAHC | 35 |
| AF189159 | B | CXCR4 | CTKPNNNTRKRIRIQRGPGRAFVTVGKIGNMRQAHC | 36 |
| AF355744 | B | CXCR4 | CTRPGNKTIRSISLGPGRAFSVTRQIIGDIRKAYC | 35 |
| AF259049 | B | CXCR4 | CTRPGNNTRRRISIGPGRAFYTTEQIIGNIRQAHC | 35 |
| FJ798404 | B | CXCR4 | CIRPGNNTSKRISIGPGRAFRATKVIGDIRKAHC | 34 |
| AY189526 | B | CXCR4 | CTRVSKNIRQKRIKIGPGRAFVATGDIGDIRKAHC | 35 |
| AY842833 | B | CXCR4 | CARPNNNTRKRIYMGTGRYMSATEKITGDIRQARC | 35 |
| FJ798570 | B | CXCR4 | CTRPNNNTRRGIHIGLGRRFYVTQVIGDVKRAHC | 34 |
| FJ375983 | C | CXCR4 | CTRPNRNTKKRITLGPGRVVYTTNEIVGDIRQHC | 34 |
| AF021670 | B | CXCR4 | CTRPNNNTRNRIYIGQGRAVYTTKQIIGDIRKAYC | 35 |
| FJ798576 | B | CXCR4 | CTRPNNNTRKGIHIGPGRAFYATGQIIGDIRQAHC | 35 |
| AF355743 | B | CXCR4 | CTRPGNKTIRSISLGPGRAFSATRQIIGDIRKAYC | 35 |
| AY529679 | C | CXCR4 | CTRPYYNKRRSMRIGIGRGQALYATKEITGDIRRAY | 37 |
| DQ286957 | B | CXCR4 | CQRPNNHTRKRITMSPGRVVYTTGEVIGDIRRAHC | 35 |
| FJ541295 | C | CXCR4 | CGRPNNHRIKGLRIGPGRAFFAMGAIRGGEIRQAHC | 36 |
| FJ541290 | C | CXCR4 | CTRPNNNTRKSVRIGIGRGQTFYATGEIVGDIRQAH | 37 |
| AF021630 | B | CXCR4 | CTRPNNNTRKRIYIGQGRAVYTTKQIVGDIRKAYC | 35 |
| KF384805 | B | CXCR4 | CTRPSNNTRKGIHIGPGRAFFATGDIIGDIRRAHC | 35 |
| AF411966 | C | CXCR4 | CTRPGSNKQRIRNIRIGPGRAFHTNGVIGDIRKAYC | 36 |
| KF770413 | C | CXCR4 | CTRPNINRERNVRIGPGRAFFRTGQMTGDIRQASC | 35 |
| KF384801 | B | CXCR4 | CTRPNNNIRKRIIHIGPGRPFYATGDIGNIRRAQC | 35 |
| AF021639 | B | CXCR4 | CTRPNNNTRKRIYIGQGRAVYTTRQIVGDIRKAYC | 35 |
| AY265948 | C | CXCR4 | CMRPNNNTRKSVRIGPGQTFFATGAIIGNIRQAHC | 35 |
| FJ541296 | C | CXCR4 | CTRPGNKTRQSIRIGIGRGQSFHATGAIIGDIRKAY | 37 |
| KF384799 | B | CXCR4 | CTRHNNNKKIQRIHIGPGRAFVATKGITGDIRQAHC | 36 |
| KF384802 | B | CXCR4 | CTRPNYSTRKSIHIGPGRAFYTTKQIRGNIIQAHC | 35 |
| DQ904342 | C | CXCR4 | CTRPGNKTIRSIRLGPGQAFYTNKGDIRQASC | 32 |
| AM156916 | B | CXCR4 | CTRPNNYTRKGIRIGPGRAVYAAEKIVGNIRQAHC | 35 |
| FJ798430 | B | CXCR4 | CTRPSNNTRRRISIGPGRAFYTTRQVIGDIRQAHC | 35 |
| KF384800 | B | CXCR4 | CTRPYRVITKRIMHIGPGRTFHTTGTIGNIRHAYC | 35 |
| L22956 | C | CXCR4 | CARPGNNTRKSIRIGPGQTFFATGAIIGDIRQAHC | 35 |
| FJ653156 | B | CXCR4 | CKRPNNNMRRHIHIGPGRAFYATDIIGNIRQAYC | 34 |
| EU578395 | B | CXCR4 | CTRPNNNTIKTIRMGIRRAFYTKEIIGDIRQAHC | 34 |
| HM215420 | B | CXCR4 | CTRPNNNTRKRVTLGPGRVWYTTGQIIGDIRKAHC | 35 |

| | | | | |
|---|---|---|---|---|
| KF766540 | C | CXCR4 | CTRPYNNTRKSIGIGPGQAFYATGDIIGDIRQAHC | 35 |
| AF259009 | B | CXCR4 | CTRPNKTIRKGLRLGPGRAFYTMGGIEGYIRQAHC | 35 |
| KF770415 | C | CXCR4 | CIRPGNNTRRRVRIGPGQTFYATGNIIGDIRQAHC | 35 |
| FJ798527 | B | CXCR4 | CTRPNNNTRRRISIGPGRAFTTRDIIGDIRQAHC | 34 |
| FJ798416 | B | CXCR4 | CTRPNNNTRQRISIGPGRAFYTTRQVIGDIRQAHC | 35 |
| AY173951 | B | CXCR4 | CTRPNNYTRKRITMGPGRVYYTTGEIIGDIRRAHC | 35 |
| FJ653150 | B | CXCR4 | CKRPNNNMRRHIHIGPGRAFYTTDIIGNIRRAYC | 34 |
| KF770411 | C | CXCR4 | CTRPNINRERKVRIGPGRAFFRTGQMTGDIRQASC | 35 |
| FJ798426 | B | CXCR4 | CTRPNNNTRQRISIGPGRAFYTTRQVVGDIRQAHC | 35 |
| AF034375 | B | CXCR4 | CTRPNNYKRKRITTGPGKVLYTTGQIIGDIRRAHC | 35 |
| AF035534 | B | CXCR4 | CTRPNNNIRKRIHIGPGRAFYTTRQIIGNIRQAHC | 35 |
| EF688451 | B | CXCR4 | CTRPNNKKIEGIRIGPGSAYFTRQIKEHMRQTHC | 34 |
| JN001995 | B | CXCR4 | CTRPNNNTKRGIYVGPGRKVYTTDRIIGNIRQAHC | 35 |
| FJ798554 | B | CXCR4 | CTRPNNNTMKSITIGPGRAFYTTGQIIGDIRQAHC | 35 |
| FJ375980 | C | CXCR4 | CTRPGNNTRKNVIGGRGQTYAHGIGDIRQAHC | 32 |
| EU604561 | B | CXCR4 | CTRPNNHTRKRVTLGPSRVYYTTGEITGDIRRAHC | 35 |
| AY529678 | C | CXCR4 | CARPGNNTRKMMRIGIGRGQTFYANGQVIGDIRQAH | 37 |
| AF034376 | B | CXCR4 | CTRPNNYKKKRITTGPGRVLYTTGQIIGDIRRAYC | 35 |
| FJ798356 | B | CXCR4 | CTRPNNNTRKGIHIGLGRRFYVTQVIGDVKRAHC | 34 |
| KF384806 | B | CXCR4 | CTRPGKKLSRIIHIGPGRAFYASDGRDIRQAYC | 33 |
| FJ376006 | C | CXCR4 | CRRPGNATRKSVRIGIGRGHTFYATGKIIGDIRKAY | 37 |
| AF034385 | B | CXCR4 | CTRPYNYKKKKITTGPGRVLYTTEEIIGDIRRAHC | 35 |
| DQ990880 | B | CXCR4 | CTRPNNNTRKRITMGPGRVLYTTGQIVGDIRKAHC | 35 |
| DQ177202 | B | CXCR4 | CTRPNNNTRKGIRIGPGRAFIATDKIIGDIRQAHC | 35 |
| EF688433 | B | CXCR4 | CIRPNNNTRRSIHIGPGRAFYATGRVIGDVRRAYC | 35 |
| DQ382372 | C | CXCR4 | CTRPANTRIKRLGIGPGQAFRTVKQIIGDIRQSHC | 35 |

**Supplementary File S2.**

**LDA#1**

First Linear discriminant analysis (LDA#1) on a training dataset of 1838 V3 sequences with known tropism (1701 R5-tropic and 137 X4-tropic) using the following 7 amino acid properties:

KYTJ820101 Hydropathy index (Kyte-Doolittle, 1982)
KRIW790103 Side chain volume (Krigbaum-Komoriya, 1979)
GOLD730102 Residue volume (Goldsack-Chalifoux, 1973)
DAYM780201 Relative mutability (Dayhoff et al., 1978b)
HUTJ700103 Entropy of formation (Hutchens, 1970)
OOBM850102 Optimized propensity to form reverse turn (Oobatake et al., 1985)
ANDN920101 Alpha-CH chemical shifts (Andersen et al., 1992)

Numerical scale of the 7 amino acid properties.

| A.acid | KYTJ820101 | KRIW790103 | GOLD730102 | DAYM780201 | HUTJ700103 | OOBM850102 | ANDN920101 |
|---|---|---|---|---|---|---|---|
| A | 1.8 | 27.5 | 88.3 | 100 | 154.33 | 1.34 | 4.35 |
| R | -4.5 | 105.0 | 181.2 | 65 | 341.01 | 0.95 | 4.38 |
| N | -3.5 | 58.7 | 125.1 | 134 | 207.90 | 2.49 | 4.75 |
| D | -3.5 | 40.0 | 110.8 | 106 | 194.91 | 3.32 | 4.76 |
| C | 2.5 | 44.6 | 112.4 | 20 | 219.79 | 1.07 | 4.65 |
| Q | -3.5 | 80.7 | 148.7 | 93 | 235.51 | 1.49 | 4.37 |
| E | -3.5 | 62.0 | 140.5 | 102 | 223.16 | 2.20 | 4.29 |
| G | -0.4 | 0.0 | 60.0 | 49 | 127.90 | 2.07 | 3.97 |
| H | -3.2 | 79.0 | 152.6 | 66 | 242.54 | 1.27 | 4.63 |
| I | 4.5 | 93.5 | 168.5 | 96 | 233.21 | 0.66 | 3.95 |
| L | 3.8 | 93.5 | 168.5 | 40 | 232.30 | 0.54 | 4.17 |
| K | -3.9 | 100.0 | 175.6 | 56 | 300.46 | 0.61 | 4.36 |
| M | 1.9 | 94.1 | 162.2 | 94 | 202.65 | 0.70 | 4.52 |
| F | 2.8 | 115.5 | 189.0 | 41 | 204.74 | 0.80 | 4.66 |
| P | -1.6 | 41.9 | 122.2 | 56 | 179.93 | 2.12 | 4.44 |
| S | -0.8 | 29.3 | 88.7 | 120 | 174.06 | 0.94 | 4.50 |
| T | -0.7 | 51.3 | 118.2 | 97 | 205.80 | 1.09 | 4.35 |
| W | -0.9 | 145.5 | 227.0 | 18 | 237.01 | -4.65 | 4.70 |
| Y | -1.3 | 117.3 | 193.0 | 41 | 229.15 | -0.17 | 4.60 |
| V | 4.2 | 71.5 | 141.4 | 74 | 207.60 | 1.32 | 3.95 |

Input data: a matrix of R5-tropic sequences (1701 rows and 7 columns) and a matrix of X4-tropic sequences (137 rows and 7 columns). The numerical descriptor of each V3 sequence is a vector of seven components, the first is the arithmetic mean of the hydropathy index and the others are the arithmetic mean of the six indices of physicochemical properties. LDA#1 yields a linear function with the following 7 coefficients:

1) 0.4921 for index KYTJ820101, Hydropathy index
2) 7.6455 for index KRIW790103, Side chain volume
3) -7.3117 for index GOLD730102, Residue volume
4) -0.9267 for index DAYM780201, Relative mutability
5) 1.2554 for index HUTJ700103, Entropy of formation
6) 10.3877 for index OOBM850102, Optimized propensity to form reverse turn
7) -33.3877 for index ANDN920101, Alpha-CH chemical shifts

Mean LDA score = -411.18 (sd = 4.50) in 137 X4-tropic V3 sequences
Mean LDA score = -421.81 (sd = 3.21) in 1701 R5-tropic V3 sequences
Cut-off discriminant value =-417.3824

With a cut-off score of -417.38, 1608 out of 1701 sequences R5-tropic (94.5%) are predicted as R5 (score below the cut-off), and 129 out of 137 sequences X4-tropic (94.2%) as X4 (score above the cut-off). The accuracy of LDA#1 is 94.5%. Accuracy is the number of R5-tropic sequences predicted correctly plus the number of X4-tropic sequences predicted correctly, divided by the total number of sequences and multiplied by 100. The percent frequency distribution of the LDA#1 score is shown in Figure 1A of the text.

**LDA#2**

Second Linear discriminant analysis (LDA#2) on a training dataset of 1838 V3 sequences with known tropism (1701 R5-tropic and 137 X4-tropic) using the following 7 amino acid properties:

KYTJ820101 Hydropathy index (Kyte-Doolittle, 1982)
KRIW790103 Side chain volume (Krigbaum-Komoriya, 1979)
HUTJ700103 Entropy of formation (Hutchens, 1970)
JOND920102 Relative mutability (Jones et al., 1992)
GARJ730101 Partition coefficient (Garel et al., 1973)
FAUJ880104 Length of the side chain (Fauchere et al., 1988)

BIGC670101 Residue volume (Bigelow, 1967)

Numerical scale of the 7 amino acid properties.

| A.acid | KYTJ820101 | KRIW790103 | HUTJ700103 | JOND920102 | GARJ730101 | FAUJ880104 | BIGC670101 |
|---|---|---|---|---|---|---|---|
| A | 1.8 | 27.5 | 154.33 | 100 | 0.28 | 2.87 | 52.6 |
| R | -4.5 | 105.0 | 341.01 | 83 | 0.10 | 7.82 | 109.1 |
| N | -3.5 | 58.7 | 207.90 | 104 | 0.25 | 4.58 | 75.7 |
| D | -3.5 | 40.0 | 194.91 | 86 | 0.21 | 4.74 | 68.4 |
| C | 2.5 | 44.6 | 219.79 | 44 | 0.28 | 4.47 | 68.3 |
| Q | -3.5 | 80.7 | 235.51 | 84 | 0.35 | 6.11 | 89.7 |
| E | -3.5 | 62.0 | 223.16 | 77 | 0.33 | 5.97 | 84.7 |
| G | -0.4 | 0.0 | 127.90 | 50 | 0.17 | 2.06 | 36.3 |
| H | -3.2 | 79.0 | 242.54 | 91 | 0.21 | 5.23 | 91.9 |
| I | 4.5 | 93.5 | 233.21 | 103 | 0.82 | 4.92 | 102.0 |
| L | 3.8 | 93.5 | 232.30 | 54 | 1.00 | 4.92 | 102.0 |
| K | -3.9 | 100.0 | 300.46 | 72 | 0.09 | 6.89 | 105.1 |
| M | 1.9 | 94.1 | 202.65 | 93 | 0.74 | 6.36 | 97.7 |
| F | 2.8 | 115.5 | 204.74 | 51 | 2.18 | 4.62 | 113.9 |
| P | -1.6 | 41.9 | 179.93 | 58 | 0.39 | 4.11 | 73.6 |
| S | -0.8 | 29.3 | 174.06 | 117 | 0.12 | 3.97 | 54.9 |
| T | -0.7 | 51.3 | 205.80 | 107 | 0.21 | 4.11 | 71.2 |
| W | -0.9 | 145.5 | 237.01 | 25 | 5.70 | 7.68 | 135.4 |
| Y | -1.3 | 117.3 | 229.15 | 50 | 1.26 | 4.73 | 116.2 |
| V | 4.2 | 71.5 | 207.60 | 98 | 0.60 | 4.11 | 85.1 |

Input data: a matrix of R5-tropic sequences (1701 rows and 7 columns) and a matrix of X4-tropic sequences (137 rows and 7 columns). The numerical descriptor of each V3 sequence is a vector of seven components, the first is the arithmetic mean of the hydropathy index and the others are the arithmetic mean of the six indices of physicochemical properties. LDA#2 yields a linear function with the following 7 coefficients:
1) 9.8573 for index KYTJ820101, Hydropathy index
2) 6.29021 for index KRIW790103, Side chain volume
3) 1.1429 for index HUTJ700103, Entropy of formation
4) -1.4462 for index JOND920102, Relative mutability
5) -34.5005 for index GARJ730101, Partition coefficient
6) 24.8239 for index FAUJ880104, Length of the side chain
7) -11.0246 for index BIGC670101, Residue volume

Mean LDA score = -251.72 (sd = 4.71) in 137 X4-tropic V3 sequences
Mean LDA score = -262.80 (sd = 3.19) in 1701 R5-tropic V3 sequences
Cut-off discriminant value =-258.3288

With a cut-off score of -258.33, 1603 out of 1701 sequences R5-tropic (94.2%) are predicted as R5 (score below the cut-off), and 129 out of 137 sequences X4-tropic (94.2%) as X4 (score above the cut-off). The accuracy of LDA#2 is 94.2%. Accuracy is the number of R5-tropic sequences predicted correctly plus the number of X4-tropic sequences predicted correctly, divided by the total number of sequences and multiplied by 100. The percent frequency distribution of the LDA#2 score is shown in Figure 1B of the text.

**LDA#3**

Third Linear discriminant analysis (LDA#3) on a training dataset of 1838 V3 sequences with known tropism (1701 R5-tropic and 137 X4-tropic) using the following 7 amino acid properties:
KYTJ820101 Hydropathy index (Kyte-Doolittle, 1982)
KRIW790103 Side chain volume (Krigbaum-Komoriya, 1979)
HUTJ700103 Entropy of formation (Hutchens, 1970)
DAYM780201 Relative mutability (Dayhoff et al., 1978b)
BIGC670101 Residue volume (Bigelow, 1967),
CHAM830105 Number of atoms in the side chain labelled 3+1 (Charton, 1983)
OOBM770102 Short and medium range non-bonded energy per atom (Oobatake-Ooi, 1977)

Numerical scale of the 7 amino acid properties.

| A.acid | KYTJ820101 | KRIW790103 | HUTJ700103 | DAYM780201 | BIGC670101 | CHAM830105 | OOBM770102 |
|---|---|---|---|---|---|---|---|
| A | 1.8 | 27.5 | 154.33 | 100 | 52.6 | 0.0 | -1.40 |
| R | -4.5 | 105.0 | 341.01 | 65 | 109.1 | 1.0 | -0.92 |
| N | -3.5 | 58.7 | 207.90 | 134 | 75.7 | 0.0 | -1.18 |
| D | -3.5 | 40.0 | 194.91 | 106 | 68.4 | 0.0 | -1.16 |
| C | 2.5 | 44.6 | 219.79 | 20 | 68.3 | 0.0 | -1.37 |
| Q | -3.5 | 80.7 | 235.51 | 93 | 89.7 | 1.0 | -1.12 |
| E | -3.5 | 62.0 | 223.16 | 102 | 84.7 | 1.0 | -1.16 |

| | | | | | | | |
|---|---|---|---|---|---|---|---|
| G | -0.4 | 0.0 | 127.90 | 49 | 36.3 | 0.0 | -1.36 |
| H | -3.2 | 79.0 | 242.54 | 66 | 91.9 | 1.0 | -1.22 |
| I | 4.5 | 93.5 | 233.21 | 96 | 102.0 | 0.0 | -1.19 |
| L | 3.8 | 93.5 | 232.30 | 40 | 102.0 | 0.0 | -1.32 |
| K | -3.9 | 100.0 | 300.46 | 56 | 105.1 | 1.0 | -1.07 |
| M | 1.9 | 94.1 | 202.65 | 94 | 97.7 | 1.0 | -1.30 |
| F | 2.8 | 115.5 | 204.74 | 41 | 113.9 | 1.0 | -1.14 |
| P | -1.6 | 41.9 | 179.93 | 56 | 73.6 | 0.0 | -1.24 |
| S | -0.8 | 29.3 | 174.06 | 120 | 54.9 | 0.0 | -1.30 |
| T | -0.7 | 51.3 | 205.80 | 97 | 71.2 | 0.0 | -1.25 |
| W | -0.9 | 145.5 | 237.01 | 18 | 135.4 | 1.5 | -1.03 |
| Y | -1.3 | 117.3 | 229.15 | 41 | 116.2 | 1.0 | -1.03 |
| V | 4.2 | 71.5 | 207.60 | 74 | 85.1 | 0.0 | -1.25 |

Input data: a matrix of R5-tropic sequences (1701 rows and 7 columns) and a matrix of X4-tropic sequences (137 rows and 7 columns). The numerical descriptor of each V3 sequence is a vector of seven components, the first is the arithmetic mean of the hydropathy index and the others are the arithmetic mean of the six indices of physicochemical properties. LDA#3 yields a linear function with the following 7 coefficients:
1) 4.6184 for index KYTJ820101, Hydropathy index
2) 4.9509 for index KRIW790103, Side chain volume
3) 1.3178 for index HUTJ700103, Entropy of formation
4) -0.7525 for index DAYM780201, Relative mutability
5) -8.9141 for index BIGC670101, Residue volume
6) 9.6644 for index CHAM830105, Number of atoms in the side chain labelled 3+1
7) 43.3139 for index OOBM770102, Short and medium range non-bonded energy per atom

Mean LDA score = -218.10 (sd = 4.32) in 137 X4-tropic V3 sequences
Mean LDA score = -228.22 (sd = 3.07) in 1701 R5-tropic V3 sequences
Cut-off discriminant value =-224.0124

With a cut-off score of -224.01, 1610 out of 1701 sequences R5-tropic (94.7%) are predicted as R5 (score below the cut-off), and 129 out of 137 sequences X4-tropic (94.2%) as X4 (score above the cut-off). The accuracy of LDA#3 is 94.6%. Accuracy is the number of R5-tropic sequences predicted correctly plus the number of X4-tropic sequences predicted correctly, divided by the total number of sequences and multiplied by 100. The percent frequency distribution of the LDA#3 score is shown in Figure 1C of the text.
